# Resolving the dark matter of *ABCA4* for 1,054 Stargardt disease probands through integrated genomics and transcriptomics

**DOI:** 10.1101/817767

**Authors:** Mubeen Khan, Stéphanie S. Cornelis, Marta del Pozo-Valero, Laura Whelan, Esmee H. Runhart, Ketan Mishra, Femke Bults, Yahya AlSwaiti, Alaa AlTabishi, Elfride De Baere, Sandro Banfi, Eyal Banin, Miriam Bauwens, Tamar Ben-Yosef, Camiel J.F. Boon, L. Ingeborgh van den Born, Sabine Defoort, Aurore Devos, Adrian Dockery, Lubica Dudakova, Ana Fakin, G. Jane Farrar, Juliana Maria Ferraz Sallum, Kaoru Fujinami, Christian Gilissen, Damjan Glavač, Michael B. Gorin, Jacquie Greenberg, Takaaki Hayashi, Ymkje Hettinga, Alexander Hoischen, Carel B. Hoyng, Karsten Hufendiek, Herbert Jägle, Smaragda Kamakari, Marianthi Karali, Ulrich Kellner, Caroline C.W. Klaver, Bohdan Kousal, Tina Lamey, Ian M. MacDonald, Anna Matynia, Terri McLaren, Marcela D. Mena, Isabelle Meunier, Rianne Miller, Hadas Newman, Buhle Ntozini, Monika Oldak, Marc Pieterse, Osvaldo L. Podhajcer, Bernard Puech, Raj Ramesar, Klaus Rüther, Manar Salameh, Mariana Vallim Salles, Dror Sharon, Francesca Simonelli, Georg Spital, Marloes Steehouwer, Jacek P. Szaflik, Jennifer A. Thompson, Caroline Thuillier, Anna M. Tracewska, Martine van Zweeden, Andrea L. Vincent, Xavier Zanlonghi, Petra Liskova, Heidi Stöhr, John De Roach, Carmen Ayuso, Lisa Roberts, Bernard H.F. Weber, Claire-Marie Dhaenens, Frans P.M. Cremers

## Abstract

Missing heritability in human diseases represents a major challenge. Although whole-genome sequencing enables the analysis of coding and non-coding sequences, substantial costs and data storage requirements hamper its large-scale use to (re)sequence genes in genetically unsolved cases. The *ABCA4* gene implicated in Stargardt disease (STGD1) has been studied extensively for 22 years, but thousands of cases remained unsolved. Therefore, single molecule molecular inversion probes were designed that enabled an automated and cost-effective sequence analysis of the complete 128-kb *ABCA4* gene. Analysis of 1,054 unsolved STGD and STGD-like probands resulted in bi-allelic variations in 448 probands. Twenty-seven different causal deep-intronic variants were identified in 117 alleles. Based on *in vitro* splice assays, the 13 novel causal deep-intronic variants were found to result in pseudo-exon (PE) insertions (n=10) or exon elongations (n=3). Intriguingly, intron 13 variants c.1938-621G>A and c.1938-514G>A resulted in dual PE insertions consisting of the same upstream, but different downstream PEs. The intron 44 variant c.6148-84A>T resulted in two PE insertions that were accompanied by flanking exon deletions. Structural variant analysis revealed 11 distinct deletions, two of which contained small inverted segments. Uniparental isodisomy of chromosome 1 was identified in one proband. Integrated complete gene sequencing combined with transcript analysis, identified pathogenic deep-intronic and structural variants in 26% of bi-allelic cases not solved previously by sequencing of coding regions. This strategy serves as a model study that can be applied to other inherited diseases in which only one or a few genes are involved in the majority of cases.

## Introduction

High throughput whole-genome sequencing (WGS) has made a huge impact in biology and is considered the most powerful genetic test to elucidate inherited human diseases (Lupski et al. 2010; Carss et al. 2017). It allows the unbiased detection of a wide spectrum of genetic variants including coding and non-coding single nucleotide variants (SNVs), as well as large structural variants (SVs) (Ellingford et al. 2016). However, sequencing and data storage costs hamper the use of WGS for routine diagnostics. As WGS can result in ‘incidental genetic findings’, a separate informed consent of the patients is required (Matthijs et al. 2016).

Based on the advantages and limitations mentioned above, WGS is not the best method to perform sequence analysis of one or a few genes that are associated with a clinically distinct condition. Autosomal recessive Stargardt disease (STGD1) due to variants in the *ABCA4* gene, represents a nice example of this concept. STGD1 is the most frequent inherited macular dystrophy with an estimated prevalence of 1/10,000 (Blacharski et al. 1988). Different combinations of variants result in a spectrum of clinical phenotypes, ranging from i) pan-retinal dystrophy with an early age at onset due to two severe or null variants, to ii) adolescent or young adult-onset disease observed in probands with typical STGD1 due to the combined effect of a severe and a mild variant, and iii) late-onset STGD1 often due to a severe variant combined with a mild/hypomorphic variant, one of which (p.Asn1868Ile) displays reduced penetrance (Allikmets et al. 1997; Cremers et al. 1998; Westeneng-van Haaften et al. 2012; Zernant et al. 2017; Cremers et al. 2018; Runhart et al. 2018). Thus far, 1,180 unique *ABCA4* variants have been reported in 8,777 alleles of 6,684 cases (Cornelis et al. 2017) (www.lovd.nl/ABCA4). A large proportion of the variants affect non-canonical splice site (NCSS) sequences, with variable effects on mRNA processing (Schulz et al. 2017; Sangermano et al. 2018; Khan et al. 2019). In addition, several deep-intronic (DI) variants have been identified (Braun et al. 2013; Zernant et al. 2014; Bauwens et al. 2015; Bax et al. 2015; Bauwens et al. 2019; Fadaie et al. 2019; Khan et al. 2019; Sangermano et al. 2019). Most of these DI variants strengthen cryptic splice sites resulting in the insertion of pseudo-exons (PEs) in the mature *ABCA4* mRNA. A few DI variants display retina-specific PE insertions (Albert et al. 2018). SVs seem to be rare in *ABCA4* (Maugeri et al. 1999; Yatsenko et al. 2003; Zernant et al. 2014; Bax et al. 2015; Bauwens et al. 2019), although systematic copy number variant analyses have not been performed in most STGD1 cases.

Due to the relatively large size of the *ABCA4* gene (50 exons; 128,313 bp), mutation screening initially was restricted to mutation-scanning of the exons and flanking splice sites, by methods such as single strand conformation polymorphism analysis (Maugeri et al. 1999; Klevering et al. 2004), denaturing gradient gel electrophoresis (Rivera et al. 2000), denaturing high pressure liquid chromatography (Maia-Lopes et al. 2009), and allele-specific primer extension (Jaakson et al. 2003). More recently, exons and splice sites have been sequenced using an amplicon tagging protocol (Zernant et al. 2011; Sciezynska et al. 2016), array-based hybridization (Schulz et al. 2017), targeted gene-panel sequencing (Consugar et al. 2015) or whole exome sequencing (Ortube et al. 2014; Zhou et al. 2014; Bryant et al. 2018). Sequence analysis of the entire 128-kb gene and up- and downstream DNA segments was performed using next-generation sequencing platforms after enrichment of *ABCA4* sequences using Raindance microdroplet-PCR target enrichment or Illumina TruSeq Custom Amplicon target enrichment (Zernant et al. 2014), Haloplex-based sequence enrichment (Bauwens et al. 2019; Sangermano et al. 2019), or WGS (Carss et al. 2017; Sangermano et al. 2019). The scanning techniques showed poor sensitivity, leaving 50-70% of STGD1 probands genetically unsolved (Maugeri et al. 1999; Rivera et al. 2000; Jaakson et al. 2003; Klevering et al. 2004; Maia-Lopes et al. 2009). Targeted next-generation sequencing methods thus far employed have been relatively expensive to analyze *ABCA4*, or (e.g. whole exome sequencing) did not cover non-coding regions and thus missed crucial causal variants.

Identification of all variants in both alleles is important to confirm the clinical diagnosis because several promising clinical trials are underway based on RNA modulation with antisense oligonucleotides (Albert et al. 2018; Bauwens et al. 2019; Sangermano et al. 2019), drug based therapies, such as compounds that inhibit lipofuscin accumulation (Charbel Issa et al. 2015), as well as gene augmentation (Allocca et al. 2008) and stem cell therapies (Lu et al. 2009). STGD1 cases will only be eligible for one of these therapies if both causal alleles are known. In addition, recent studies have identified alleles carrying a coding variant in *cis* with a deep-intronic variant, and only these combinations represented fully penetrant alleles (Bauwens et al. 2019; Sangermano et al. 2019).

Recently, we reported the design and use of 483 single-molecule molecular inversion probes (smMIPs) to sequence the 50 coding exons and 12 intronic regions carrying 11 pathogenic DI variants of 412 genetically unsolved STGD1 cases (Khan et al. 2019). In this study, we aimed to design a fully automated, high throughput, cost-effective and comprehensive sequence analysis of the entire *ABCA4* gene which could serve as a model study to investigate human inherited diseases due to variants in one or a few genes. Using 3,866 smMIPs we sequenced 1,054 genetically unsolved STGD or STGD-like probands, 123 bi-allelic control cases carrying *ABCA4* single nucleotide variants (SNVs), and 15 affected individuals carrying known heterozygous SVs in *ABCA4*. Novel NCSS and DI variants were tested *in vitro* for splice defects. Additionally, a very high and reproducible read coverage allowed us to perform copy number variant analysis.

## Results

### smMIPs performance and *ABCA4* sequencing

A pilot sequencing study was conducted using 15 STGD1 samples and five controls, revealing all previously identified 34 variants (Supplemental Table S1). The average number of reads for the 20 DNA samples ranged from 10 to 152,500 per smMIP, with an overall average coverage of 933× for each smMIP.

In total 1,192 DNA samples were analyzed for variants in *ABCA4* using six NextSeq500 runs. The average number of reads of the 3,866 smMIPs was 377×. As most nucleotide positions are targeted with two smMIPs, the effective average coverage was ~700×. To determine the coverage of *ABCA4* in more detail, we calculated the average coverage of each nucleotide position for runs 1 to 5 combined (Supplemental Table S2). To visualize the results, nucleotide positions that were not or poorly covered (≤10 reads), moderately covered (11-49 reads) or well covered (≥50 reads) are depicted in Supplemental Fig. S1. From the 128,366 nt of *ABCA4*, 1,980 nt (1.5%) were not or poorly covered, 1,410 nt (1.1%) were moderately covered, and 124,976 nt (97.4%) were well covered. Although *ABCA4* introns carry several repetitive elements (Supplemental Fig. S1), they only had a small effect on smMIPs design. Several larger repeats are present in up- and downstream regions of *ABCA4*, which resulted in the absence - or poor performance - of smMIPs. Sequencing of 1,192 samples yielded a total of 7,756 unique variants in *ABCA4* that are listed in Supplemental Table S3.

### Sensitivity and specificity of the smMIPs-based sequencing

To assess the sensitivity of the new smMIPs sequencing platform, we tested 123 previously genotyped samples (Khan et al. 2019; Sangermano et al. 2019) in three series (runs 2, 3 and 6) (Supplemental Table S4) as well as 15 control DNA samples carrying 13 different SVs spread throughout the *ABCA4* gene (run 6) (Supplemental Table S5). All previously known SNVs (n=300) could be identified, yielding a sensitivity of 100%. However, six additional variants were found due to low coverage in the previous studies, while three variants had not been annotated correctly previously.

### *ABCA4* gene sequencing and identification of variants

*ABCA4* sequencing was performed for 1,054 genetically unsolved STGD and STGD-like patients. This revealed 323 unique (likely) pathogenic SNVs and 11 SVs in 1,144 alleles (Supplemental Table S6). Sixty-four of 323 SNVs (26%) and all 11 SVs were novel. Details are provided in Table 1A-C. Thirteen percent of these alleles were represented by DI variants and SVs and another 10% accounted for NCSS variants (Fig. 1A). All variants and the respective cases were uploaded into the *ABCA4* variant and STGD1 cases database LOVD, at www.lovd.nl/ABCA4.

**Table 1:** Novel rare variants in *ABCA4* and their functional classification. **A.** Novel *ABCA4* coding and canonical splice site variants identified in this study, including genomic positions (human genome version 19; hg19) and predicted effects at the protein level. SIFT scores range from 0 to 1, amino acid substitutions are predicted damaging if the score is ≤ 0.05 and tolerated if the score is > 0.05. Mutation Taster is based on a Bayes classifier that calculates probabilities for the variant to be either a disease mutation or a polymorphism. PolyPhen-2 scores range between 0.000 to 1.000 (scores close to 1 indicate damaging effect of variants). All the variants were classified according to the ACMG guidelines from Class 1-5 (1, benign; 2, likely benign; 3, variant of unknown significance; 4, likely pathogenic; 5, pathogenic). AF, allele frequency; nFE, non-Finnish European; n.a., not applicable; SIFT, Sorting Intolerant From Tolerant; ACMG, American College of Medical Genetics and Genomics; Prob. dam, probably damaging; Pos. dam, possibly damaging. **B.** Novel causal non-canonical splice site variants. For the novel non-canonical splice site variants identified in this study, data are provided regarding their cDNA positions (hg19), RNA and protein effects. Consequences of the splice defects on exons are indicated. Pathogenicity of the variants was assigned according to the percentage of wild-type mRNA obtained by semi-quantification of both wild-type and mutant transcripts. WT, wild-type; # variant already reported, but not tested by Zernant et al, 2014. **C.** Novel causal deep-intronic *ABCA4* variants. For the novel deep-intronic variants identified in this study, data are provided regarding their cDNA positions (hg19), RNA and protein effects. Pathogenicity of the variants was assigned according to the percentage of wild-type mRNA obtained by semi-quantification of both wild-type and mutant transcripts. Variants with asterisk do not create a pseudo-exon but led to exon elongations. WT, wild-type; # variant already reported, but not tested by Zernant et al, 2014; $ variants for which Sanger sequencing is not available

| Genomic position<br>(hg 19) | DNA variant | Protein variant | AF in study | gnomAD_AF<br>(nFE) | *In silico* analysis |  |  | ACMG<br>class |
| --- | --- | --- | --- | --- | --- | --- | --- | --- |
|  |  |  |  |  | SIFT<br>(scores) | Mutation<br>Taster | PolyPhen-2<br>(scores) |  |
| 94586555_94586556 | c.46_47del | p.(Thr16Profs*37) | 0.0005 | - | n.a. | n.a. | n.a. | 5 |
| 94578597 | c.92G>A | p.(Trp31*) | 0.0005 | - | n.a. | n.a. | n.a. | 5 |
| 94577117 | c.179C>G | p.(Ala60Gly) | 0.0009 | - | Deleterious (0) | Disease causing | Prob. dam. (0.993) | 4 |
| 94564529 | c.589G>C | p.(Asp197His) | 0.0005 | 0.00007954 | Deleterious (0) | Disease causing | Pos. dam. (0.943) | 3 |
| 94564366 | c.752del | p.(Phe251Serfs*11) | 0.0005 | - | n.a. | n.a. | n.a. | 5 |
| 94548932 | c.834del | p.(Asp279Ilefs*21) | 0.0005 | - | n.a. | n.a. | n.a. | 5 |
| 94546264 | c.869G>A | p.(Arg290Gln) | 0.0005 | - | Deleterious (0.05) | Polymorphism | Benign (0.020) | 3 |
| 94546117 | c.1016G>T | p.(Trp339Leu) | 0.0005 | - | Deleterious (0) | Disease causing | Benign (0.205) | 3 |
| 94544936 | c.1181T>A | p.(Leu394Gln) | 0.0005 | - | Deleterious (0.01) | Disease causing | Pos. dam. (0.904) | 3 |
| 94544209 | c.1293G>C | p.(Trp431Cys) | 0.0005 | - | Deleterious (0) | Disease causing | Prob. dam. (1.000) | 3 |
| 94543441 | c.1359T>G | p.(Asp453Glu) | 0.0005 | - | Deleterious (0) | Disease causing | Benign (0.049) | 3 |
| 94543277 | c.1667T>G | p.(Met556Arg) | 0.0005 | - | Deleterious (0) | Disease causing | Benign (0.033) | 3 |
| 94528751 | c.1677G>A | p.(Trp559*) | 0.0005 | - | n.a. | n.a. | n.a. | 5 |
| 94528698 | c.1730T>A | p.(Val577Glu) | 0.0005 | - | Deleterious (0.02) | Disease causing | Benign (0.034) | 3 |
| 94528304 | c.1766G>C | p.(Trp589Ser) | 0.0005 | - | Deleterious (0) | Disease causing | Prob. dam. (0.990) | 3 |
| 94528265 | c.1805G>T | p.(Arg602Leu) | 0.0005 | - | Deleterious (0) | Disease causing | Benign (0.288) | 3 |
| 94526295 | c.1958G>T | p.(Arg653Leu) | 0.0005 | - | Deleterious (0) | Disease causing | Benign (0.057) | 3 |
| 94522302 | c.2237T>A | p.(Met746Lys) | 0.0005 | - | Deleterious (0) | Disease causing | Pos. dam. (0.476) | 3 |
| 94522296 | c.2243delinsTT | p.(Cys748Phefs*18) | 0.0005 | - | n.a. | n.a. | n.a. | 5 |
| 94522192 | c.2347C>T | p.(Gln783*) | 0.0005 | - | n.a. | n.a. | n.a. | 5 |
| 94520773 | c.2481del | p.(Thr829Argfs*14) | 0.0005 | - | n.a. | n.a. | n.a. | 5 |
| 94520771 | c.2483C>T | p.(Pro828Leu) | 0.0005 | - | Deleterious (0) | Disease causing | Benign (0.025) | 3 |
| 94512619 | c.2774G>A | p.(Trp925*) | 0.0005 | - | n.a. | n.a. | n.a. | 5 |
| 94510227 | c.2992dup | p.(Leu998Profs*25) | 0.0005 | - | n.a. | n.a. | n.a. | 5 |
| 94509011 | c.3071T>A | p.(Met1024Lys) | 0.0005 | - | Deleterious (0.01) | Disease causing | Benign (0.175) | 4 |
| 94508993 | c.3089T>A | p.(Leu1030Gln) | 0.0005 | - | Deleterious (0) | Disease causing | Prob. dam. (1.000) | 4 |
| 94508454_94508456 | c.3191-2_3191del | p.(?) | 0.0009 | - | n.a. | n.a. | n.a. | 5 |
| 94508330 | c.3315del | p.(Lys1106Serfs*42) | 0.0009 | - | n.a. | n.a. | n.a. | 5 |
| 94506959 | c.3329-1G>A | p.(?) | 0.0005 | 0.000007955 | n.a. | n.a. | n.a. | 5 |
| 94506943 | c.3344T>C | p.(Met1115Thr) | 0.0005 | 0.000003977 | Deleterious (0.01) | Disease causing | Benign (0.062) | 4 |
| 94502847 | c.3667dup | p.(Glu1223Glyfs*14) | 0.0005 | - | n.a. | n.a. | n.a. | 5 |
| 94502832 | c.3682G>A | p.(Glu1228Lys) | 0.0005 | - | Deleterious (0) | Disease causing | Benign (0.356) | 3 |
| 94502720 | c.3794C>G | p.(Ser1265Cys) | 0.0005 | - | Deleterious (0) | Disease causing | Prob. dam. (0.997) | 3 |

| Genomic position<br>(hg 19) | DNA variant | Protein variant | AF in study | gnomAD_AF<br>(nFE) | In silico analysis |  |  | ACMG<br>class |
| --- | --- | --- | --- | --- | --- | --- | --- | --- |
|  |  |  |  |  | SIFT<br>(scores) | Mutation<br>Taster | PolyPhen-2<br>(scores) |  |
| 94495000 | c.4539+1G>C | p.(?) | 0.0005 | - | n.a. | n.a. | n.a. | 5 |
| 94490511 | c.4633A>G | p.(Ser1545Gly) | 0.0005 | - | Deleterious (0) | Disease causing | Benign (0.002) | 3 |
| 94490509 | c.4634+1del | p.(?) | 0.0019 | - | n.a. | n.a. | n.a. | 5 |
| 94490508 | c.4634+2T>C | p.(?) | 0.0005 | - | n.a. | n.a. | n.a. | 5 |
| 94487488 | c.4687G>A | p.(Gly1563Arg) | 0.0005 | - | Deleterious (0) | Disease causing | Prob.dam. (0.998) | 3 |
| 94487469 | c.4706del | p.(Val1569Alafs*12) | 0.0005 | - | n.a. | n.a. | n.a. | 5 |
| 94487463 | c.4712T>A | p.(Ile1571Asn) | 0.0005 | 0.000007953 | Deleterious (0.01) | Disease causing | Benign (0.004) | 3 |
| 94487436_94487439 | c.4736_4739del | p.(Phe1579*) | 0.0009 | - | n.a. | n.a. | n.a. | 5 |
| 94487235 | c.4809del | p.(Asp1604Ilefs*6) | 0.0005 | - | n.a. | n.a. | n.a. | 5 |
| 94487213 | c.4831del | p.(Thr1611Leufs*51) | 0.0005 | - | n.a. | n.a. | n.a. | 5 |
| 94487195 | c.4848+1G>T | p.(?) | 0.0005 | - | n.a. | n.a. | n.a. | 5 |
| 94486850 | c.4964C>T | p.(Thr1655Ile) | 0.0005 | - | Deleterious (0) | Disease causing | Prob.dam. (0.999) | 3 |
| 94485185 | c.5149G>T | p.(Gly1717*) | 0.0005 | - | n.a. | n.a. | n.a. | 5 |
| 94481372 | c.5235C>G | p.(Ile1745Met) | 0.0005 | 0.00001195 | Deleterious (0) | Disease causing | Prob.dam. (1.000) | 3 |
| 94480248 | c.5313-2del | p.(?) | 0.0005 | - | n.a. | n.a. | n.a. | 5 |
| 94480208_94480209 | c.5350_5351insGA | p.(Leu1784Argfs*47) | 0.0005 | - | n.a. | n.a. | n.a. | 5 |
| 94480155 | c.5404A>G | p.(Ile1802Val) | 0.0005 | - | Deleterious (0) | Disease causing | Benign (0.172) | 3 |
| 94476893 | c.5509C>T | p.(Pro1837Ser) | 0.0005 | - | Deleterious (0) | Disease causing | Prob.dam. (1.000) | 4 |
| 94476830 | c.5572T>A | p.(Tyr1858Asn) | 0.0005 | 0.000007958 | Deleterious (0) | Disease causing | Prob.dam. (0.957) | 3 |
| 94476430 | c.5640T>A | p.(Phe1880Leu) | 0.0005 | 0.0006082 | Tolerated (0.58) | Polymorphism | Benign (0.008) | 3 |
| 94471062 | c.6082A>G | p.(Thr2028Ala) | 0.0009 | - | Deleterious (0) | Disease causing | Prob.dam. (1.000) | 4 |
| 94467507 | c.6189C>A | p.(Tyr2063*) | 0.0005 | - | n.a. | n.a. | n.a. | 5 |
| 94467480 | c.6216T>G | (Ser2072Arg) | 0.0005 | - | Deleterious (0) | Disease causing | Prob.dam. (1.000) | 4 |
| 94467442 | c.6254T>C | p.(Leu2085Pro) | 0.0005 | - | Deleterious (0) | Disease causing | Prob.dam. (0.998) | 4 |
| 94466634_94466635 | c.6309_6310del | p.(Gln2104Glyfs*30) | 0.0005 | - | n.a. | n.a. | n.a. | 5 |
| 94466564 | c.6380C>T | p.(Ser2127Phe) | 0.0005 | - | Deleterious (0) | Disease causing | Prob.dam. (0.998) | 4 |
| 94466474 | c.6397T>C | p.(Cys2133Arg) | 0.0005 | - | Deleterious (0) | Disease causing | Prob.dam. (1.000) | 4 |
| 94466399 | c.6472A>G | p.(Lys2158Glu) | 0.0005 | - | Deleterious (0) | Disease causing | Prob.dam. (0.998) | 4 |
| 94463529 | c.6617T>A | p.(Leu2206His) | 0.0005 | - | Deleterious (0) | Disease causing | Prob.dam. (1.000) | 3 |
| 94461693 | c.6788G>C | p.(Arg2263Pro) | 0.0005 | 0.000003981 | Deleterious (0) | Disease causing | Prob.dam. (0.998) | 3 |
| 94461663 | c.6816+2T>A | p.(?) | 0.0005 | - | n.a. | n.a. | n.a. | 5 |

**B. Novel causal non-canonical splice site variants in *ABCA4***
| DNA variant | RNA effect | Protein effect | Splice defect | % WT mRNA | Proposed severity |
| --- | --- | --- | --- | --- | --- |
| c.1554+3A>T | r.[1357_1554del,=] | p.[Asp453_Glu518del,=] | Exon 11 skipping | 51 | Moderate |
| c.2654-8T>G<sup>#</sup> | r.[2653_2654ins2654-40_2654-1,=] | p.[Gly863Valfs*47,=] | Exon 18 elongation | 13 | Severe |
| c.3862G>A | r.[3863g>a,3814_3862del] | p.[Gly1288Ser,Ile1272Valfs*101] | Exon 26 skipping | 69 | mild |
| c.4129-3C>T | r.[=,3864_4128del,4129_4253del,3864_4253del] | p.[=,Ile1377Hisfs*3,Gly1288Aspfs*45] | Exon 28, 27 and 27 & 28 deletion | 0 | Severe |
| c.4540-8T>A | r.4539_4540ins4540-6_4540-1 | p.(Gln1513insProGln) | Exon 31 elongation | 0 | Severe |
| c.4848+3A>G | r.[4774_4848del,=] | p.[Gly1592_Lys1616del,=] | Exon 34 skipping | 10 | Severe |
| c.5461-6T>G | r.5461_5714del | p.(Thr1821Aspfs*6) | Exon 39 & 40 skipping | 0 | Severe |
| c.5898+5G>A | r.[=,5898_5899ins_5899+1_5890-1,5898_5899ins5899+1_5899+170] | p.[=,Cys1967Valfs*24] | Exon 42 elongation and intron 42 retention | 48 | Moderate |
| c.6386+3A>G | r.[6386_6387ins6386+1_6387-1,6340_6386del,=] | p.[Ser2129Serfs*29, Val2114Hisfs*4,=] | Exon 46 elongation and 47-nt reduction | 26 | Moderately severe |

C. Novel causal deep-intronic *ABCA4* variants
| DNA variant | RNA effect | Protein effect | % WT mRNA | Proposed severity |
| --- | --- | --- | --- | --- |
| c.67-2023T>G<sup>#</sup> | r.[66_67ins67-2266_67-2024,=] | p.[Ile23Ilefs*30,=] | 33 | Moderate |
| c.570+1798A>G<sup>#</sup> | r.570_571ins570+1733_570+1797 | p.(Phe191Leufs*6) | 0 | Severe |
| c.769-788A>T | r.[768_769ins769-778_769-617,=] | p.[Leu257Aspfs*3,=] | 4 | Severe |
| c.859-640A>G | r.858_859ins859-685_859-640 | p.(Phe287Tyrfs*69) | 0 | Severe |
| c.859-546G>A | r.[858_859ins859-545_859-685,=] | p.[Phe287Tyrfs*33,=] | 36 | Moderate |
| c.1937+37C>G | r.1937_1938ins1938+1_1938+36 | p.(Phe647*) | 0 | Severe |
| c.1938-621G>A | r.[=,1937_1938ins1938-797_1938-624,<br>1937_1938ins[1937+396_1937+529;1938-797_1938-624] | p.[=,Phe647Alafs*22,Phe647Serfs*22] | 93 | Mild |
| c.1938-514A>G | r.[1937_1938ins[1937+396_1937+529;1938-623_1938-515],1937_1938ins1938-623_1938-515,=] | p.[Phe647Serfs*155,Phe647Serfs*22,=] | 13 | Severe |
| c.2588-706C>T | r.[2587_2588ins2588-839_2588-708,=] | p.[Gly863Alafs*3,=] | 5 | Severe |
| c.3863-1064A>G<sup>\$</sup> | r.(?) | p.(?) | n.a. | Unknown |
| c.4634+741A>G | r.[4634_4635ins4634+614_4634+740,=] | p.[Ser1545Serfs*51,=] | 11 | Severe |
| c.6148-84A>T | r.[=,6147_6148ins6148-262_6148-90,6006_6147delins6148-310_6148-90,6148_6149del] | p.[=,Val2050Valfs*68,Ile2003Hisfs*30,<br>Val2050_Leu2094del] | 43 | Moderate |
| c.6283-78G>T | r.[=,6283_6283ins6283-282_6283-80] | p.[=,Asp2095Aspfr*12] | 88 | Mild |

**Fig. 1:**
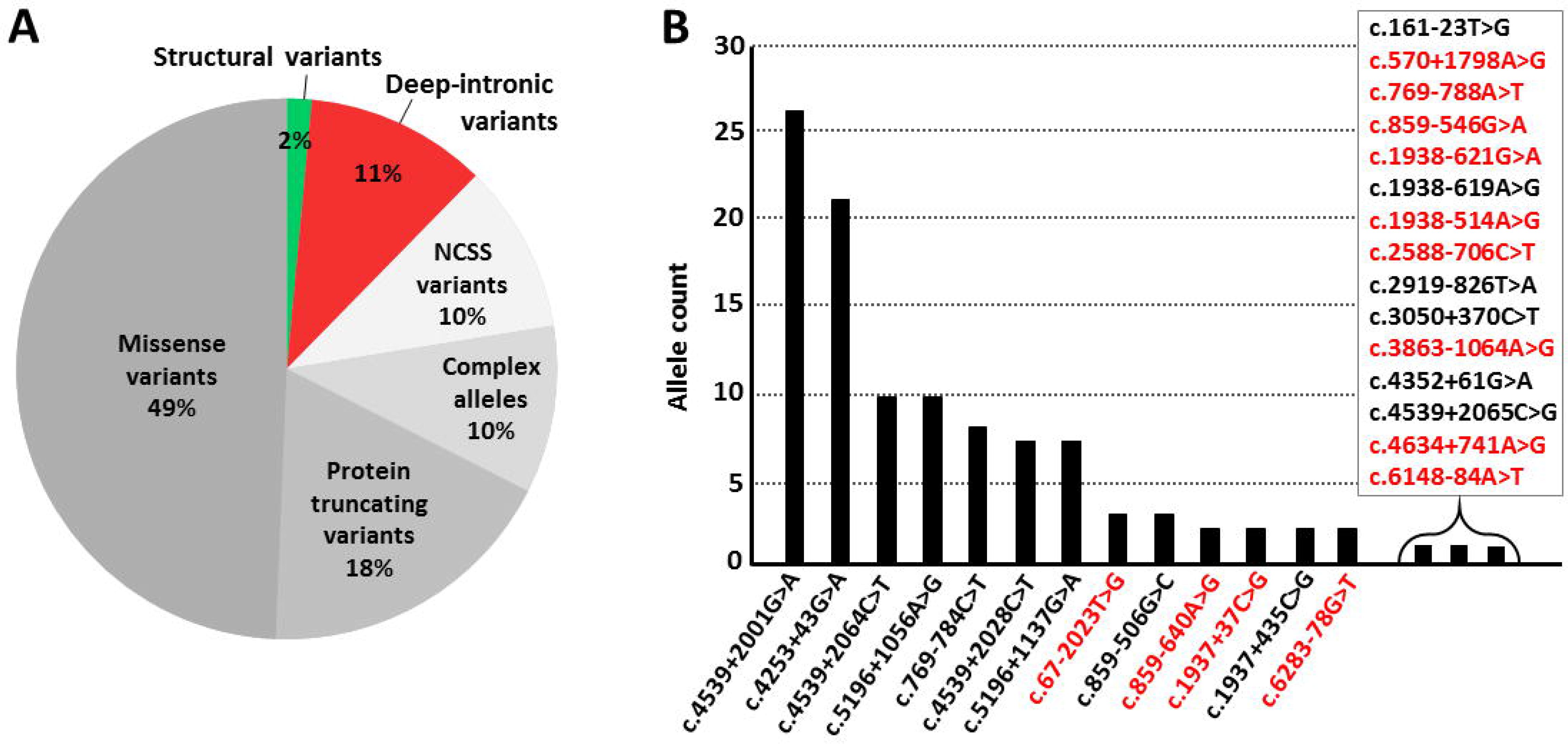
Distribution of different types of alleles and deep-intronic variants in *ABCA4*. **A.** The contribution of each type of variant or allele in bi-allelic, and mono-allelic cases except those carrying c.5603A>T, is represented. Protein truncating variants comprise nonsense, frameshift and canonical splice site variants. The 10% complex alleles only consist of combinations of missense variants, the most frequent of which were c.[1622T>C;3113C>T] (n=30; 27% of all complex alleles) and c.[4469G>A;5603A>T] (n=27; 25% of all complex alleles). They do not include the complex alleles which contain non-canonical splice site (NCSS) variants, deep-intronic variants or protein truncating variants, when present in *cis* with other variants. If these would have been included, 16% of the alleles would consist of complex alleles. **B.** Deep-intronic variant allele count in this study. Novel deep-intronic variants are highlighted in red. One hundred and seventeen causal deep-intronic variants were identified. The deep-intronic variants c.4539+2001G>A (n=26) and c.4253+43G>A (n=21) were found most frequently. Most of the novel deep-intronic variants were found in single STGD1 probands.

Two (likely) pathogenic variants were found in 326 probands, three of them carrying p.Asn1868Ile in a homozygous manner, and one (likely) causal variant in *trans* with p.Asn1868Ile was found in 125 probands. Only one (likely) causal variant was identified in 174 probands. In 65, the p.Asn1868Ile variant was the only identified variant (Supplemental Table S7). No (likely) causal variants were found in 364 cases.

Among the SNVs, the most common causal alleles were c.5603A>T (n=134), c.5882G>A (n=84), c.[5461−10T>C;5603A>T] (n=44), c.[1622T>C;3113C>T] (n=30), c.[4469G>A;5603A>T] (n=27), c.4539+2001G>A (n=26), c.6079C>T (n=23) and c.4253+43G>A (n=21) (Supplemental Table S6). To visualize the relative frequency of causal STGD1-causing alleles, we excluded 65 heterozygous c.5603A>T alleles that were found as the only *ABCA4* allele in these cases, as they were most likely present because of its high allele frequency (0.06) in the general population (Zernant et al. 2017; Cremers et al. 2018; Runhart et al. 2018) (Supplemental Fig. S2).

### Splice defects due to non-canonical splice site variants

The effect on splicing of nine NCSS variants was tested in nine wild-type splice constructs previously described (Sangermano et al. 2018) (Supplemental Fig. S3). All of the nine tested novel NCSS variants showed a splice defect when tested in HEK293T cells. Severity was assigned according to the percentage of remaining WT mRNA, as described previously (Sangermano et al. 2018). Five NCSS variants were deemed severe as they showed no or less than 15% of the WT mRNA, three were considered moderately severe with WT RNA present between 26% and 51%, and one was deemed mild as it resulted in 69% WT mRNA (Table 1B; Supplemental Fig. S4).

### Deep-intronic variants identification and functional characterization

Fifty-eight DI variants were selected for splice assays. Most of the DI variants were selected as they strengthen an already existing cryptic splice acceptor site (SAS) or splice donor site (SDS) or create a new putative splice site with a relative strength of at least 75% in two of five splice strength prediction algorithms. To test their effects, 27 WT midigenes splice constructs were employed, 23 of which were described previously (Sangermano et al. 2018), and four of which were new (Supplemental Fig. S3).

Thirteen of 58 tested DI variants showed a splice defect upon reverse transcription (RT)-PCR and Sanger validation (Table 1C; Fig. 2). For the variants that did not show any splice defect, RT-PCR results are shown in Supplemental Fig. S5.

**Fig. 2:**
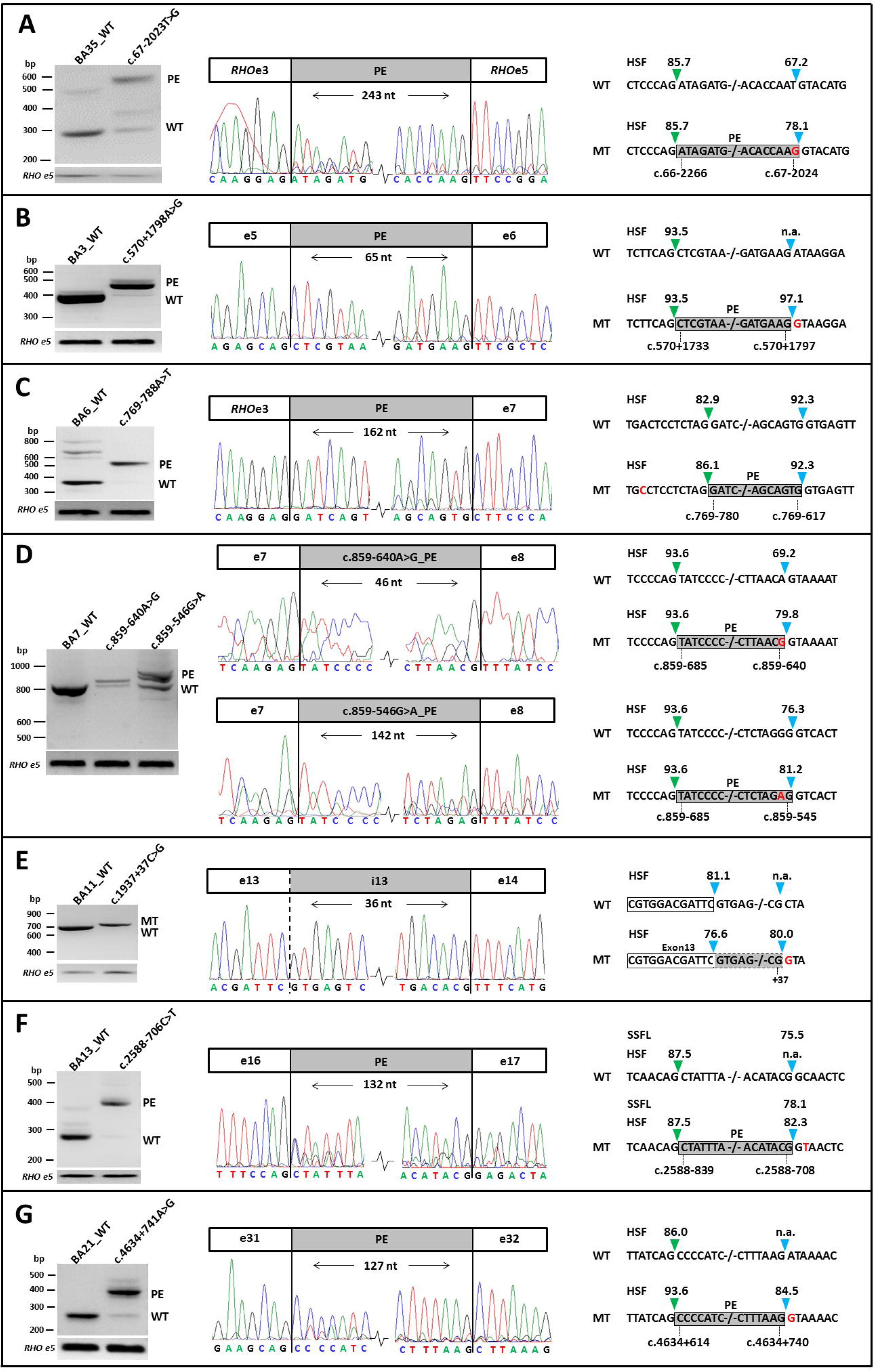
Novel splice defects due to deep-intronic *ABCA4* variants. Wild-type (WT) and mutant (MT) midigenes were transfected in HEK293T cells and the extracted RNA was subjected to RT-PCR. Left panels show the *ABCA4*-specific RT-PCR products with Rhodopsin exon 5 (*RHO* e5) RT-PCR as a transfection efficiency control. In the middle panels, Sanger sequencing results of the RT-PCR products are given. At the right side pseudo-exons (PE) and an exon elongation are depicted with splice site strength predictions for WT and MT, with green rectangles representing the splice acceptor sites and blue rectangles representing the splice donor sites. Red highlighted nucleotides represent the mutations. Except for c.1937+37C>G (**D**), which resulted in a 36-nt exon 13 elongation, all deep-intronic variants lead to PEs. The intron 7 variants in part **C** result in partially overlapping PEs that share the same splice acceptor site at position c.859−685. HSF, human splicing finder; SSFL, slice site finder like; PE, pseudoexon; n.a., not applicable.

Six of the novel DI variants, i.e. variants c.570+1798A>G, c.769−788A>T, c.859−640A>G, c.1938−514A>G, c.2588−706C>T and c.4634+741A>G, resulted in out of frame PE inclusions in the RNA and were deemed severe (Fig. 2, Fig. 3). Variants c.67−2023T>G and c.859−546G>A were classified as moderately severe as 33% and 36% of the WT products were present, respectively. As predicted due to the presence of a downstream cryptic SDS, variant c.1937+37C>G led to an elongation of exon 13 by 36 nucleotides, which resulted in the introduction of a premature stop codon (p.Phe647*). Moreover, two intron 13 variants, c.1938−621G>A and c.1938−514A>G, showed a complex splice pattern and led to the generation of two mutant transcripts each (Fig. 3A-C). Each of these products contained a shared PE of 134 nt (PE1) as well as mutation-specific PEs, denoted PE2 (174 nt) or PE3 (109 nt) for c.1938−621G>A and c.1938−514A>G, respectively (Supplemental Fig. S6). For variant c.1938−621G>A only 7% of the total cDNA product showed PE inclusion whereas for c.1938−514A>G, 87% of the cDNA products were mutant. To investigate the nature of the PE1 insertions, we studied the exon 12-17 segment of the mRNA obtained from PPCs derived from a control person. As depicted in Supplemental Fig. S7, transcripts containing PE1 or PE1 and PE2, were identified when photoreceptor progenitor cells (PPCs) were grown under nonsense-mediated decay-suppressing conditions. The sum quantity of these two products was 2.9% of total mRNA suggesting that there are small amounts of PE insertions involving PE1 in the healthy retina.

**Fig. 3:**
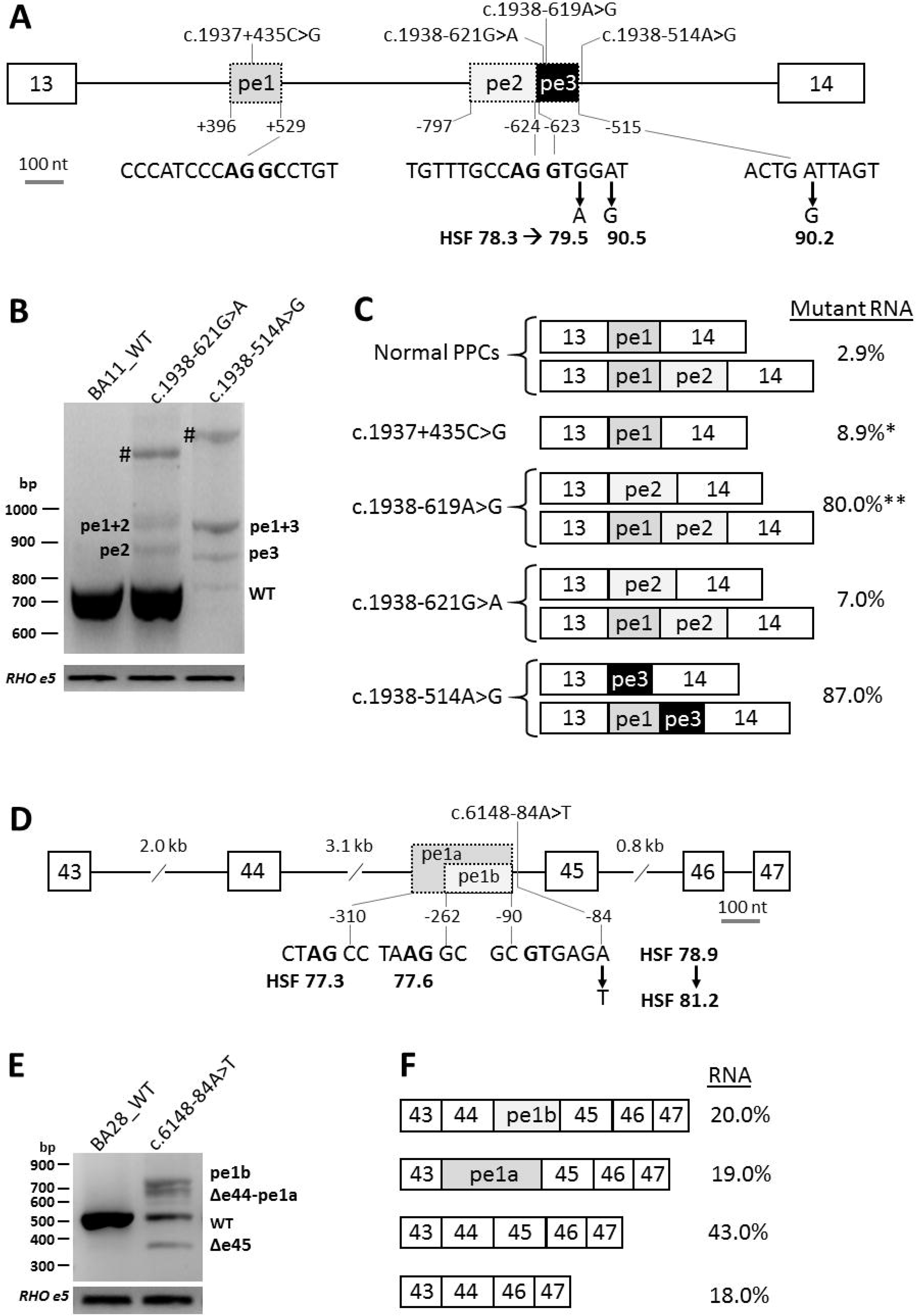
Splice defects due to variants in *ABCA4* intron 13 and 44. **A**. Genomic structure of intron 13 containing three pseudo-exons (PEs) due to four deep-intronic variants. PE2 and PE3 share a splice donor site (for PE2) and splice acceptor site (for PE3). Variants c.1938−621G>A and c.1938n−619A>G strengthen the same cryptic splice donor site of PE2 slightly or strongly, respectively, as based in the Human Splicing Finder (HSF). Variant c.1938−514A>G creates a new strong splice donor site of PE3. The canonical and putative canonical splice sequences are given in bold lettering. The first and last positions of the PEs are provided. **B.** Agarose gel analysis of RT-PCR products for intron 13 variants upon HEK293T cell splice assays. PE2 and PE3 were observed as single insertions, but also in combination with PE1. #Heteroduplex fragments of the lower bands. **C.** Schematic representation of all mutant transcripts identified upon RT-PCR in HEK293T cell splice assays and of PE1 and PE1/PE2 observed as naturally occurring PEs when analyzing photoreceptor progenitor cells (PPCs) derived from a healthy individual. Interestingly, PE1 was previously shown to be induced by variant c.1937+435C>G (*Sangermano et al. 2019) and also can be part of mutant transcripts, together with PE2 or PE3. This is surprising as it is located far upstream of the other causal variants. **Reported by Fadaie et al. 2019. **D.** Variant c.6148−84A>T strengthens a splice donor site and results in PE1a or PE1b by employing upstream or downstream splice acceptor sites, respectively. These splice acceptor sites are comparable in predicted strength based on HSF. The canonical splice sequences are given in bold. **E.** Agarose gel analysis of RT-PCR products due to c.6148−84A>T. The largest fragment shows a 173-nt PE insertion between exons 44 and 45. The second largest band contains a 221-nt PE insertion (PE1a) and skipping of exon 44. The third-largest fragment represents the WT mRNA and the smallest fragment misses exon 45. The relative amounts of the products are listed at the right side.

Intriguingly, DI variant c.6148−84A>T showed a dual effect, leading to the skipping of exon 45 as well as creation of two different PEs. One mutant product showed a 221-nt PE insertion coupled with the skipping of exon 44, while the other mutant product showed the inclusion of a 173-nt PE (Fig. 3D-F). Finally, variant c.3863−1064A>G showed a complex splice pattern compared to the WT and variant c.6283−78G>T led to the insertion of a 203-nt PE in intron 45 (Supplemental Fig. S6). However, the exact boundaries of the presumed PE for variant c.3863−1064A>G could not yet be determined due to technical difficulties.

Overall, 13 novel DI variants were found in 18 alleles. Next to the novel variants, 14 previously reported pathogenic DI variants (Braun et al. 2013; Albert et al. 2018; Bauwens et al. 2019; Fadaie et al. 2019; Sangermano et al. 2019) were found in a total of 99 alleles, details of which are shown in Fig. 1B and Supplemental Table S6.

### Identification of novel structural variants in STGD1 cases

Among 1,054 STGD and STGD-like patients analyzed, we identified 11 unique novel heterozygous SVs, all exon-spanning deletions, in 16 patients (Table 2; Supplemental Table S8 to S13). The corresponding deletions encompass between 1 and 33 exons, ranging from 411 bp to 55.7 kb (Fig. 4). All deletions were found in a heterozygous state in single cases, except the smallest (c.699_768+341del), which encompassed part of exon 6 and 341 bp of intron 6, and was found in six unrelated patients of Spanish origin. Deletion breakpoints were determined employing genomic PCR and Sanger sequencing for 9 of the 11 deletions. Two deletion junctions (deletions 7 and 11) could not be amplified as the 3’ breakpoint was located downstream of the gene beyond the regions targeted by smMIPs. Surprisingly, Sanger sequencing revealed two complex rearrangements as deletions 5 and 6 carried inverted fragments of 279 and 224 bp respectively, residing between large deletions. These small inversions could not be identified with the copy number variant (CNV) detection tool.

**Table 2:** Novel large structural variants in *ABCA4*. For all the novel large deletion variants identified in this study, data are provided with their genomic and DNA positions (hg19), as well as protein effect when possible. Genomic positions indicated are the first and the last deleted nucleotides. Sanger sequencing was performed for all except the ones indicated with an asterisk.

| No. | Genomic position (hg19) | DNA variant | Protein variant | Consequences | Number of cases |
| --- | --- | --- | --- | --- | --- |
| 1 | 94569917-94562911 | c.443-1219_768+1439del | p.(Gly148Alafs*23) | Exons 5 and 6 deletion | 1 |
| 2 | 94565348-94561288 | c.571-801_768+3062del | p.(Phe191_Val256del) | Exon 6 deletion | 1 |
| 3 | 94564419-94564009 | c.699_768+341del | p.(Gln234Phefs*5) | Partial exon 6 deletion | 6 |
| 4 | 94529906-94527518 | c.1555-1033_1937+615delinsAGC | p.(Cys519Phefs*119) | Exon 12 to 13 deletion | 1 |
| 5 | 94532364-94526398 | c.1555-3491_1938-83delins1734_1761-107inv | p.(Cys519Phefs*22) | Exon 12 to 13 deletion and small inverted segment | 1 |
| 6 | 94531301-94522479 | c.1555-2428_2161-101delins2160+7_2160+230invATGAATGins | p.(?) | Exon 12 to 14 deletion and small inverted segment | 1 |
| 7* | 94514513-? | c.(2653+1_2654-1)_(*1_?)del | p.(Gly885Valfs*71) | Exon 18 to 50 deletion | 1 |
| 8 | 94496279-94487503 | c.4254-197_4672delinsGCTTTTT | p.(?) | Exon 29 to exon 33 deletion | 1 |
| 9 | 94472532-94470240 | c.6005+658_6147+757delinsTTTAACAGTGTT | p.(Ser2002Argfs*12) | Exon 44 deletion | 1 |
| 10 | 94467351-94463600 | c.6282+63_6546del | p.(Asp2095_Leu2182del) | Exon 46 to part of exon 48 deletion | 1 |
| 11* | 94461751-? | c.(6729+1_6730-1)_(*1_?)del | p.(Val2244*) | Exon 49 to 50 deletion | 1 |

**Fig. 4:**
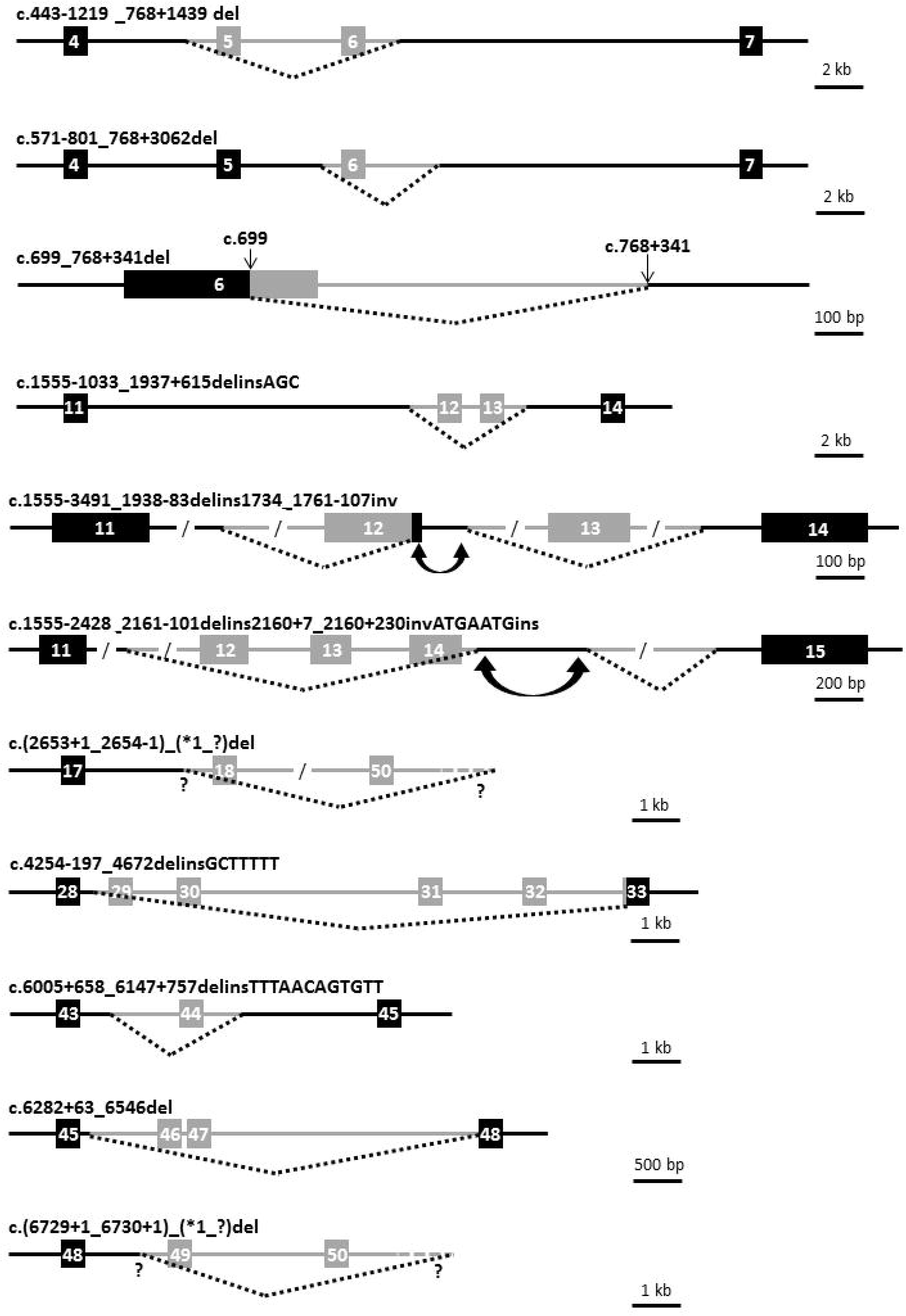
Novel heterozygous structural variants in *ABCA4*. Schematic representation of the 11 structural variants identified. Exons are represented as boxes, black when they are not deleted and grey when they are deleted. Introns are represented as continuous lines, whereas stippled lines depict the deleted regions. Question marks denote that the exact location of breakpoints were unknown. Inverted double arrows represent inverted sequences.

### Microhomologies, repetitive elements and non-B DNA conformation at deletion breakpoints

The breakpoints of the deletions were subjected to extensive bioinformatic analysis to find elements underlying their formation. The presence of microhomology, repetitive elements and non-B DNA conformations was investigated except for deletions 7 and 11 as exact boundaries could not be determined by Sanger sequencing. All other studied SVs presented microhomology at the breakpoint junctions, ranging in size from 1 to 6 bp (Supplemental Fig. S8), four of which presented short insertions (Supplemental Table S14). In eight of 11 (72.7%) of the deletion breakpoints, a known repetitive element was observed, including seven non-Long Terminal Repeats (non-LTR) retrotransposons, among which there were one SINE and four LINEs, three DNA transposons from the *hAT* superfamily and two retrotransposons from the LTR superfamily. However, none of the breakpoints were part of a known element belonging to the same class and no *Alu* sequence was observed at the breakpoint junctions. Finally, the most prevalent non-B conformations observed among our breakpoints are Oligo(G)n tracts as 21 of these repeats were found in seven SVs (Supplemental Tables S14, S15). Inverted repeats were observed in five breakpoint regions. No direct repeats or mirror repeats have been detected, excluding therefore triplex and slipped hairpin structure formation, respectively.

### Uniparental isodisomy of chromosome 1

In STGD1 proband DNA14-33085, a causal homozygous DI variant, c.859−506G>C (p.[Phe287Thrfs*32,=]), was identified. Segregation analysis revealed this variant to be present in his unaffected father, but not in his unaffected mother. To test the possibility that the mother carried a deletion spanning this variant, we performed CNV analysis in the proband’s *ABCA4* gene. No deletion was identified (Supplemental Table S13) and no heterozygous SNPs were observed in or near *ABCA4* in the proband’s DNA. To test whether the chromosome 1 of the father carrying the c.859−506G>C *ABCA4* variant was passed on to the proband as two copies (uniparental isodisomy, UPD), whole exome sequencing was conducted for the proband’s DNA. As shown in Supplemental Fig. S9, chromosome 1 of the proband carries only homozygous SNPs, strongly suggesting the occurrence of UPD.

## Discussion

Employing 3,866 smMIPs, 97.4% of the 128-kb *ABCA4* gene could be sequenced robustly in 1,054 genetically unsolved probands with a STGD or a STGD-like phenotype. In this way, 448 (42.5%) of the probands could be genetically solved. We not only identified nine novel NCSS variants and 13 novel DI variants, but also, facilitated by a high reproducible average read coverage, 11 novel heterozygous SVs. The large set-up of this study allowed us to provide a ‘landscape’ overview of the different types of mutations causing STGD1. As depicted in Fig. 1A, we can appreciate that DI variants constitute a significant cause of STGD1, i.e. 11.0% of the alleles, identified in 23.7% of bi-allelic probands. Deletions constitute 1.8% of alleles and were found in 3.5% of bi-allelic cases. Seven probands carried two DI variants or one DI variant and one SV. Taken together, ‘dark matter’ alleles were found in 116/448 (25.9%) bi-allelic STGD1 probands. However, if we consider only the bi-allelic patients found in the previously unscreened STGD1 cohort, ‘dark matter’ represents 13/182 (7.1%) of the identified alleles. Together, these results strongly argue for a complete sequence analysis of the *ABCA4* gene to fully appreciate its mutational landscape.

### Current state of knowledge on deep-intronic variants in *ABCA4*

Twenty-seven different DI variants were identified in 117 alleles. The 13 novel DI variants were all unique, with the exception of c.67−2023T>G (n=3), and were functionally tested. Variants c.4539+2001G>A, c.4253+43G>A, c.4539+2064C>T and c.5196+1056A>G were the most frequent DI variants in this cohort.

Interestingly, the two DI variants that were found in intron 13, i.e. c.1938−621G>A and c.1938−514G>A, were in close vicinity to two previously described variants, c.1937+435C>G (Sangermano et al. 2019) and c.1938−619A>G (Zernant et al. 2014; Fadaie et al. 2019). As shown in Fig. 3A-C, c.1938−619A>G and c.1938−621G>A led to the same mutant transcripts with PE inclusion. The smaller and larger mutant transcript contained PE2, and PE1 and PE2 together, respectively. Nucleotide change c.1938−619A>G was considered to be a severe variant as it resulted in 80% of mutant product (Fadaie et al. 2019), and c.1938−621G>A, based on the *in vitro* splice assay, was considered to have a mild effect as the majority of the splice products (93%) was normal. The PE resulting from c.1937−514A>G (PE3) is located adjacent to PE2 as they share a dual SAS/SDS (Fig. 3A-C). The involvement of PE1, located 491, 493 and 775 nt upstream of variants c.1937−621G>A, c.1937−619A>G and c.1937−514A>G, respectively, is very surprising. Control PPCs also show a small percentage (2.9%) of mRNAs containing PE1 or PE1-PE2. Interestingly, the SDS of PE1 also can be employed as a SAS which, in theory could render this intronic SAS/SDS a target for recursive splicing (Hafez and Hausner 2015). Together, these findings suggest that there is a ‘natural sensitivity’ for PE1 to be recognized as a PE even if the splice defect is located far downstream. To our knowledge, this has not been described thus far. Also of interest intron 44 variant c.6148−84A>T resulted in a complex splicing pattern involving different PE insertions with or without flanking exon 44 or 45 deletions. Follow-up studies employing patient-derived retinal-like cells are required to validate these complex splicing patterns.

In Supplemental Table S16, we listed all published 349 DI variant alleles (Braun et al. 2013; Zernant et al. 2014; Bauwens et al. 2015; Bax et al. 2015; Schulz et al. 2017; Albert et al. 2018; Zernant et al. 2018; Bauwens et al. 2019; Fadaie et al. 2019; Khan et al. 2019; Nassisi et al. 2019; Sangermano et al. 2019). The three most frequent are c.4253+43G>A (n=100), c.4539+2001G>A (n=64) and c.5196+1137G>A (n=47). When comparing the effect of *ABCA4* DI variants, a fairly high percentage (18/33; 54.5%) affect existing cryptic splice sites, rather than creating new splice sites (15/33; 45.5%). For some DI variants, the splice defects in HEK293T cells or patient-derived PPCs are very small (c.769−784C>T, c.1937+435G>C, c.1937−621G>A) (Runhart et al. 2019; Sangermano et al. 2019) (this study) or smaller than expected (c.4539+2001G>A, c.4539+2028C>T). We hypothesize that retina-specific splice motifs and factors play a role which are largely missing (HEK293T cells) or underrepresented (PPCs) compared to the normal retina.

### Current state of knowledge on structural variants in *ABCA4*

In this study, 11 unique SVs with sizes ranging from 411 bp to 55.7 kb, were readily identified employing an easy-to-use visual detection tool taking advantage of the high number of reads obtained from smMIPs based-sequencing. Although this tool needs further automation to increase its performance for the detection of smaller deletions or duplications, it demonstrated its efficiency for deletions as small as 411 bp. To our knowledge, 36 different SVs have been identified in STGD1 patients (Supplemental Table S17), 23 of which have been described elsewhere. Twenty-five are deletions, ranging in size from 23 bp to complete deletion of the *ABCA4* gene. There are six duplications, ranging from 24 bp to 26 kb, two indels and one small insertion of 24 bp. As shown in Supplemental Fig. S10, these SVs are spread over the entire gene. All SVs are rare, except for deletions spanning exons 20-22 and exon 6, both found in 6 probands, in Belgium/Germany/Netherlands and from Iberic origin, respectively, suggesting founder effects. Fifteen large rearrangements in *ABCA4* have also been reported in the general population databases gnomAD and Database of Genomic Variants (DGV) (Supplemental Table S18). While their sizes range from 53 bp to 45.75 kb, we consider only five of these to have clinical significance as they span one or more exons. One deletion spanning the 5’ UTR, exon 1 and intron 1, was present in gnomAD and DGV with allele frequencies of 4.7 × 10^−5^ and 2.5 × 10^−4^, respectively. In STGD1 cases, the allele frequency of this SV was 4 × 10^−4^.

While recurrent rearrangements are mostly due to non-allelic homologous recombination (NAHR) between two low-copy repeat sequences, non-recurrent events can be the result of i) non-replicative repair mechanisms: like non-homologous end-joining (NHEJ), microhomology-mediated end-joining (MMEJ) and NAHR between repetitive elements (*Alu* or L1 for example); and ii) replicative-based repair mechanisms: fork stalling and template switching, microhomology-mediated break-induced replication, serial replication slippage (SRS) and break-induced SRS (Lupski 2007). Moreover, the local genomic architecture (the presence of microhomology, repetitive elements, sequences forming non-B DNA conformations and sequence motifs) contributes to genomic instability, leading to genomic rearrangements by impairing the replication process. For example, a microhomology of 1–4 bp may facilitate non-homologous end-joining (Lieber 2010) and longer microhomologies of between 5-25 bp may favor MMEJ (McVey and Lee 2008). The assessment of the local architecture of deletions identified in this study lead us to rule out the NAHR hypothesis (as no *Alu* sequence or L1 at any breakpoint was observed) and to propose the NHEJ or replication slippage models as the main implicated mechanisms (Supplemental Table S14; Supplemental Fig. S8). Indeed, the presence of microhomologies < 5 bp in most of the junctions, and of scars characterized by insertion of several random nucleotides, could be a signature for NHEJ. Alternatively, several examples of an impaired replication fork have been noted that supports the replicative-based repair model. Indeed, despite the absence of repetitive elements of the same class at both sides of the breakpoints, their presence may initiate the formation of secondary structures, as repetitive elements could be more difficult to replicate, leading to an increased chance of replication fork stalling or collapsing (Vissers et al. 2009). Finally, Oligo(G)n tracts displayed a significant overrepresentation in the breakpoint regions. Such structures can induce tetraplex formation (Bacolla and Wells 2004) and could also trigger rearrangement.

### Uniparental isodisomy chromosome 1

UPD was found in one STGD1 case in this study, which represents the third STGD1 case showing UPD thus far reported (Fingert et al. 2006; Riveiro-Alvarez et al. 2007). UPD is a rare event, with an estimated occurrence of 1 in 5,000 or even less individuals (Liehr 2010). UPD was also described in six other inherited retinal dystrophy patients in which chromosomes 1, 2 and 6 were implicated (Thompson et al. 2000; Rivolta et al. 2002; Thompson et al. 2002; Wiszniewski et al. 2007; Roosing et al. 2013; Souzeau et al. 2018). We cannot exclude that there are additional UPD cases in our cohort as segregation analysis was not performed for all homozygous cases. Our finding stresses the importance of segregation analysis in the parents’ DNAs as the recurrence risk for future offspring is very low in UPD families.

### Missing heritability

In 174/1,054 (16.5%) of probands, we identified only one (likely) causal allele. In view of the high carrier frequency of *ABCA4* variants in the general population, estimated to be ~5% (Maugeri et al. 1999; Cornelis et al. 2017), about one-third of these mono-allelic cases may be explained in this way. This may even be higher as we intentionally recruited mono-allelic STGD and STGD-like probands for this study. Some causal variants may have escaped our attention. First, we have not focused on variants affecting transcription regulation. Thus far, there is limited evidence for *ABCA4* variants affecting transcription (Bauwens et al. 2019), but the reported putative regulatory variants were not found in this study. As *in silico* tools (Alamut algorithms, SpliceAI (Jaganathan et al. 2019)) may not predict retina-specific splice defects, we may have missed some causal variants. Also, smMIPs-based sequencing may miss heterozygous deletions smaller than ~400 bp and will not detect insertions or inversions larger than ~40 bp. In the absence of variant data for other maculopathy-associated genes, digenic inheritance can also not be disregarded as it was shown in *CNGA3*/*CNGB3*-associated achromatopsia (Burkard et al. 2018). Finally, complex alleles constitute a large fraction (10%) of all *ABCA4* alleles in this study. For some, such as c.[2588G>C;5603A>T] there are strong arguments to believe that these alleles only act as a mild (fully penetrant) allele, when present in *trans* with a severe allele (Zernant et al. 2017). It cannot be excluded that more than two variants, each with a very small contribution for causality, are present in *ABCA4* alleles. More refined functional tests of coding and non-coding *ABCA4* variants are needed to understand the full genetic landscape of STGD1.

The major advantages of smMIPs-based *ABCA4* sequencing compared to WGS are that: i) it is at least an order of magnitude cheaper than WGS, ii) results in much smaller data storage, and iii) requires no separate informed consent regarding incidental findings. However, disadvantages of smMIPs are that: i) it is restricted to one or a few genes if including introns, ii) it is more cost-effective when large series are analyzed, iii) the analysis is only suitable for the detection of CNVs but not for inversions and insertions, and finally, iv) the sequencing procedure and variant calling requires a specialized set-up.

In our study a significant fraction of probands carried one (likely) causal variant or c.5603A>T as a single allele (239; 22.7%) or no causal variant (364; 34.5%). Additionally, three cases were found homozygous for c.5603A>T, suggesting that missing variants can be identified in other macular-related genes. A more comprehensive smMIPs-based screening platform for these STGD-like cases would likely require the sequence analysis of an additional ~80 genes associated with inherited central vision defects.

As shown in this study, smMIPs-based analysis of the complete sequence(s) of one or a few genes implicated in clinically well-defined human diseases may allow the (re)analysis of hundreds to thousands of samples, in particular by targeting cohorts in developing countries in which low-cost analysis is crucial. For instance, the in-depth coverage of smMIPs-based sequencing could be used for molecular diagnosis of the most frequent forms of cancer as it allows the detection of clonal events at very low levels (~5%) (Friedman et al. 2015), as well as the detection of SVs, not only in DNA extracted from leukocytes but also from paraffin embedded tumors, generating fragmented DNA, as shown for *BRCA1/2* and many other cancer genes (Eijkelenboom et al. 2016; Neveling et al. 2017; Weren et al. 2017). A similar approach can be applied to all other frequent monogenic disorders to find missing variants in non-coding regions which can be potential targets for RNA-based molecular therapy such as antisense oligonucleotides (Albert et al. 2018; Bauwens et al. 2019; Sangermano et al. 2019).

In conclusion, comprehensive sequence analysis of *ABCA4* in 1,054 unsolved STGD and STGD-like probands, splice assays in HEK293T cells and SV analysis, resulted in the identification of ‘dark matter’ variants in 26% of bi-allelic STGD1 probands. Novel complex types of splice defects were identified for intron 13 and 44 variants. Together with published causal DI variants and SVs, a detailed genomic and transcriptomic landscape of *ABCA4*-associated STGD1 was thereby established.

## Materials and Methods

### Control individuals and STGD1 probands in pilot and main sequencing studies

To assess the performance of the smMIPs pool, a pilot study was conducted using five control individuals and 15 STGD1 cases (Supplemental Table S1). To further evaluate the sensitivity of the smMIPs-based sequencing method in subsequent sequencing studies, 138 bi-allelic STGD1 samples were included in the study, 123 of which carried *ABCA4* SNVs and 15 of which carried a deletion or duplication in *trans* with other alleles (Supplemental Table S4, S5).

### Unsolved and partially solved STGD1 and macular dystrophy probands

Twenty-one international and four national centers ascertained 1,054 genetically unsolved or partially solved probands with a STGD or STGD-like phenotype. Two hundred twenty-one of these cases were not previously molecularly studied (Supplemental Table S19). We discerned two patient groups. The first patient group consisted of 993 genetically unsolved probands that carried one (n=345) or no (n=648) *ABCA4* allele. For two subjects, DNA was not available and both parents of the probands were studied, assuming autosomal recessive inheritance. The second patient group consisted of 61 ‘partially solved’ probands, carrying the c.5603A>T (p.Asn1868Ile) variant in *trans* with other alleles. This last group was also investigated as it was suspected that there could be unidentified deep-intronic variants in *cis* with c.5603A>T, as the penetrance of c.5603A>T, when in *trans* with a severe *ABCA4* variant, was ~5% in the population (Cremers et al. 2018; Runhart et al. 2018). Samples were collected according to the tenets of the Declaration of Helsinki and written informed consent was obtained for all patients participating in the study. DNA samples were quantified, diluted to 15 ng/μl and visually inspected after agarose gel electrophoresis.

### smMIPs design

For the targeted sequencing of the *ABCA4* gene, as well as 23.9 kb up- and 14.9 kb downstream sequences, 3,866 smMIPs were designed using MIPgen pipeline, as described previously (Boyle et al. 2014). Four hundred and ninety-six of the 3,866 smMIPs spanned known single nucleotide polymorphisms (SNPs) at the extension or ligation arm. Therefore, 3,370 smMIPs effectively capture 110-nt regions. Details of the smMIPs can be found in Supplemental Table S20. All smMIPs were designed in a way that each nucleotide position in principle is targeted by two different smMIPs to increase sensitivity and avoid dropout of alleles due to variants in the smMIP arms.

### Sequencing set-up

First, a pilot study was conducted to assess the performance of smMIPs and coverage per sample. Twenty DNA samples were sequenced on a NextSeq500 platform using an Illumina mid-output sequencing kit (~130 million reads). Thereafter, 1,192 STGD and STGD-like samples were sequenced using 100 ng DNA per sample. Sequencing libraries were prepared in a fully automated manner as described elsewhere (Khan et al. 2019). To sequence these 1,192 cases, six sequencing runs were performed on an Illumina NextSeq500 platform using high-output kits (~400 million reads). The DNA of 21 patients was sequenced twice. In the first attempt, not enough DNA was used which resulted in less than 100,000 reads per sample, whereas the average number of reads for all samples was 1,355,833. These 21 samples were successfully sequenced in the second run with between 641,635 and 2,074,344 reads per sample.

### Data processing

Raw sequencing data (FASTQ) files were used for mapping and variant annotation by an in-house bioinformatics pipeline. Raw reads were aligned to the reference genome (human genome build GRCh37/hg19). Variant calling and annotation were performed as described previously (Khan et al. 2019). Variants were annotated with predicted effects, gene component information as well as frequency information from various population databases.

### Data analysis and variant classification

To select the causative variants, we applied several filters. First, we retained variants with allele frequencies (AFs) <0.005 in different population databases such as dbSNP, the Genome Aggregation Database (gnomAD; http://gnomad.broadinstitute.org/) and Genome of the Netherlands (GoNL). We then applied a filter based on a ‘quality score’ of >200, based on the average number of reads targeting each genomic position in a single run and other standard filters based on the position of SNVs (e.g. exons, canonical splice sites and known NCSS variants). A number of previously listed causal variants with relatively high AFs such as c.2588G>G, c.5461−10T>C, c.5603A>T and c.5882G>A (AFs in gnomAD: 0.00430, 0.00022, 0.04219 and 0.00456 respectively) were selected separately (Cremers et al. 1998; Cornelis et al. 2017; Schulz et al. 2017; Zernant et al. 2017). To assess the variants, *in silico* analysis was performed using online pathogenicity assessment tools such as Sorting Intolerant From Tolerant (SIFT; https://sift.bii.a-star.edu.sg) (Kumar et al. 2009), Polymorphism Phenotyping v2 (PolyPhen-2) (Adzhubei et al. 2010), and Mutation Taster (Schwarz et al. 2014). Among others, Grantham and PhyloP scores were also assessed to predict the physiochemical effect of nucleotide changes on ABCA4 protein and conservation of nucleotide positions, respectively.

For the selection of known coding and non-coding causal variants, *ABCA4* variant databases, such as the Leiden Open Variation database (LOVD; www.lovd.nl/ABCA4) (Cornelis et al. 2017), and the Human Genome Mutation Database (HGMD; http://www.hgmd.cf.ac.uk/ac/index.php), were used. We supplemented the list of SNVs and small indels with additional variants reported in the literature and employed a script to identify these in an automated manner. Novel candidate variants were selected based on AFs and *in silico* predictions. Thereafter, variants were classified according to the guidelines of the American College of Medical Genetics and Genomics and the Association for Molecular Pathology (ACMG/AMP) (Richards et al. 2015).

Similarly, we detected previously reported causal splice variants (NCSS variants and DI variants) by listing all 165 variants previously tested using *in vitro* splice assays (Braun et al. 2013; Bauwens et al. 2015; Sangermano et al. 2016; Sangermano et al. 2018; Bauwens et al. 2019; Fadaie et al. 2019; Khan et al. 2019; Sangermano et al. 2019) or RT-PCR analysis of patient-derived photoreceptor progenitor cells (PPCs) (Sangermano et al. 2016; Albert et al. 2018).

### Candidate splice defect variant selection

Intronic variants located at positions −14 to −3 upstream of exons and at positions +3 to +6 downstream of exons were considered NCSS variants, and variants outside these positions were considered DI variants. Exonic *ABCA4* NCSS variants can also be found within the first or the last two nucleotides of an exon (Cornelis et al, 2017). To select novel NCSS and DI variants for *in vitro* splice assays, the following selection criteria were used: i) minor allele frequency <0.005 in general population databases such as dbSNP, gnomAD and GoNL, ii) difference in wild-type (WT) and mutant splice site scores obtained by *in silico* analysis performed by using five different algorithms including SpliceSiteFinder-like (SSFL), MaxEntScan, NNSPLICE, GeneSplicer, Human Splicing Finder (HSF) via Alamut Visual software version 2.7 (Biosoftware, 2014; Interactive Biosoftware, Rouen, France; www.interactivebiosoftware) (Reese et al. 1997; Pertea et al. 2001; Cartegni et al. 2003; Yeo and Burge 2004; Desmet et al. 2009), iii) additional splice site score comparison between WT and mutant was made by using the SpliceAI program, which unlike other programs searches for altered splice sites up to 5 kb upstream and 5 kb downstream of the variant (Jaganathan et al. 2019), and iv) enrichment of variants’ allele frequencies in the patient cohort compared to general population databases such as gnomAD and GoNL. A list of all the selected candidates for *in vitro* splice assays and details of *in silico* analysis are provided in Supplemental Table S21.

### Midigene-based splice assay

The splice effect of nine NCSS variants and 58 deep-intronic variants was assessed by midigene-based splice assays employing WT constructs described previously (Sangermano et al. 2018), as well as four newly designed midi- and minigenes, namely BA32 (exons 35-38), BA33 (hg19: intron 1, g.94,585,917-94,584,820), BA34 (hg19: intron 1, g.94,582,386-94,581,151) and BA35 (hg19: intron 1, g.94,581,151-94,579,996) (Supplemental Fig. S3; Supplemental Table S22). All constructs contained *ABCA4* genomic sequences amplified from DNA of the bacterial artificial chromosome clone CH17-325O16 (hg19: insert g.94,434,639-94,670,492). The WT clones served as templates to generate mutant constructs by site-directed mutagenesis, which subsequently were validated by Sanger sequencing. Details of mutagenesis primers and Sanger sequencing primers are given in Supplemental Table S23. WT and mutant constructs were transfected in parallel in HEK293T cells. Total RNA was extracted 48 hours post-transfection and subjected to RT-PCR using *ABCA4* exonic primers when possible, as previously described (Sangermano et al. 2018). Rhodopsin exon 5 amplification was used as transfection and loading control. RT-PCR primer sequences can be found in the Supplemental Table S24.

### Copy number variant analysis

An excel script was employed to detect copy number variants (CNVs) using smMIP read depth. Only large (>400 bp) deletions and duplications in principle can be identified, but not insertions and inversions. Therefore the term CNV instead of SV was used in the following section. The read coverage of a single target was first normalized in comparison to all 3,375 targets by dividing the number of reads for a single target by the total number of reads for a given DNA sample. The coverage of each target of a patient was subsequently compared with the coverage of each target in all probands of that series using the following ratio: R = average coverage of each target of a proband/average coverage of each target of all probands in the run. In order to identify large deletions and duplications, the following ranges were defined: homozygous deletions, ratios (R) <0.3; heterozygous deletions, 0.3≤R<0.7; no CNV, 0.8≤R≤1.2; heterozygous duplications, 1.2<R≤1.7; homozygous duplications, R>1.7. To discriminate actual CNVs from artifacts, ratios per target were compared within each patient and among all the patients. Candidate CNVs were considered if at least 6 consecutive targets were deleted or duplicated.

To assess the accuracy of the CNV detection tool, 13 different CNVs spread throughout the *ABCA4* gene were analyzed in a single run in 15 control DNA samples (Supplemental Tables S4; S5). Since 1,192 probands were sequenced in six NextSeq500 runs, CNV analysis was performed for each run independently (Supplemental Tables S8 to S13). CNVs were confirmed by PCR amplification using *Taq* DNA polymerase (Thermofisher Scientific, Waltham, MA, USA) or Q5 High-Fidelity DNA polymerase (New England Biolabs, Ipswich, MA, USA) with an extension time of 45s. Primer details are shown in Supplemental Table S25. Sanger sequencing was performed with the same primers to delineate the breakpoints of the rearrangement.

### Identification of repetitive elements and presence of microhomology at deletion breakpoints

For a better understanding, breakpoint (BP) numbers were assigned according to their 5’-3’ localization. To characterize the flanking regions of the deletion breakpoints and to identify the underlying mechanisms of the SV formation, 150-bp segments flanking each breakpoint were analyzed and submitted to RepeatMasker (http://www.repeatmasker.org) and Censor (https://www.girinst.org/censor/) online software tools. The presence of microhomology at the breakpoints was assessed by a multiple sequence alignment between the junction fragment and the proximal and distal breakpoint regions, using Clustal Omega (https://www.ebi.ac.uk/Tools/msa/clustalo/). To investigate the possible involvement of non-B DNA conformation elements on SV formation, they were searched within the same 150-bp regions surrounding the breakpoint. DNA sequences leading to non-B DNA conformations were assessed with two different online software tools. Quadruplex forming G-Rich Sequences (QGRS) Mapper (http://bioinformatics.ramapo.edu/QGRS/analyze.php) allowed the identification of (Oligo)n tracts forming tetraplex structures and non-B DNA motif search tool (nBMST) (https://nonb-abcc.ncifcrf.gov/apps/nBMST/default) provided location for direct repeats and slipped motifs, G quadruplex forming repeats, inverted repeats and cruciform motifs, mirror repeats and triplex motifs, Z-DNA motifs and short tandem repeats.

### Semi-quantification of RT-PCR products

To quantify the ratios between correct and aberrant RT-PCR products, densitometric analysis was performed using Image J software after gel electrophoresis as described elsewhere (Schneider et al. 2012). Only peaks corresponding to *ABCA4* bands were taken into account (Supplemental Table S26).

### Uniparental disomy detection

In one STGD1 case (DNA14-33085), we previously identified an apparent homozygous causal DI variant (c.859-506G>C) in *ABCA4*, but only one of the parents was shown to carry this variant. Therefore, a deletion search and segregation analysis were performed. Furthermore, to test the hypothesis of uniparental disomy (UPD), a chromosome 1 haplotype was constructed using whole exome sequencing.

## Supporting information

Supplemental Fig. S1. ABCA4 smMIPs coverage

Supplemental Fig. S2. Distribution of ABCA4 alleles

Supplemental Fig. S3. Midigene construct details

Supplemental Fig. S4. Non-canonical splice site variants RT-PCR results

Supplemental Fig. S5. RT-PCR results of variants with no effect

Supplemental Fig. S6. Deep-intronic variants Sanger sequencing results

Supplemental Fig. S7. Control photoreceptor progenitor cells RT-PCR results

Supplemental Fig. S8. Multiple sequence alignment and schematic overview of breakpoint analysis

Supplemental Fig. S9. Uniparental isodisomy analysis

Supplemental Fig. S10. Overview of all known structural variants in ABCA4

Supplemental Table S1. smMIPs pilot study results

Supplemental Table S2. smMIPs coverage per nucleotide position

Supplemental Table S3. Overview of all identified ABCA4 variants and their allele frequency in gnomAD

Supplemental Table S4. Control patient variants in runs 1 to 6 smMIPs sequencing

Supplemental Table S5. Copy number variants in control STGD1 cases

Supplemental Table S6. Overview of all causal ABCA4 variants

Supplemental Table S7. STGD1 cases and the (likely) causal variants and alleles

Supplemental Table S8. Copy number variant analysis ABCA4 sequencing run 1

Supplemental Table S9. Copy number variant analysis ABCA4 sequencing run 2

Supplemental Table S10. Copy number variant analysis ABCA4 sequencing run 3

Supplemental Table S11. Copy number variant analysis ABCA4 sequencing run 4

Supplemental Table S12. Copy number variant analysis ABCA4 sequencing run 5

Supplemental Table S13. Copy number variant analysis ABCA4 sequencing run 6

Supplemental Table S14. In silico breakpoints analysis and underlying mechanisms for copy number variant formation

Supplemental Table S15. Sequences of non-B DNA conformations

Supplemental Table S16. Overview of all published causal deep-intronic ABCA4 variants

Supplemental Table S17. All known ABCA4 structural variants

Supplemental Table S18. Structural ABCA4 variants in gnomAD and DGV

Supplemental Table S19. STGD1 collaborators and cases

Supplemental Table S20. ABCA4 smMIPs details

Supplemental Table S21. Splice variant candidates in silico analysis

Supplemental Table S22. Gateway primers and sequencing primers details

Supplemental Table S23. Mutagenesis primers details

Supplemental Table S24. RT-PCR primers

Supplemental Table S25. Primer sequences used for copy number variant breakpoints identification

Supplemental Table S26. Semi-quantification of RT-PCR products from midigene splice assays

## Acknowledgements

We thank Ellen Blokland, Duaa Elmelik, Emeline Gorecki, Marlie Jacobs-Camps, Charlene Piriou, Mariateresa Pizzo, and Saskia van der Velde-Visser for technical assistance. We thank Béatrice Bocquet, Dominique Bonneau, Krystyna H. Chrzanowska, Hélene Dollfus, Isabelle Drumare, Monika Heusipp, Takeshi Iwata, Beata Kocyła-Karczmarewicz, Atsushi Mizota, Nobuhisa Nao-i, Adrien Pagin, Valérie Pelletier, Rafal Ploski, Agnieszka Rafalska, Rosa Riveiro, Malgorzata Rydzanicz, Blanca Garcia Sandoval, Kei Shinoda, Francesco Testa, Kazushige Tsunoda, Shinji Ueno and Catherine Vincent-Delorme for their cooperation and ascertaining STGD1 cases. We thank Rolph Pfundt for his assistance in WES data analysis. We thank ERN-EYE and ERDC networks, the Japan Eye Genetics Consortium, and the East Asian Inherited Retinal Disease Society.

