## Supplemental Fig. S1. ABCA4 smMIPs coverage for "Resolving the dark matter of *ABCA4* for 1,054 Stargardt disease probands through integrated genomics and transcriptomics"

### Slide 1
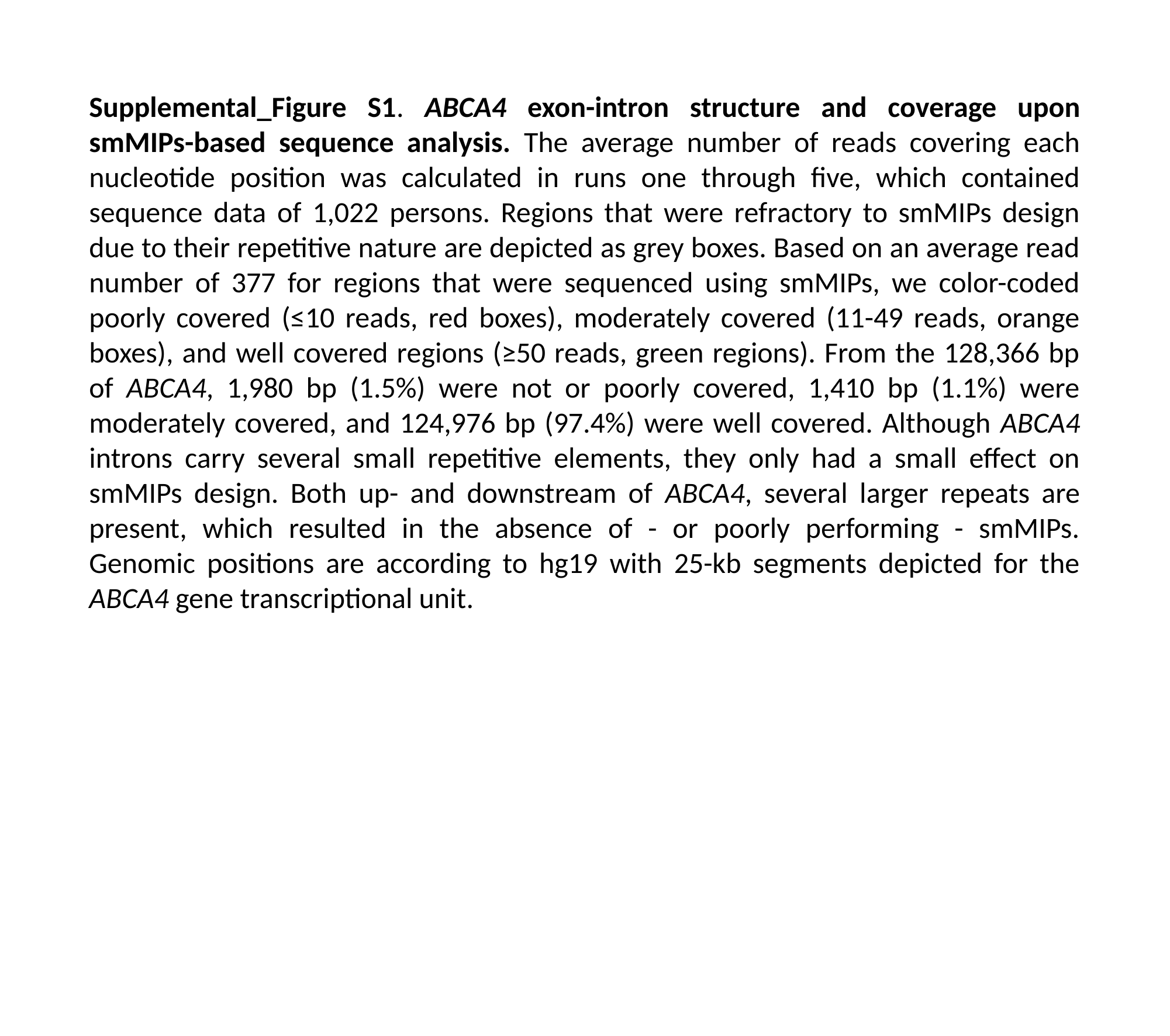

Supplemental_Figure S1. ABCA4 exon-intron structure and coverage upon smMIPs-based sequence analysis. The average number of reads covering each nucleotide position was calculated in runs one through five, which contained sequence data of 1,022 persons. Regions that were refractory to smMIPs design due to their repetitive nature are depicted as grey boxes. Based on an average read number of 377 for regions that were sequenced using smMIPs, we color-coded poorly covered (≤10 reads, red boxes), moderately covered (11-49 reads, orange boxes), and well covered regions (≥50 reads, green regions). From the 128,366 bp of ABCA4, 1,980 bp (1.5%) were not or poorly covered, 1,410 bp (1.1%) were moderately covered, and 124,976 bp (97.4%) were well covered. Although ABCA4 introns carry several small repetitive elements, they only had a small effect on smMIPs design. Both up- and downstream of ABCA4, several larger repeats are present, which resulted in the absence of - or poorly performing - smMIPs. Genomic positions are according to hg19 with 25-kb segments depicted for the ABCA4 gene transcriptional unit.

### Slide 2
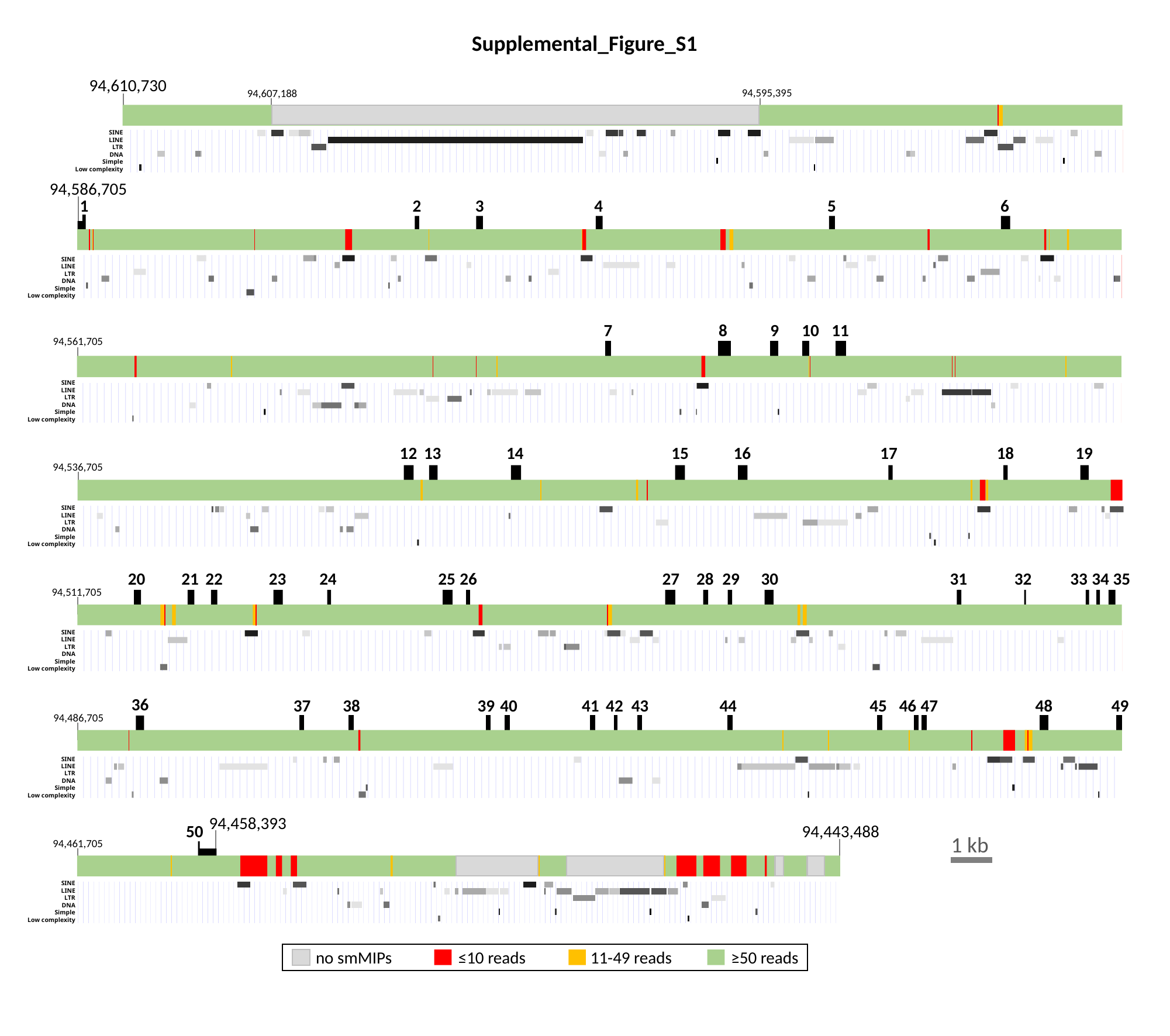

Supplemental_Figure_S1
94,610,730
94,595,395
94,607,188
SINE
LINE
LTR
DNA
Simple
Low complexity
94,586,705
1
2
3
4
5
6
SINE
LINE
LTR
DNA
Simple
Low complexity
7
8
9
10
11
94,561,705
SINE
LINE
LTR
DNA
Simple
Low complexity
12
13
14
15
16
17
18
19
94,536,705
SINE
LINE
LTR
DNA
Simple
Low complexity
20
21
22
23
24
25
26
27
28
29
30
31
32
33
34
35
94,511,705
SINE
LINE
LTR
DNA
Simple
Low complexity
36
37
38
39
40
41
42
43
44
45
47
48
49
46
94,486,705
SINE
LINE
LTR
DNA
Simple
Low complexity
94,458,393
94,443,488
50
1 kb
94,461,705
SINE
LINE
LTR
DNA
Simple
Low complexity
no smMIPs
≤10 reads
11-49 reads
≥50 reads
