## Supplementary figures and images for "Resolving the dark matter of *ABCA4* for 1,054 Stargardt disease probands through integrated genomics and transcriptomics"

### Supplemental Fig. S2. Distribution of ABCA4 alleles

#### Slide 1
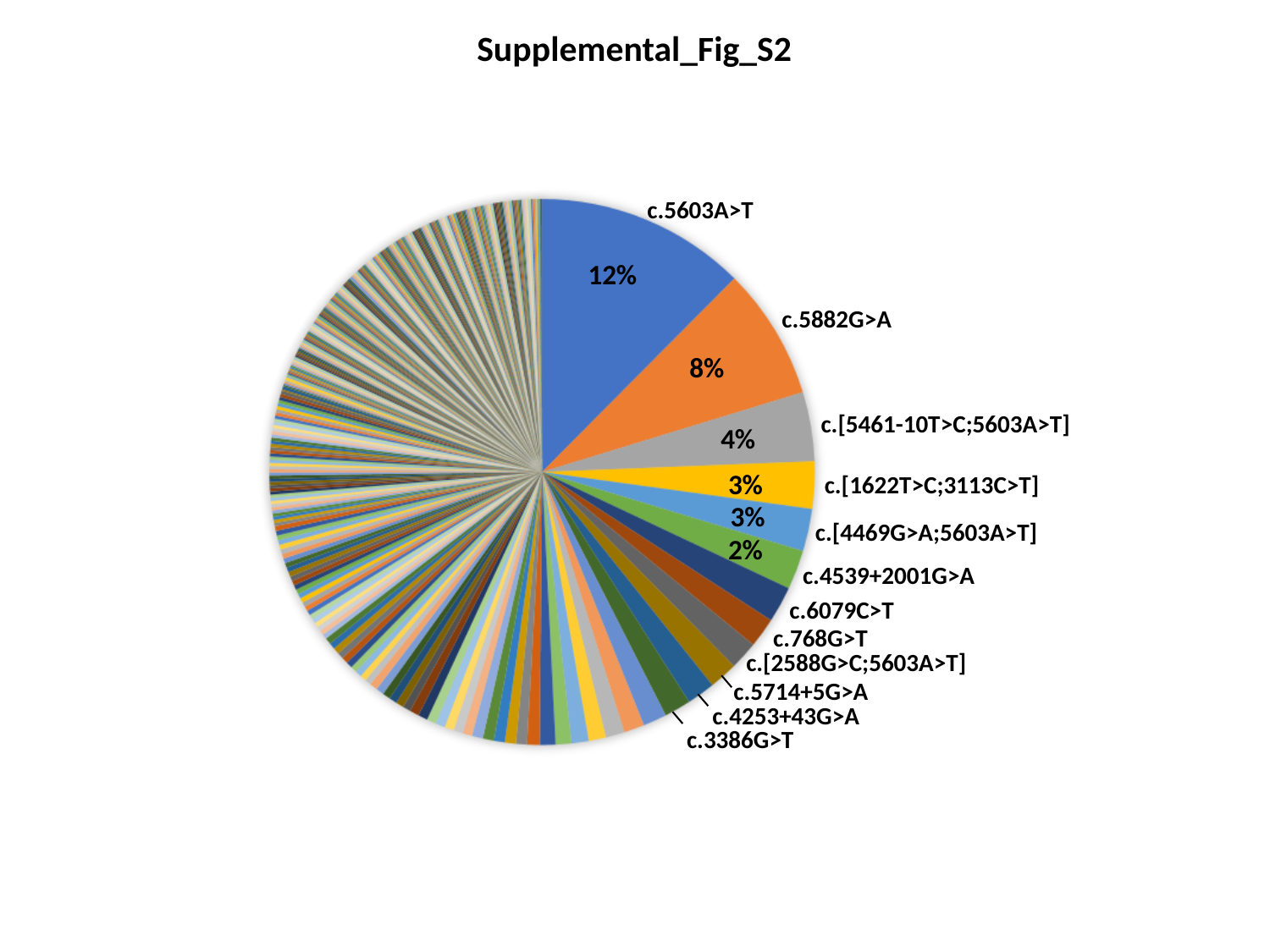

Supplemental_Fig_S2
c.5603A>T
12%
c.5882G>A
8%
c.[5461-10T>C;5603A>T]
4%
3%
c.[1622T>C;3113C>T]
3%
c.[4469G>A;5603A>T]
2%
c.4539+2001G>A
c.6079C>T
c.768G>T
c.[2588G>C;5603A>T]
c.5714+5G>A
c.4253+43G>A
c.3386G>T
