## Supplemental Fig. S3. Midigene construct details for "Resolving the dark matter of *ABCA4* for 1,054 Stargardt disease probands through integrated genomics and transcriptomics"

### Slide 1
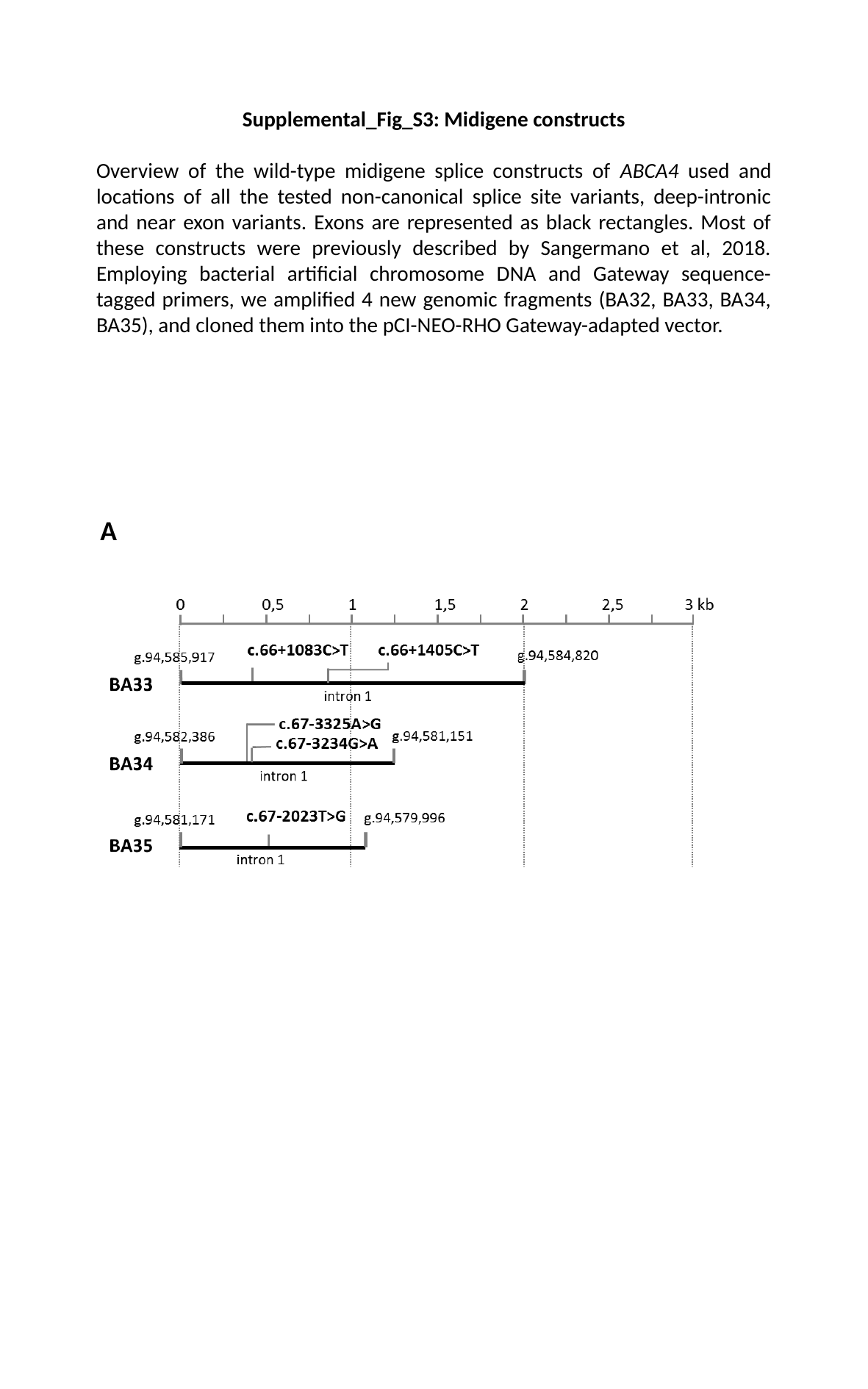

Supplemental_Fig_S3: Midigene constructs
Overview of the wild-type midigene splice constructs of ABCA4 used and locations of all the tested non-canonical splice site variants, deep-intronic and near exon variants. Exons are represented as black rectangles. Most of these constructs were previously described by Sangermano et al, 2018. Employing bacterial artificial chromosome DNA and Gateway sequence-tagged primers, we amplified 4 new genomic fragments (BA32, BA33, BA34, BA35), and cloned them into the pCI-NEO-RHO Gateway-adapted vector.
A

### Slide 2
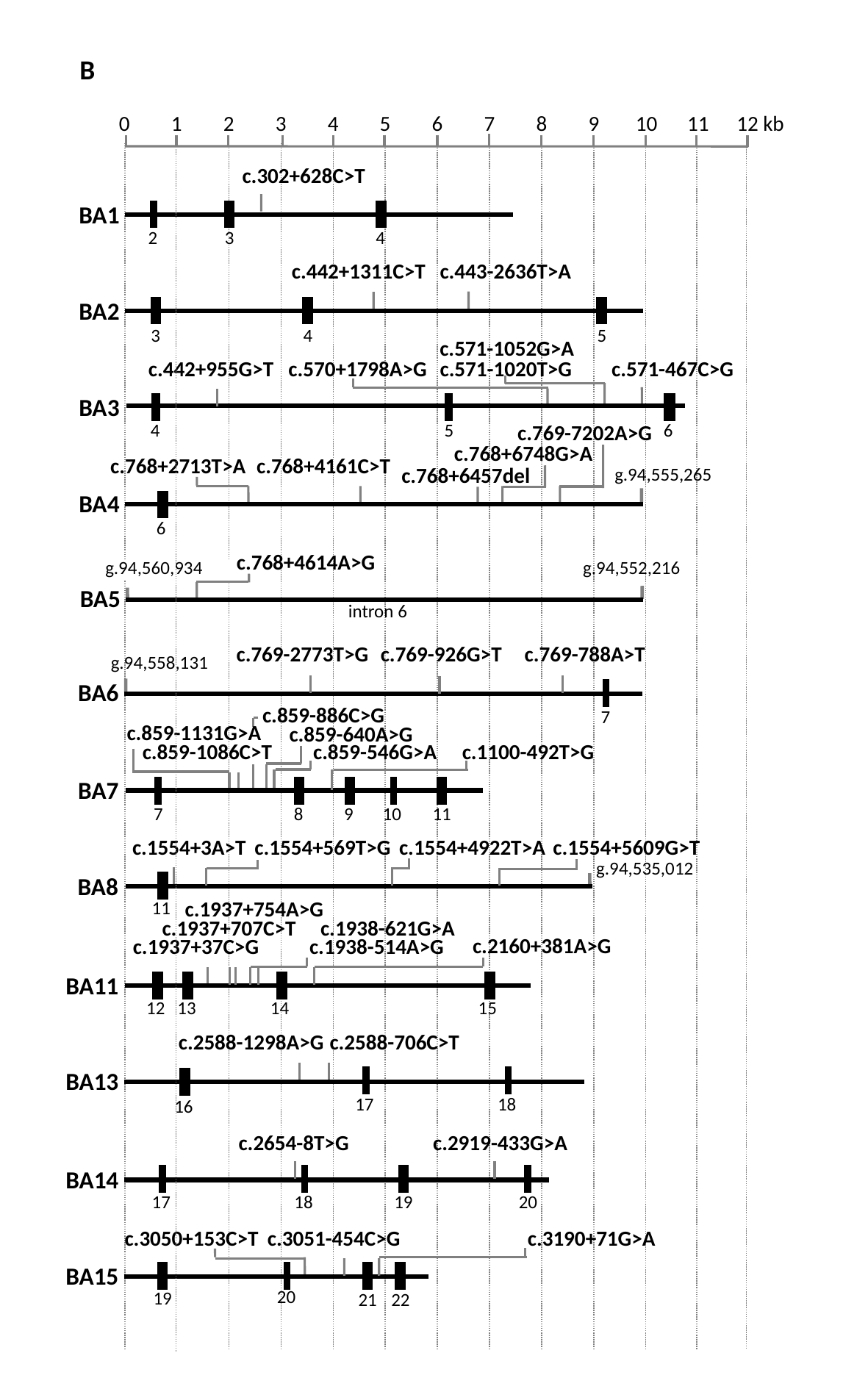

B
0
1
2
3
4
5
6
7
8
9
10
11
12 kb
c.302+628C>T
BA1
2
3
4
c.442+1311C>T
c.443-2636T>A
BA2
3
4
5
c.571-1052G>A
c.442+955G>T
c.570+1798A>G
c.571-1020T>G
c.571-467C>G
BA3
4
5
6
c.769-7202A>G
c.768+6748G>A
c.768+2713T>A
c.768+4161C>T
c.768+6457del
g.94,555,265
BA4
6
c.768+4614A>G
g.94,560,934
g.94,552,216
BA5
intron 6
c.769-788A>T
g.94,558,131
BA6
7
c.769-2773T>G
c.769-926G>T
c.859-886C>G
c.859-1131G>A
c.859-640A>G
BA7
7
8
9
10
11
c.859-1086C>T
c.859-546G>A
c.1100-492T>G
c.1554+3A>T
c.1554+569T>G
c.1554+4922T>A
c.1554+5609G>T
g.94,535,012
BA8
11
c.1937+754A>G
c.1937+707C>T
c.1938-621G>A
c.2160+381A>G
c.1938-514A>G
c.1937+37C>G
BA11
12
13
14
15
c.2588-1298A>G
c.2588-706C>T
BA13
17
18
16
c.2654-8T>G
c.2919-433G>A
BA14
17
18
19
20
c.3050+153C>T
c.3051-454C>G
c.3190+71G>A
BA15
20
19
21
22

### Slide 3
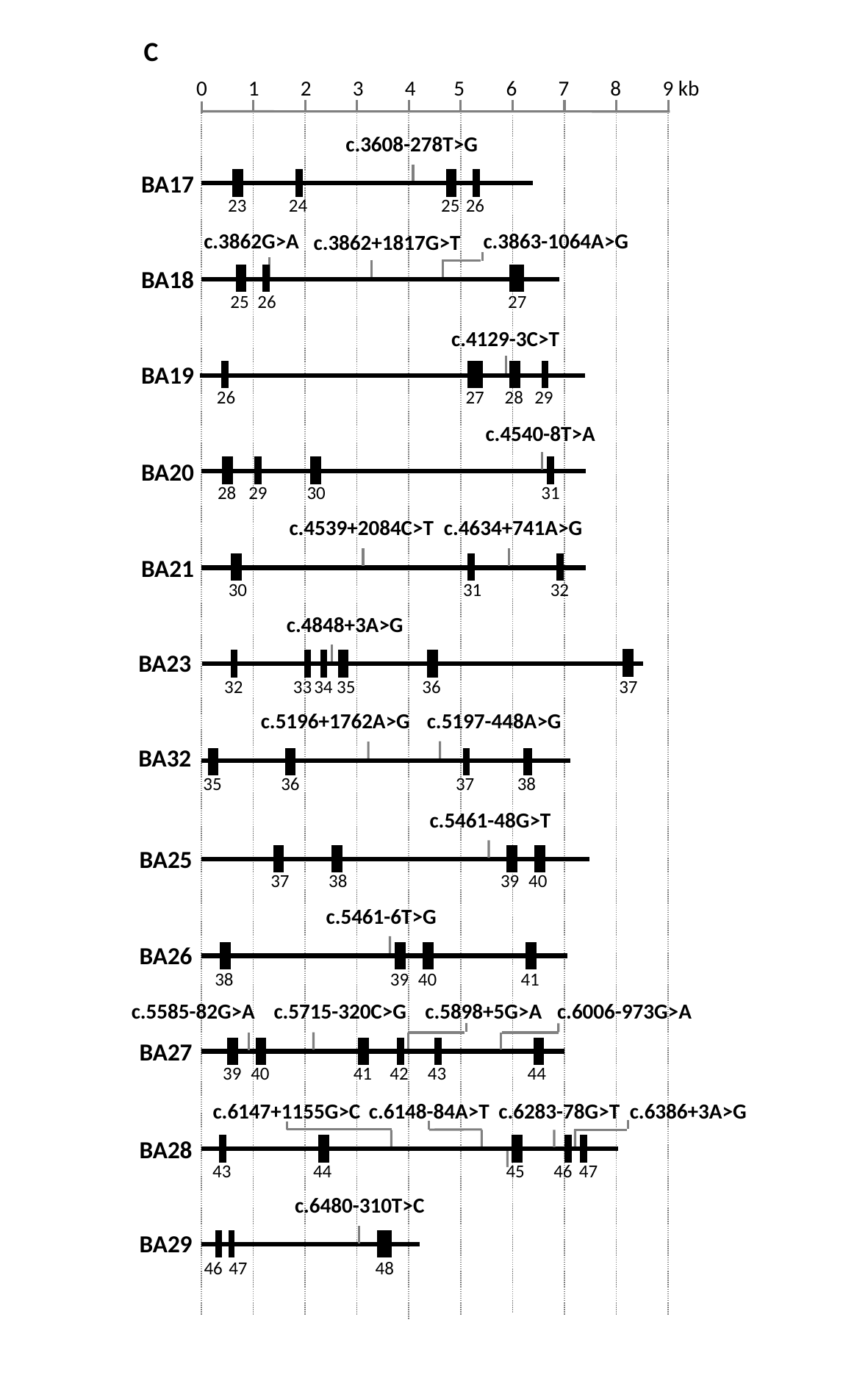

C
0
1
2
3
4
5
6
7
8
9 kb
c.3608-278T>G
BA17
23
24
25
26
c.3862G>A
c.3863-1064A>G
c.3862+1817G>T
BA18
25
26
27
c.4129-3C>T
BA19
26
27
28
29
c.4540-8T>A
BA20
28
29
30
31
c.4539+2084C>T
c.4634+741A>G
BA21
30
31
32
c.4848+3A>G
BA23
32
33
34
35
36
37
c.5196+1762A>G
c.5197-448A>G
BA32
35
36
37
38
c.5461-48G>T
BA25
40
37
38
39
c.5461-6T>G
BA26
40
41
38
39
c.5585-82G>A
c.5715-320C>G
c.5898+5G>A
c.6006-973G>A
BA27
40
41
39
42
43
44
c.6147+1155G>C
c.6148-84A>T
c.6283-78G>T
c.6386+3A>G
BA28
47
43
44
46
45
c.6480-310T>C
BA29
46
47
48
