## Supplemental Fig. S4. Non-canonical splice site variants RT-PCR results for "Resolving the dark matter of *ABCA4* for 1,054 Stargardt disease probands through integrated genomics and transcriptomics"

### Slide 1
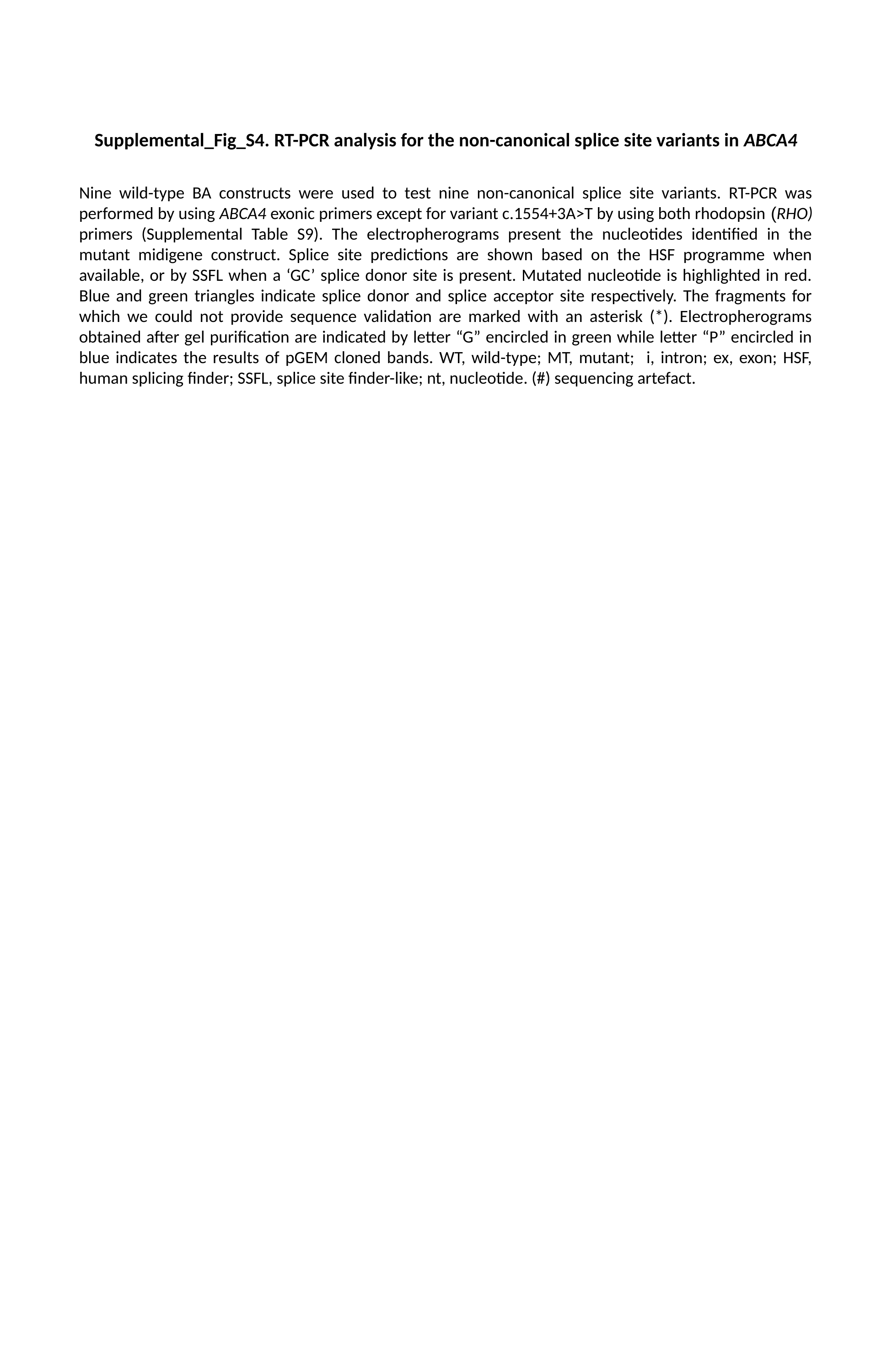

Supplemental_Fig_S4. RT-PCR analysis for the non-canonical splice site variants in ABCA4
Nine wild-type BA constructs were used to test nine non-canonical splice site variants. RT-PCR was performed by using ABCA4 exonic primers except for variant c.1554+3A>T by using both rhodopsin (RHO) primers (Supplemental Table S9). The electropherograms present the nucleotides identified in the mutant midigene construct. Splice site predictions are shown based on the HSF programme when available, or by SSFL when a ‘GC’ splice donor site is present. Mutated nucleotide is highlighted in red. Blue and green triangles indicate splice donor and splice acceptor site respectively. The fragments for which we could not provide sequence validation are marked with an asterisk (*). Electropherograms obtained after gel purification are indicated by letter “G” encircled in green while letter “P” encircled in blue indicates the results of pGEM cloned bands. WT, wild-type; MT, mutant; i, intron; ex, exon; HSF, human splicing finder; SSFL, splice site finder-like; nt, nucleotide. (#) sequencing artefact.

### Slide 2
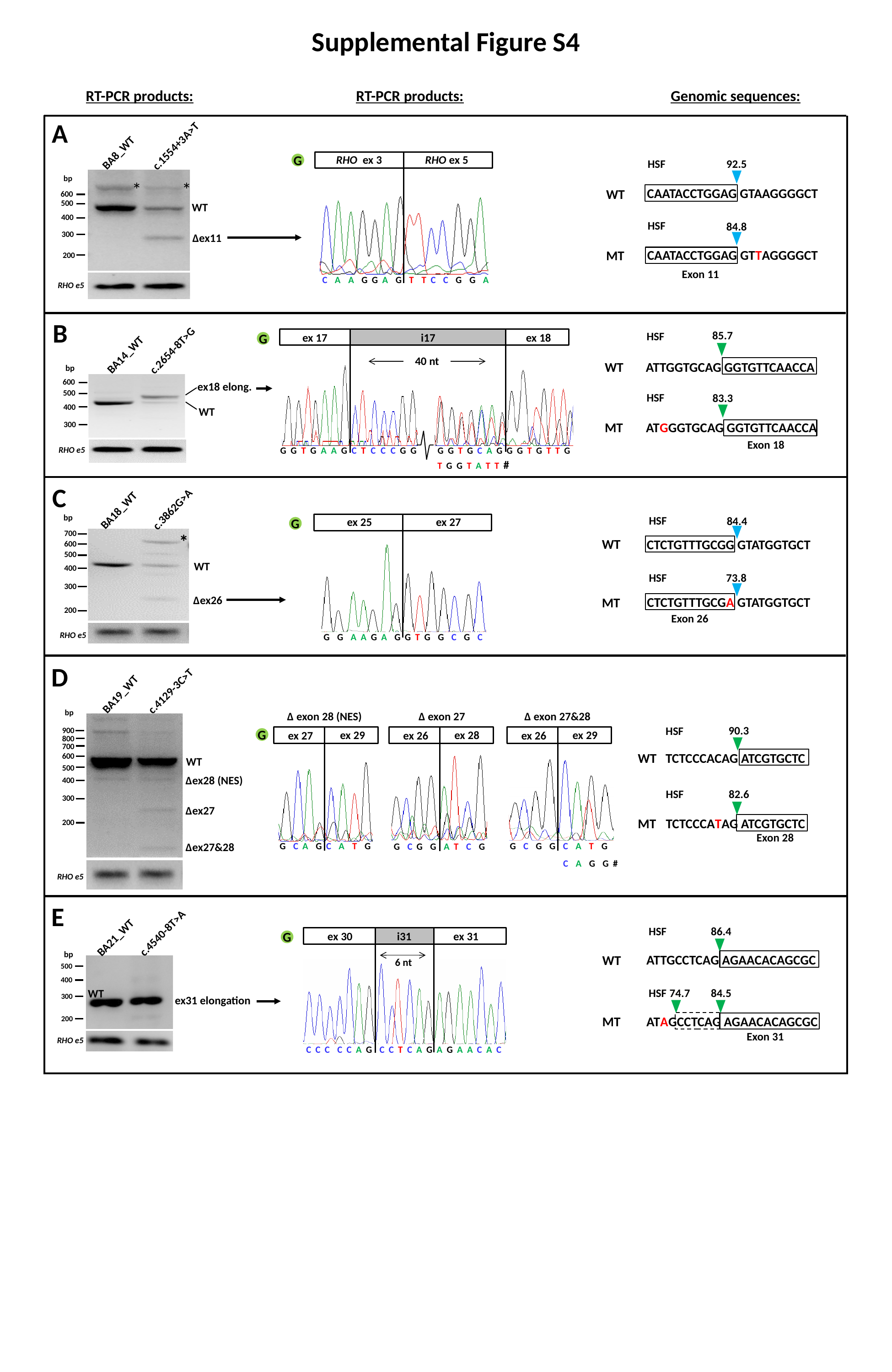

Supplemental Figure S4
RT-PCR products:
RT-PCR products:
Genomic sequences:
c.1554+3A>T
BA8_WT
bp
*
*
600
500
400
300
200
WT
∆ex11
RHO e5
A
G
RHO ex 3
RHO ex 5
C A A G G A G T T C C G G A
92.5
HSF
CAATACCTGGAG GTAAGGGGCT
WT
HSF
84.8
CAATACCTGGAG GTTAGGGGCT
MT
Exon 11
B
c.2654-8T>G
BA14_WT
bp
600
500
400
300
ex18 elong.
WT
RHO e5
85.7
HSF
G
ex 17
i17
 ex 18
40 nt
WT
ATTGGTGCAG GGTGTTCAACCA
HSF
83.3
MT
ATGGGTGCAG GGTGTTCAACCA
Exon 18
G G T G A A G C T C C C G G
G G T G C A G G G T G T T G
T G G T A T T #
C
BA18_WT
c.3862G>A
bp
HSF
84.4
 ex 25
 ex 27
G G A A G A G G T G G C G C
G
700
600
500
400
300
200
*
WT
CTCTGTTTGCGG GTATGGTGCT
WT
HSF
73.8
∆ex26
CTCTGTTTGCGA GTATGGTGCT
MT
Exon 26
RHO e5
D
c.4129-3C>T
BA19_WT
bp
 ∆ exon 28 (NES)
 ∆ exon 27
 ∆ exon 27&28
90.3
HSF
900
G
 ex 29
 ex 28
 ex 29
 ex 27
 ex 26
 ex 26
800
700
TCTCCCACAG ATCGTGCTC
WT
600
WT
500
 ∆ex28 (NES)
400
HSF
82.6
300
 ∆ex27
MT
TCTCCCATAG ATCGTGCTC
200
Exon 28
G C A G C A T G
G C G G C A T G
 ∆ex27&28
G C G G A T C G
C A G G #
RHO e5
E
HSF
86.4
c.4540-8T>A
BA21_WT
500
400
300
200
ex31 elongation
RHO e5
bp
G
ex 30 i31 ex 31
6 nt
 C C C C C A G C C T C A G A G A A C A C
ATTGCCTCAG AGAACACAGCGC
WT
HSF
74.7
84.5
WT
ATAGCCTCAG AGAACACAGCGC
MT
Exon 31

### Slide 3
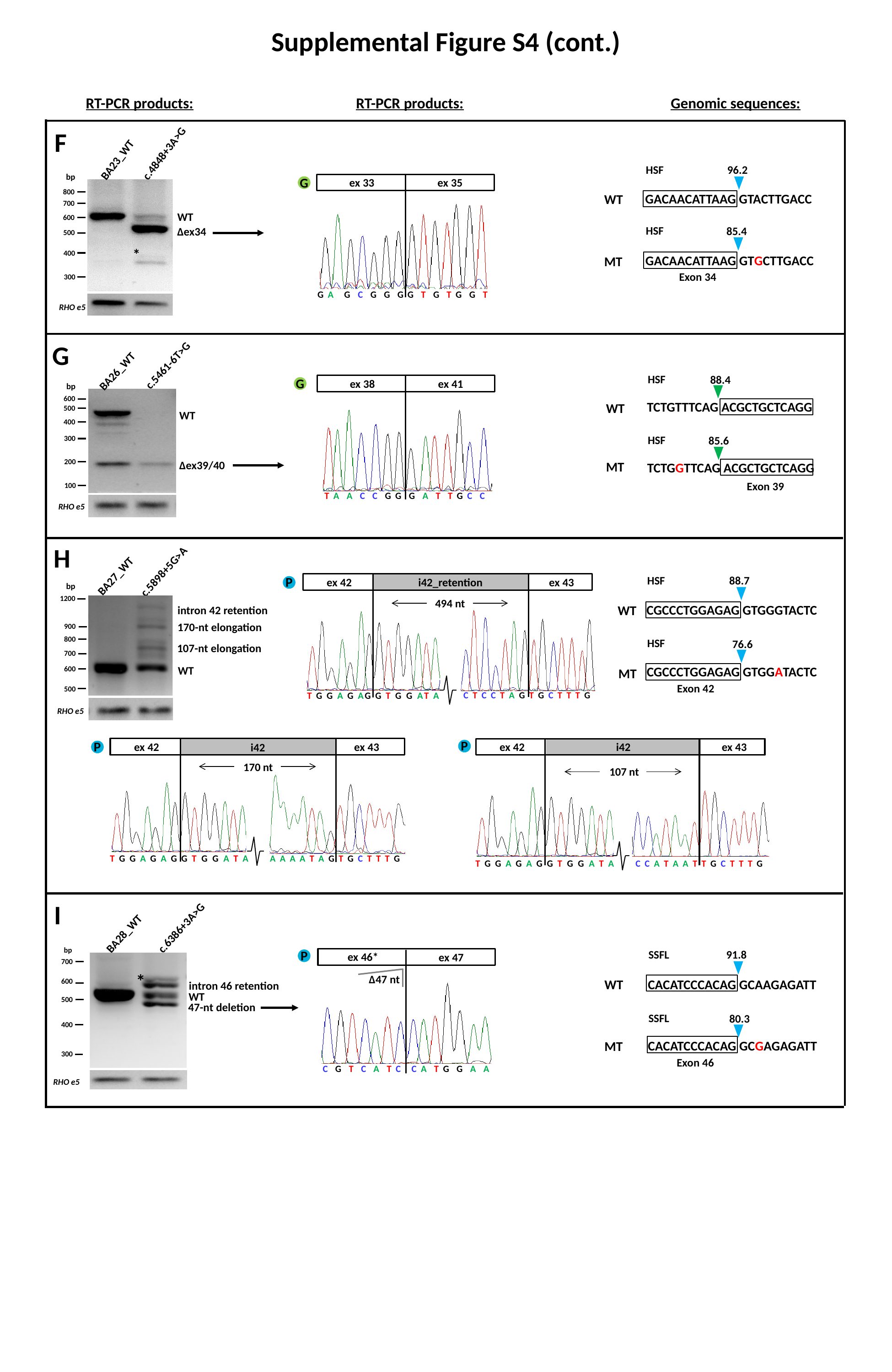

Supplemental Figure S4 (cont.)
RT-PCR products:
RT-PCR products:
Genomic sequences:
F
c.4848+3A>G
BA23_WT
96.2
HSF
bp
G
 ex 33
 ex 35
800
700
600
500
400
300
GACAACATTAAG GTACTTGACC
WT
WT
HSF
85.4
∆ex34
*
GACAACATTAAG GTGCTTGACC
MT
Exon 34
G A G C G G G
G T G T G G T
RHO e5
G
BA26_WT
c.5461-6T>G
HSF
88.4
G
 ex 38
 ex 41
bp
600
500
400
300
200
100
TCTGTTTCAG ACGCTGCTCAGG
WT
WT
HSF
85.6
∆ex39/40
MT
TCTGGTTCAG ACGCTGCTCAGG
Exon 39
 T A A C C G G G A T T G C C
RHO e5
c.5898+5G>A
BA27_WT
1200
900
800
700
600
500
RHO e5
intron 42 retention
170-nt elongation
107-nt elongation
WT
H
88.7
HSF
P
 ex 42
 ex 43
i42_retention
bp
494 nt
CGCCCTGGAGAG GTGGGTACTC
WT
HSF
76.6
CGCCCTGGAGAG GTGGATACTC
MT
Exon 42
T G G A G AG G T G G AT A
C T C C T A G T G C T T T G
P
P
 ex 42
 ex 43
 ex 42
i42
 ex 43
i42
 170 nt
 107 nt
T G G A G A G G T G G A T A
A A A A T A G T G C T T T G
T G G A G A G G T G G A T A
C C A T A A T T G C T T T G
c.6386+3A>G
BA28_WT
bp
700
600
500
400
300
WT
RHO e5
I
P
91.8
SSFL
 ex 46*
 ex 47
 C G T C A T C C A T G G A A
∆47 nt
*
CACATCCCACAG GCAAGAGATT
WT
intron 46 retention
47-nt deletion
SSFL
80.3
CACATCCCACAG GCGAGAGATT
MT
Exon 46
