## Supplemental Fig. S5. RT-PCR results of variants with no effect for "Resolving the dark matter of *ABCA4* for 1,054 Stargardt disease probands through integrated genomics and transcriptomics"

### Slide 1
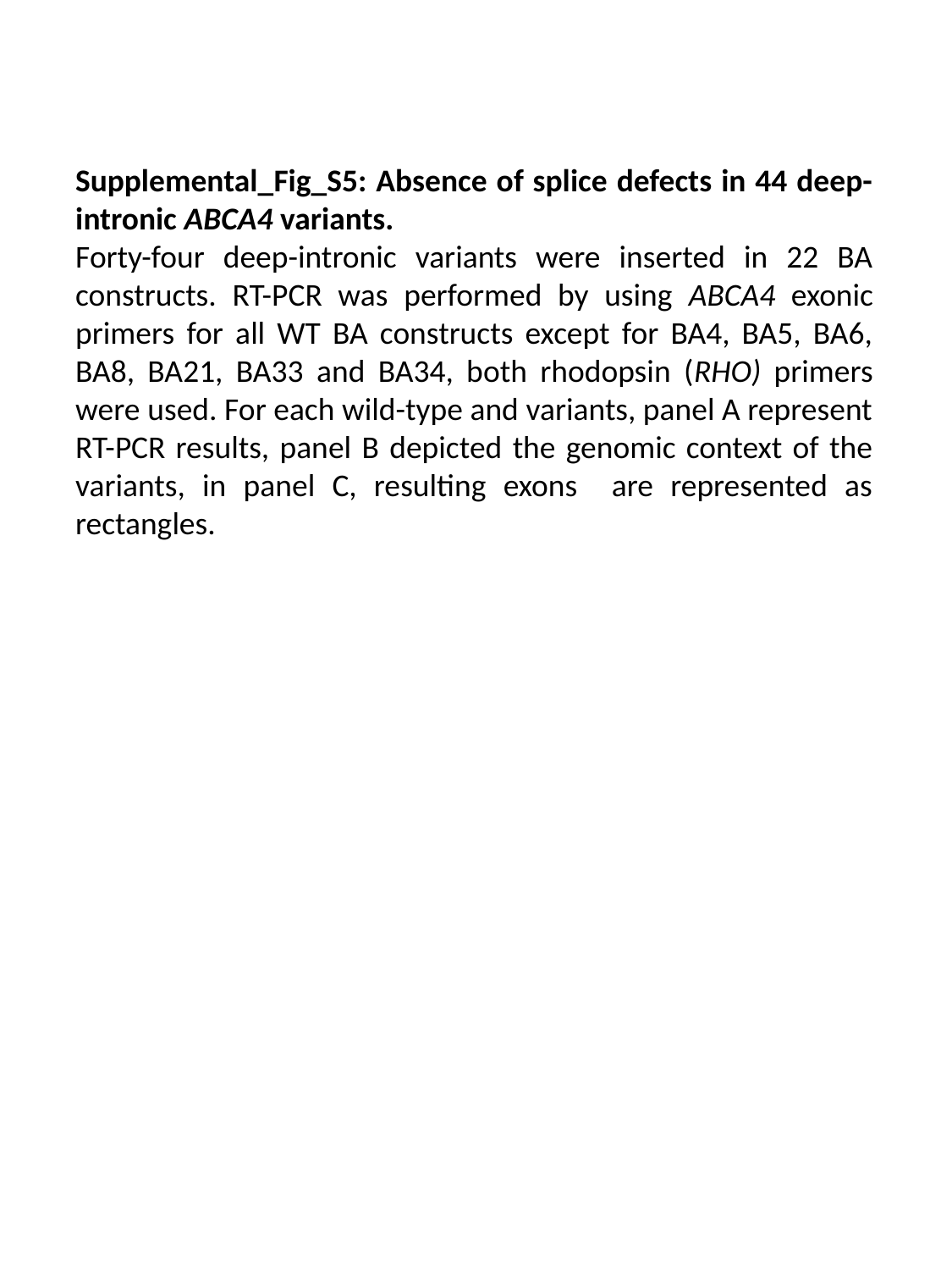

Supplemental_Fig_S5: Absence of splice defects in 44 deep-intronic ABCA4 variants.
Forty-four deep-intronic variants were inserted in 22 BA constructs. RT-PCR was performed by using ABCA4 exonic primers for all WT BA constructs except for BA4, BA5, BA6, BA8, BA21, BA33 and BA34, both rhodopsin (RHO) primers were used. For each wild-type and variants, panel A represent RT-PCR results, panel B depicted the genomic context of the variants, in panel C, resulting exons are represented as rectangles.

### Slide 2
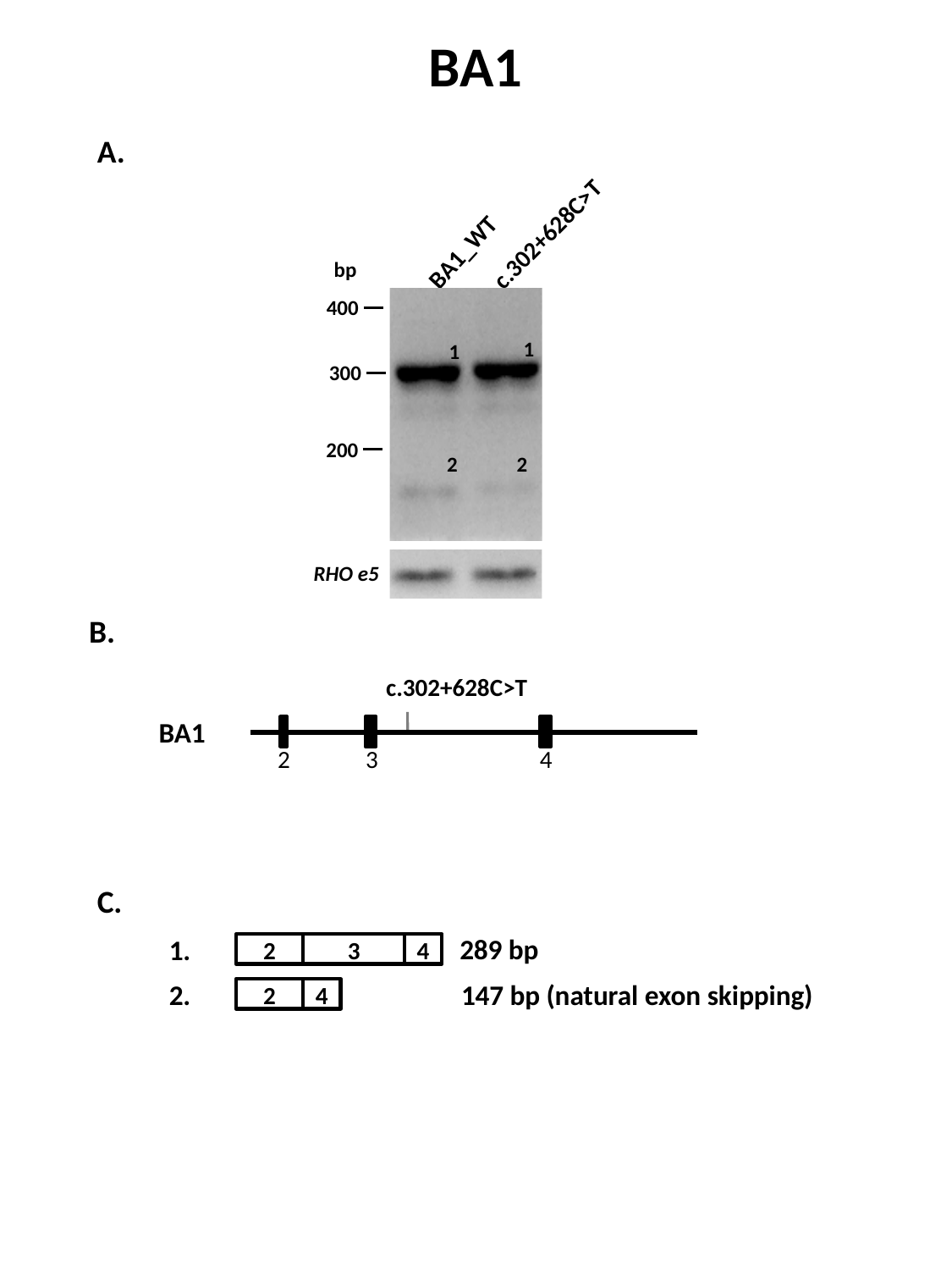

BA1
A.
c.302+628C>T
BA1_WT
bp
400
1
1
300
200
2
2
RHO e5
B.
c.302+628C>T
BA1
2
3
4
C.
289 bp
1.
2
3
4
2.
147 bp (natural exon skipping)
2
4

### Slide 3
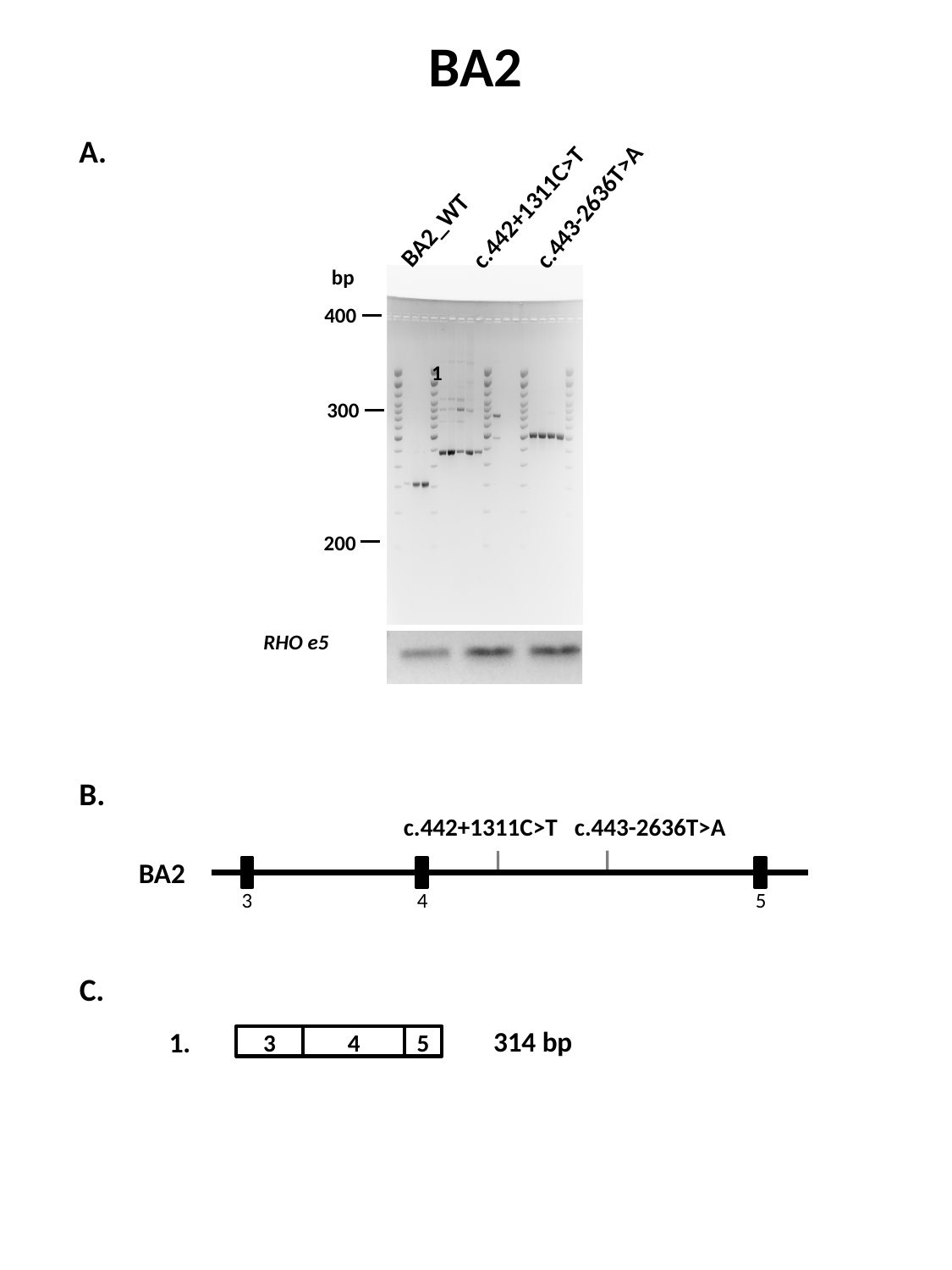

BA2
c.442+1311C>T
c.443-2636T>A
BA2_WT
bp
400
300
200
A.
1
RHO e5
B.
c.442+1311C>T
c.443-2636T>A
BA2
3
4
5
C.
314 bp
1.
3
4
5

### Slide 4
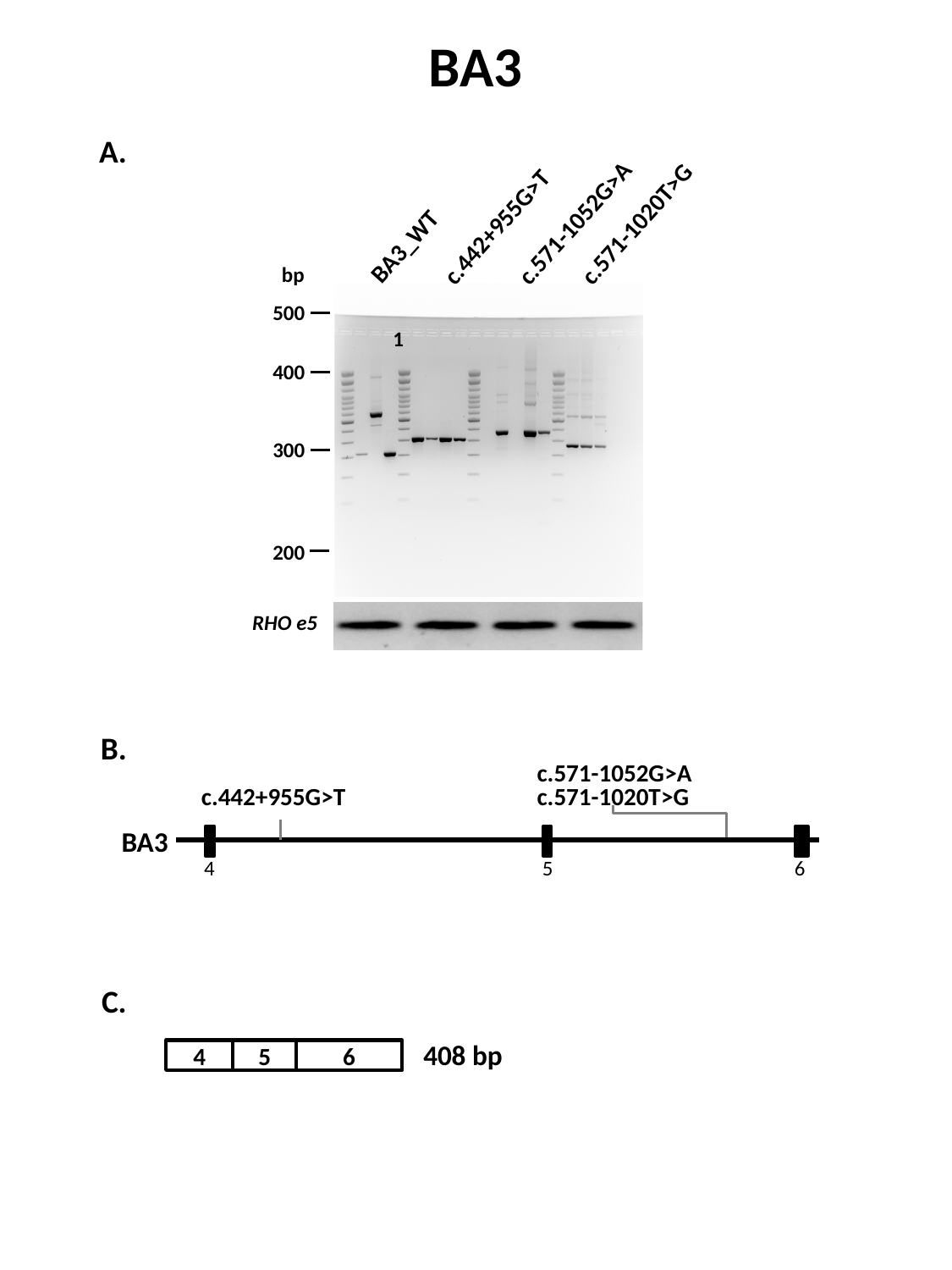

BA3
c.442+955G>T
c.571-1052G>A
c.571-1020T>G
BA3_WT
bp
500
400
300
200
A.
1
RHO e5
B.
c.571-1052G>A
c.442+955G>T
c.571-1020T>G
BA3
4
5
6
C.
408 bp
4
5
6

### Slide 5
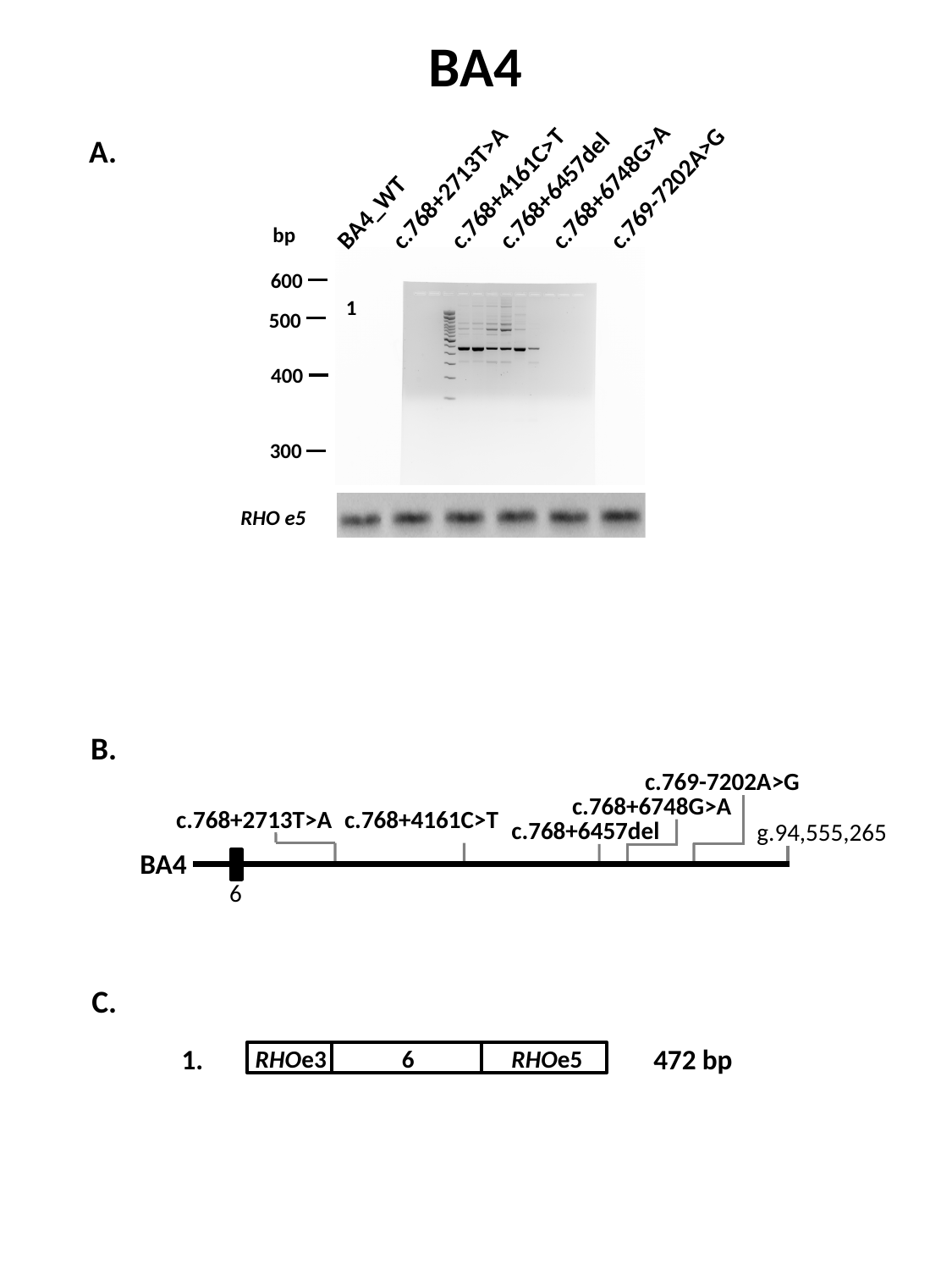

BA4
c.768+2713T>A
c.768+6748G>A
c.769-7202A>G
c.768+4161C>T
c.768+6457del
BA4_WT
bp
600
500
400
300
1
A.
RHO e5
B.
c.769-7202A>G
c.768+6748G>A
c.768+2713T>A
c.768+4161C>T
c.768+6457del
g.94,555,265
BA4
6
C.
1.
472 bp
RHOe3
RHOe5
6

### Slide 6
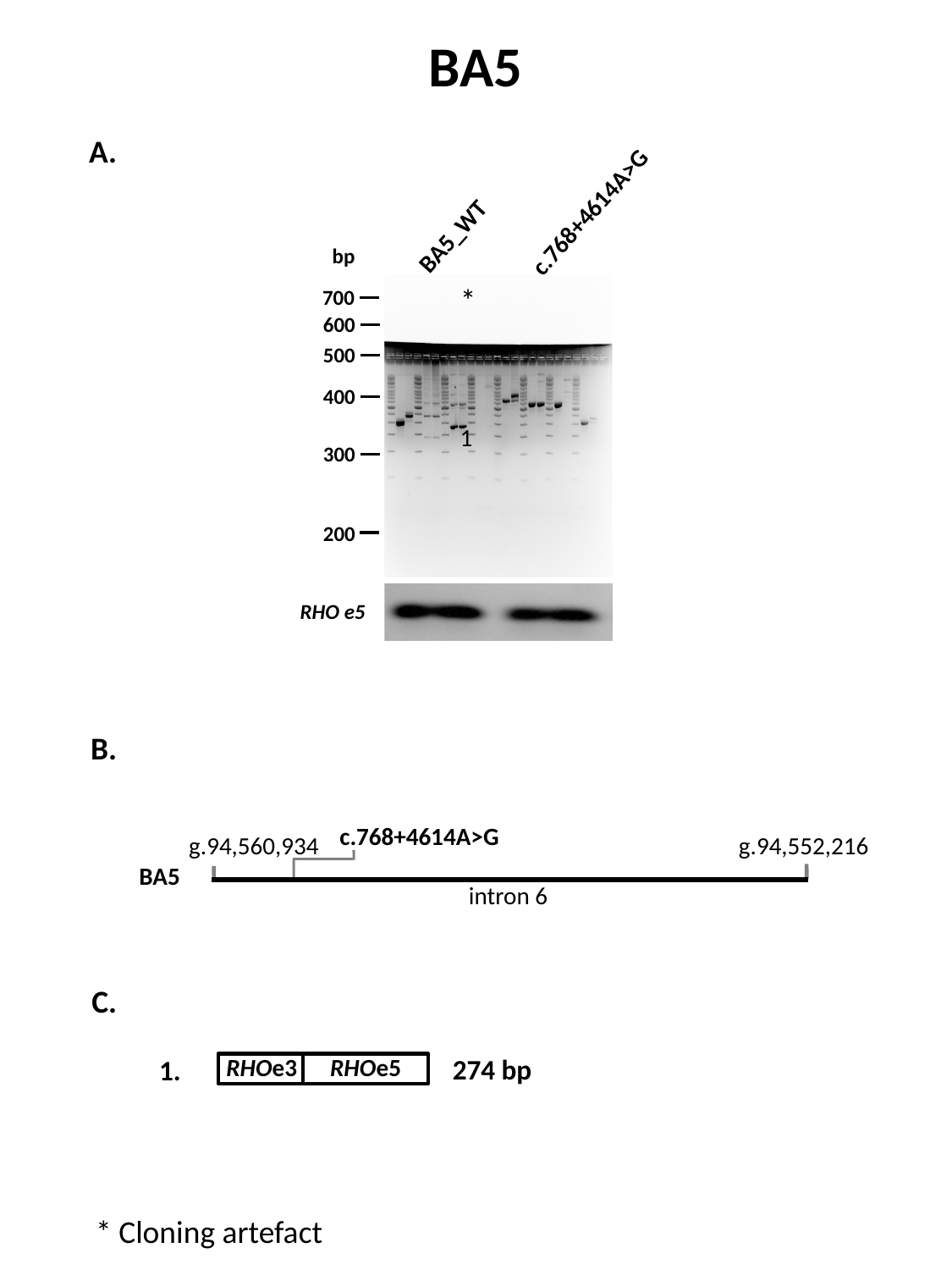

BA5
c.768+4614A>G
BA5_WT
bp
700
600
500
400
300
200
*
*
1
A.
RHO e5
B.
c.768+4614A>G
g.94,560,934
g.94,552,216
BA5
intron 6
C.
274 bp
RHOe3
RHOe5
1.
* Cloning artefact

### Slide 7
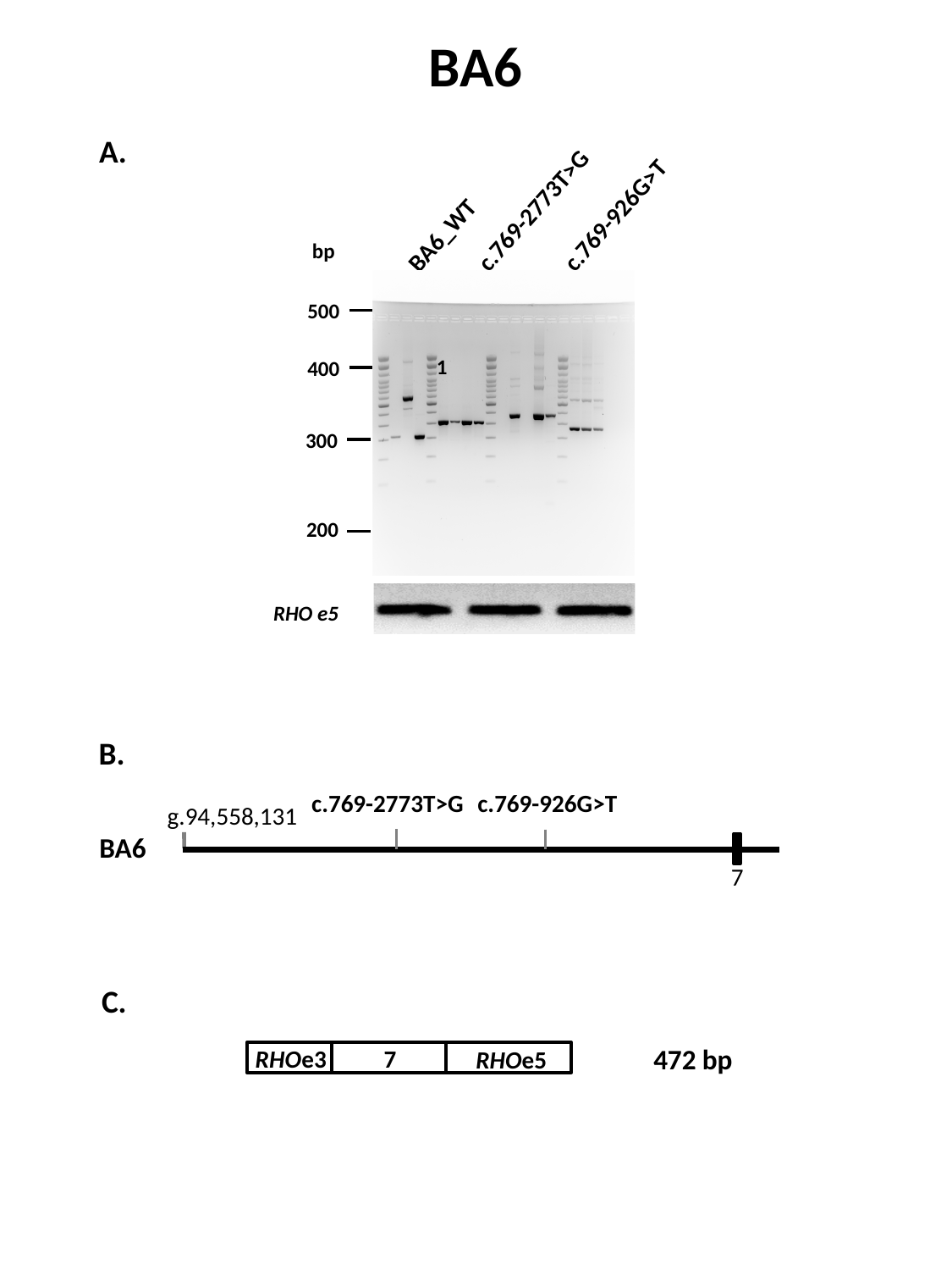

BA6
c.769-2773T>G
c.769-926G>T
BA6_WT
bp
500
400
300
200
A.
1
RHO e5
B.
c.769-2773T>G
c.769-926G>T
g.94,558,131
BA6
7
C.
472 bp
RHOe3
7
RHOe5

### Slide 8
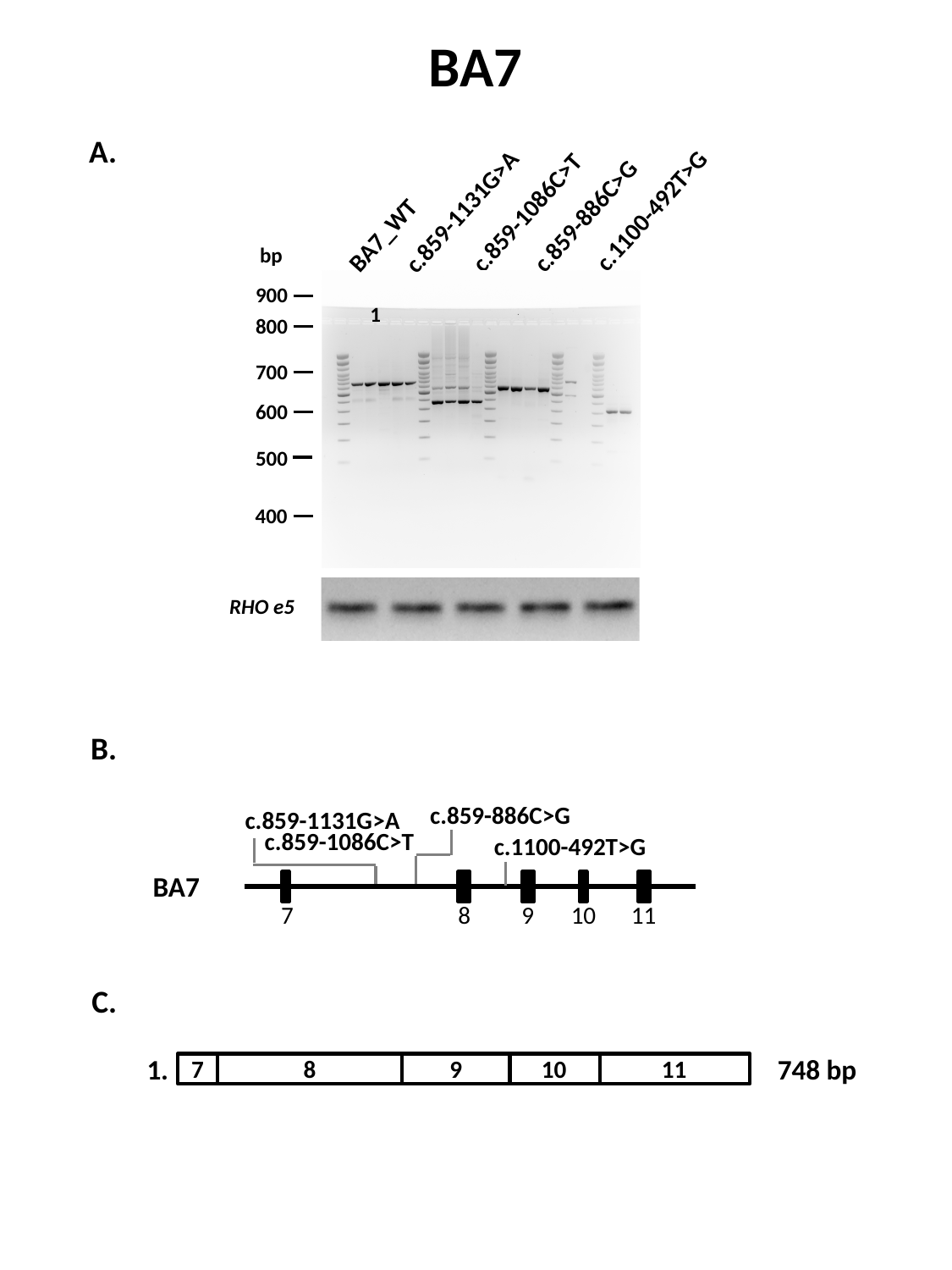

BA7
c.1100-492T>G
c.859-1086C>T
c.859-1131G>A
BA7_WT
c.859-886C>G
bp
900
800
700
600
500
400
A.
1
RHO e5
B.
c.859-886C>G
c.859-1131G>A
c.859-1086C>T
c.1100-492T>G
BA7
7
8
9
10
11
C.
748 bp
1.
10
7
8
9
11

### Slide 9
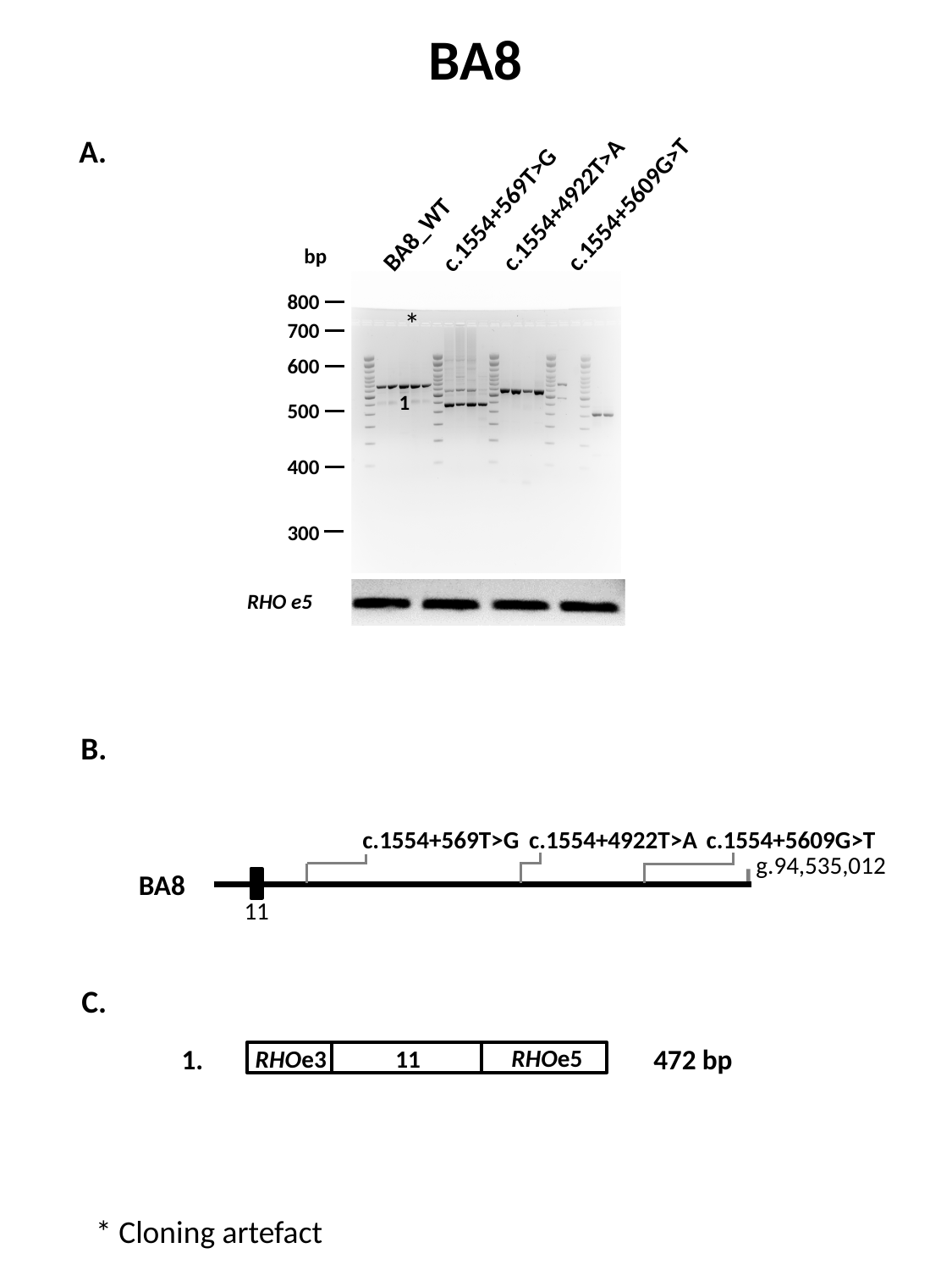

BA8
c.1554+4922T>A
c.1554+5609G>T
c.1554+569T>G
BA8_WT
bp
800
700
600
500
400
300
A.
*
1
RHO e5
B.
c.1554+569T>G
c.1554+4922T>A
c.1554+5609G>T
g.94,535,012
BA8
11
C.
1.
472 bp
RHOe5
RHOe3
11
* Cloning artefact

### Slide 10
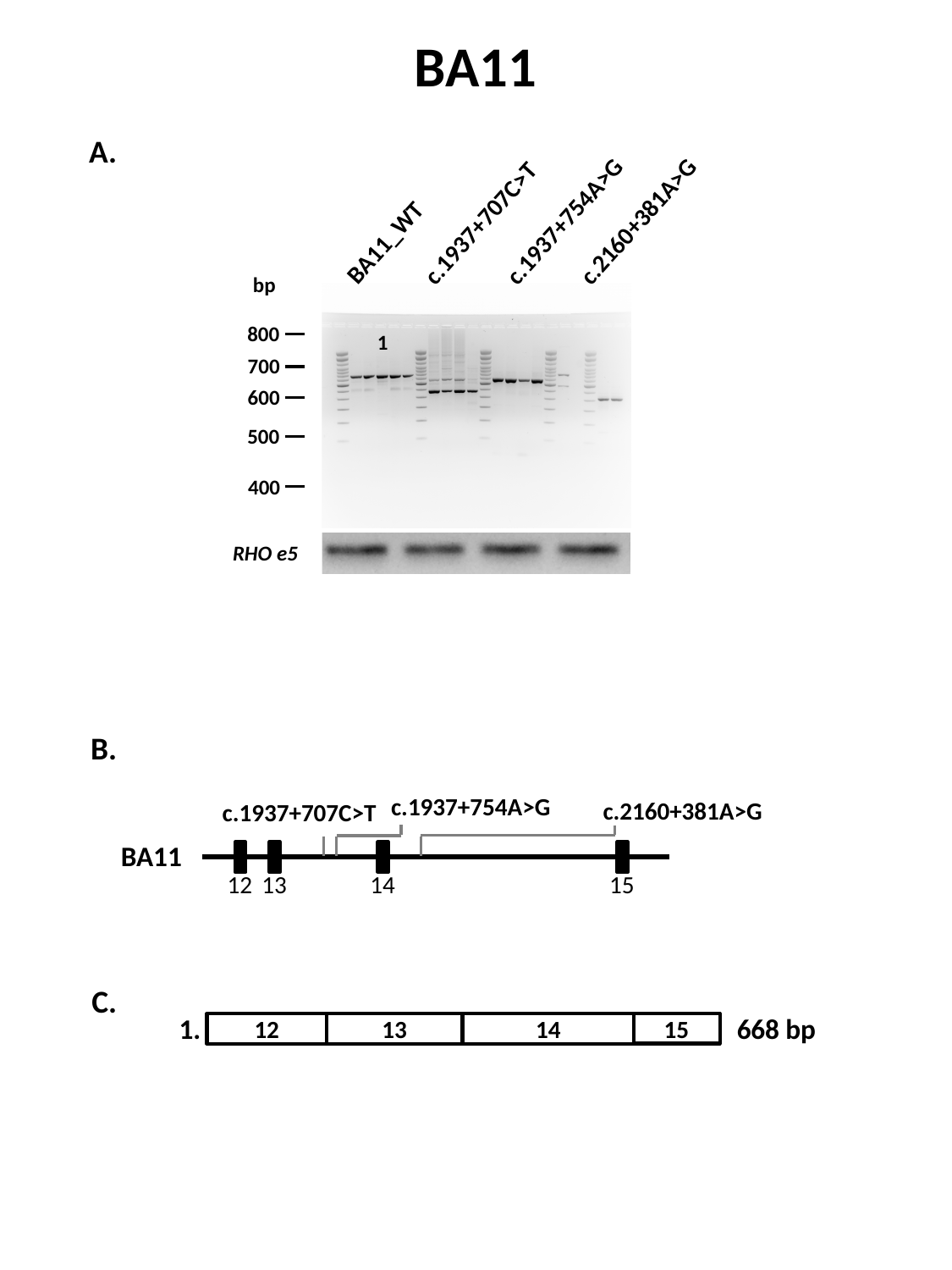

BA11
c.1937+754A>G
c.1937+707C>T
c.2160+381A>G
BA11_WT
bp
800
700
600
500
400
A.
1
RHO e5
B.
c.1937+754A>G
c.2160+381A>G
c.1937+707C>T
BA11
12
13
14
15
C.
668 bp
1.
12
13
14
15

### Slide 11
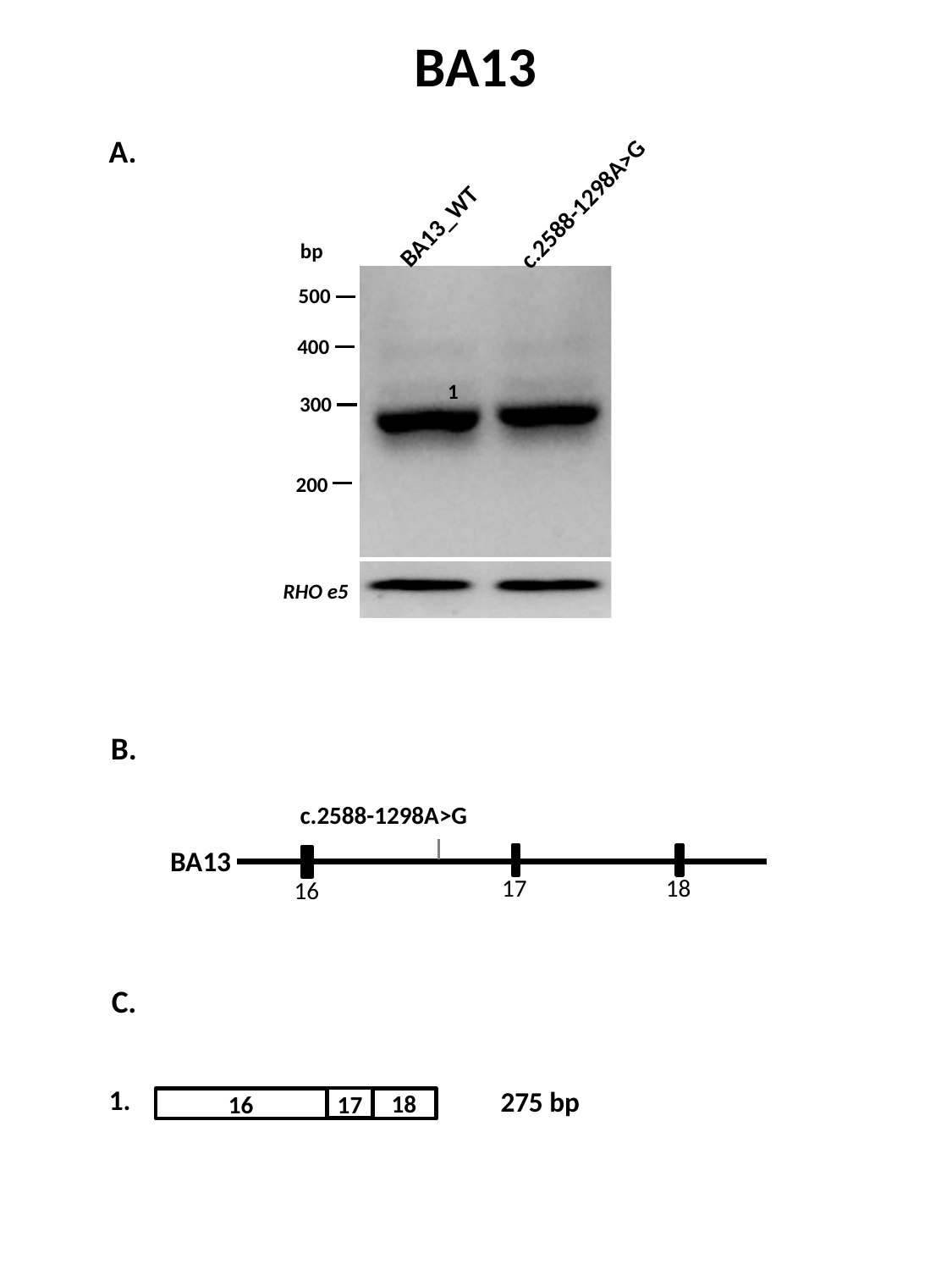

BA13
c.2588-1298A>G
BA13_WT
bp
500
400
300
200
A.
1
RHO e5
B.
c.2588-1298A>G
BA13
17
18
16
C.
1.
275 bp
18
17
16

### Slide 12
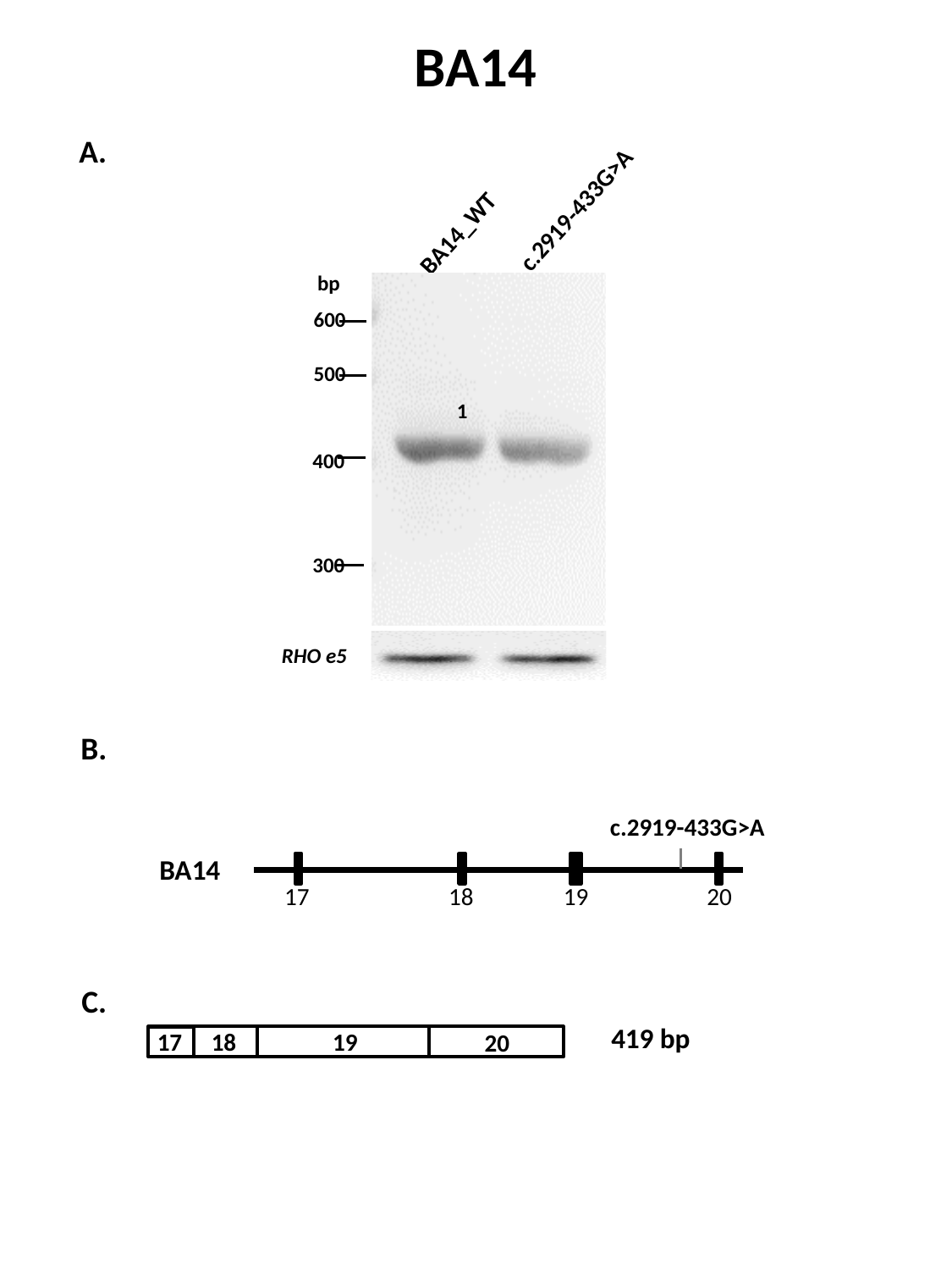

BA14
c.2919-433G>A
BA14_WT
bp
600
500
400
300
A.
1
RHO e5
B.
c.2919-433G>A
BA14
17
18
19
20
C.
419 bp
17
18
19
20

### Slide 13
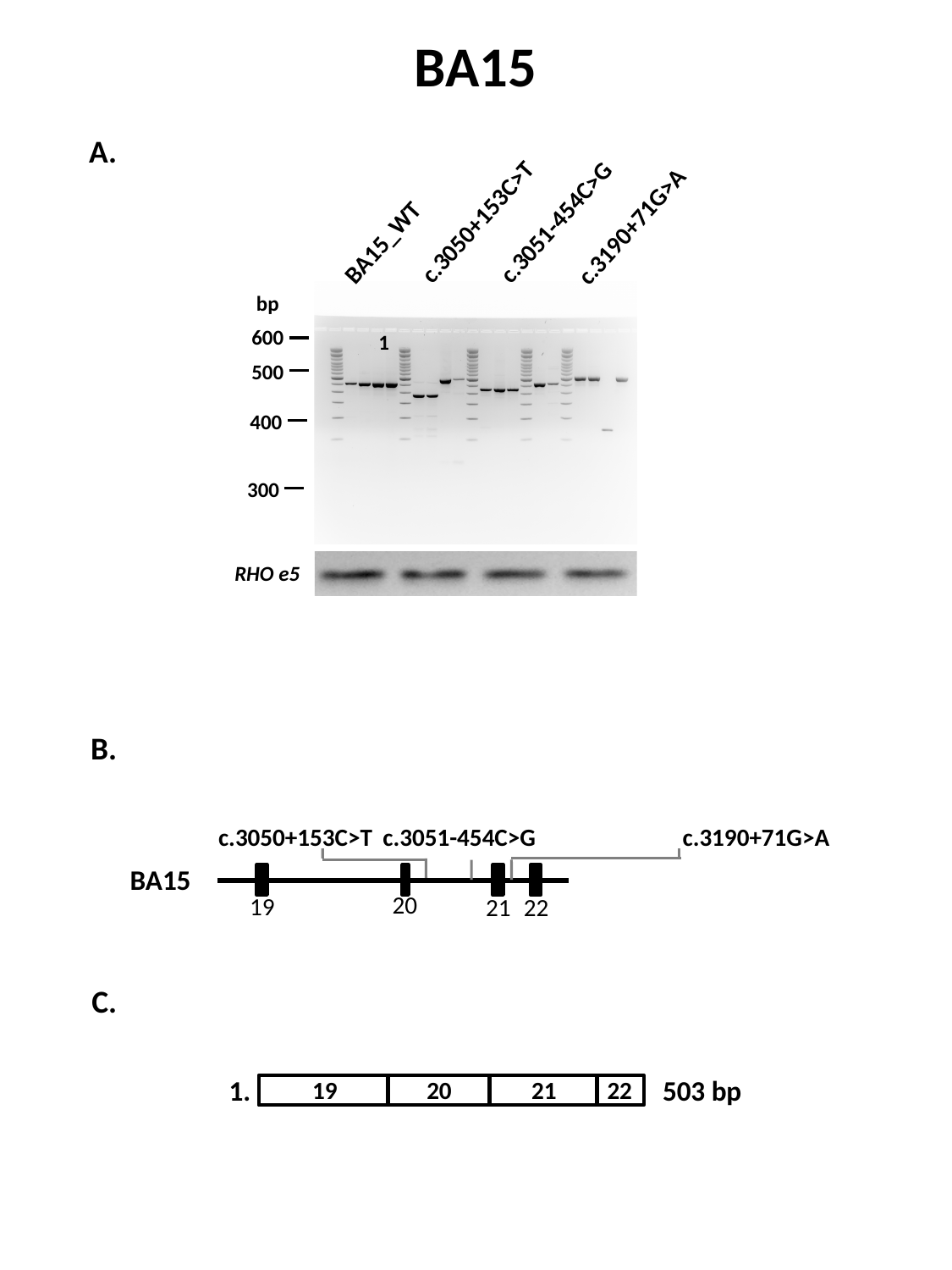

BA15
c.3051-454C>G
c.3190+71G>A
c.3050+153C>T
BA15_WT
bp
600
500
400
300
A.
1
RHO e5
B.
c.3050+153C>T
c.3051-454C>G
c.3190+71G>A
BA15
20
19
21
22
C.
1.
19
20
21
22
503 bp

### Slide 14
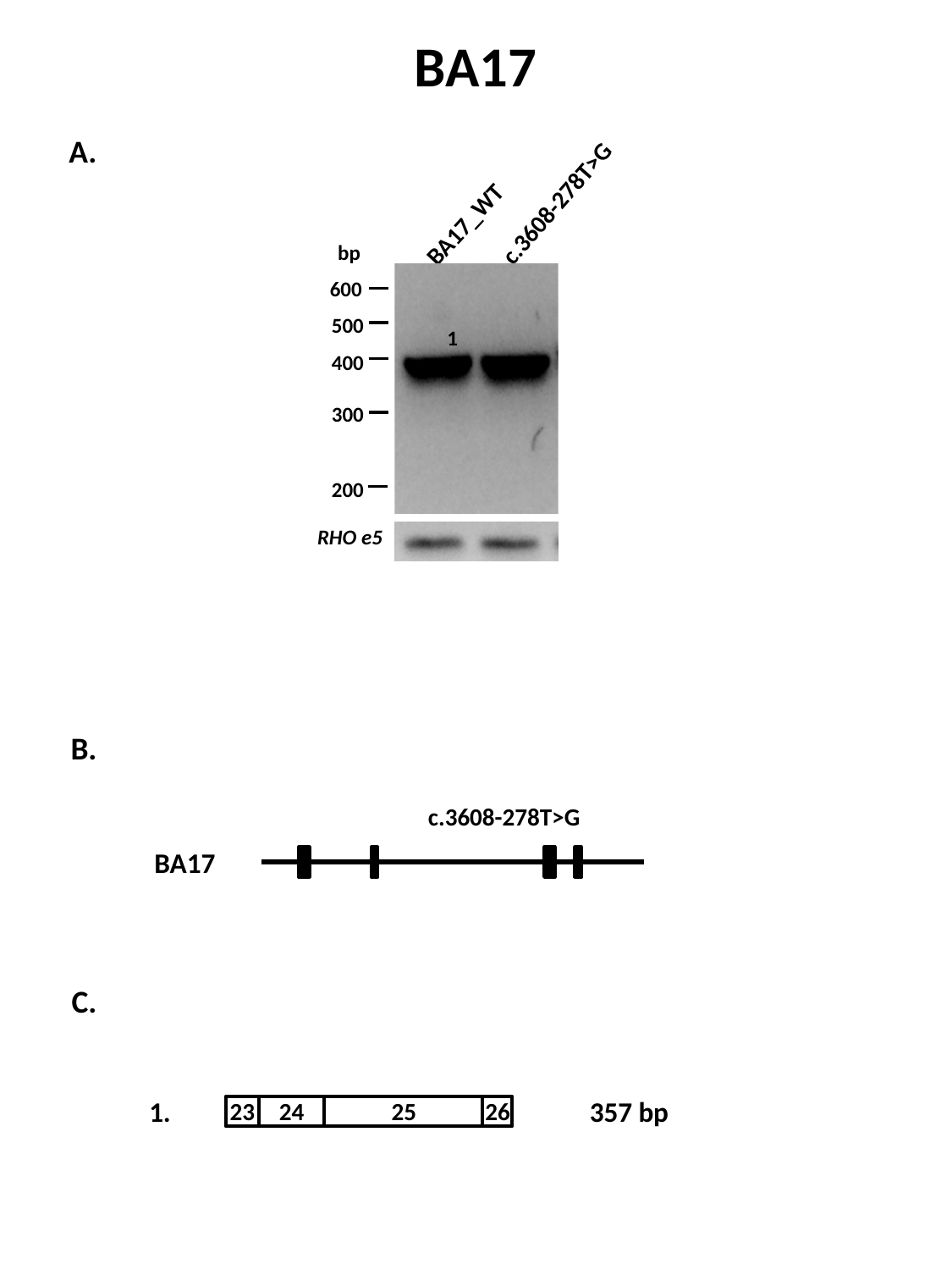

BA17
c.3608-278T>G
BA17_WT
bp
600
500
400
300
200
A.
1
RHO e5
B.
c.3608-278T>G
BA17
C.
1.
23
24
25
26
357 bp

### Slide 15
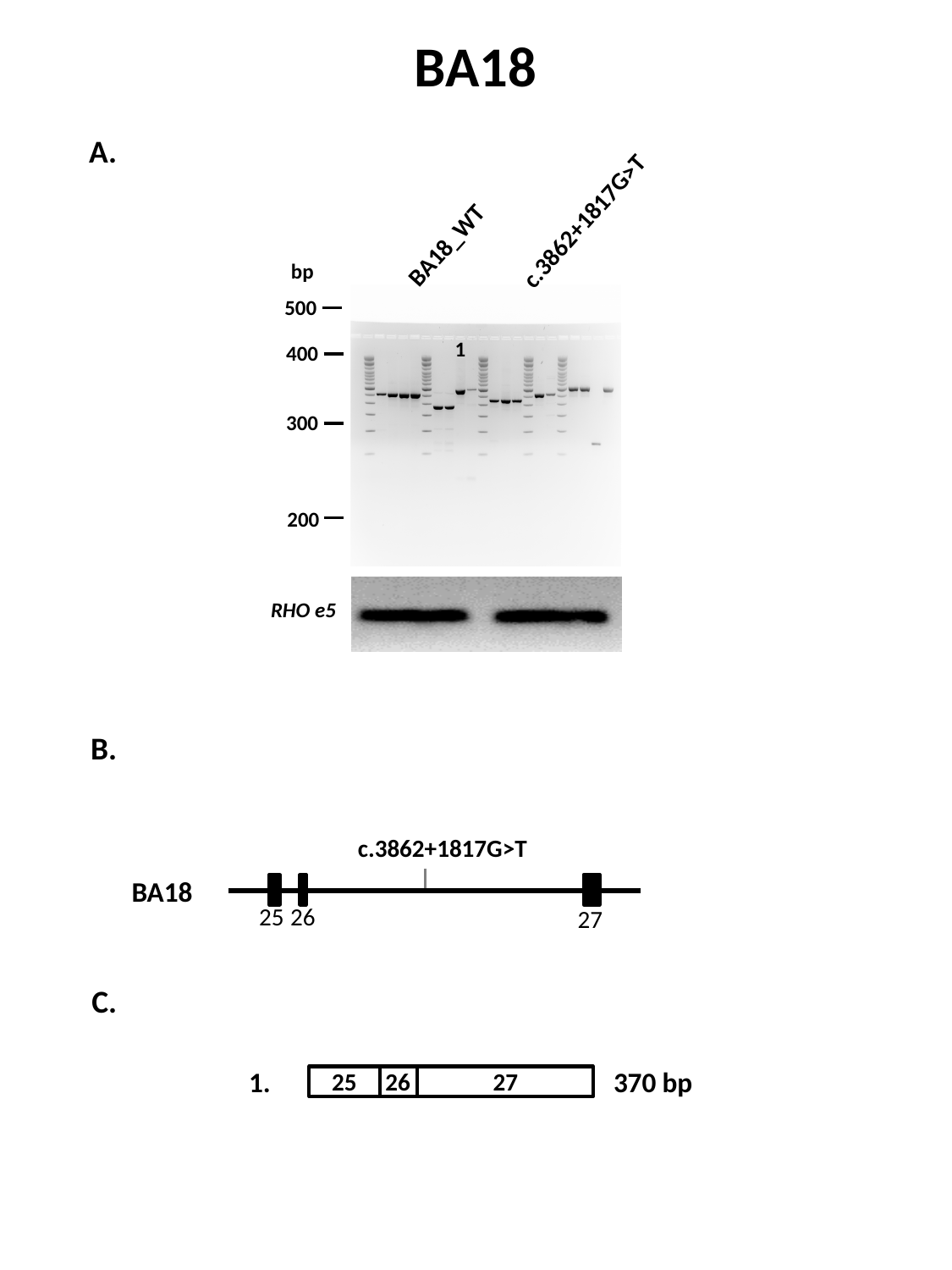

BA18
c.3862+1817G>T
BA18_WT
bp
500
400
300
200
A.
1
RHO e5
B.
c.3862+1817G>T
BA18
25
26
27
C.
1.
25
26
27
370 bp

### Slide 16
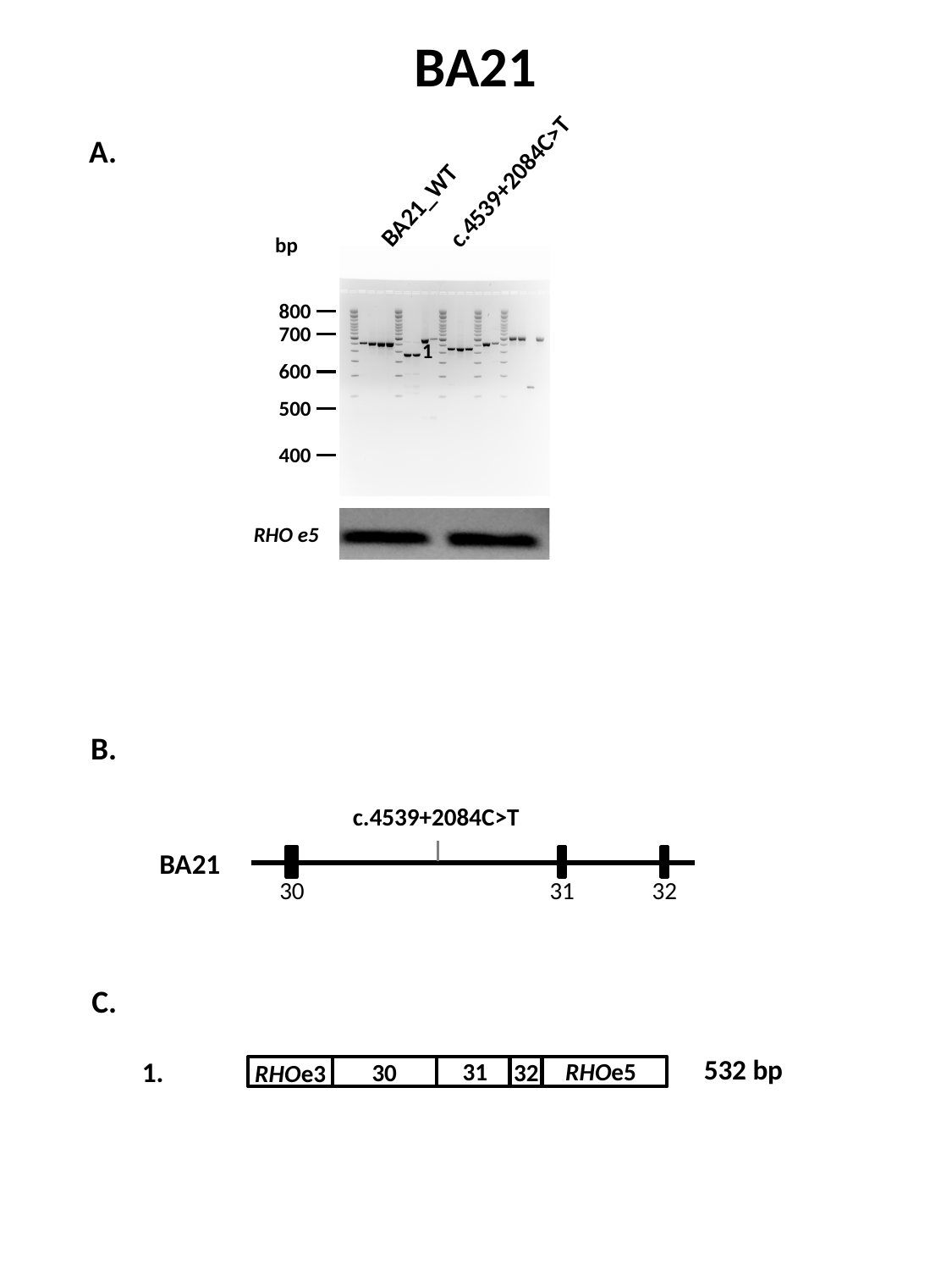

BA21
A.
c.4539+2084C>T
BA21_WT
bp
800
700
1
600
500
400
RHO e5
B.
c.4539+2084C>T
BA21
30
31
32
C.
532 bp
1.
31
RHOe5
30
32
RHOe3

### Slide 17
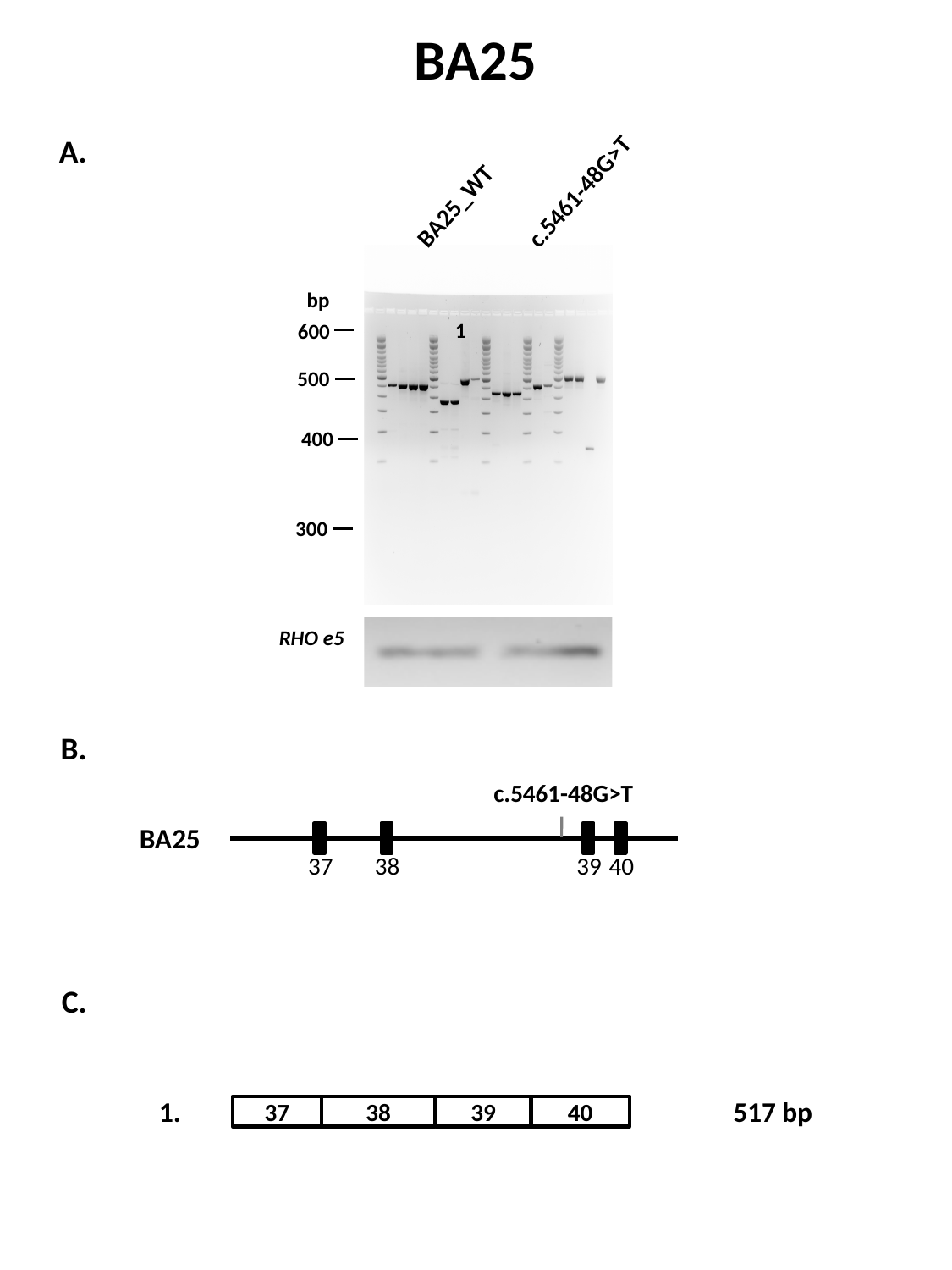

BA25
BA25_WT
c.5461-48G>T
bp
600
500
400
300
A.
1
RHO e5
B.
c.5461-48G>T
BA25
40
37
38
39
C.
1.
37
38
39
40
517 bp

### Slide 18
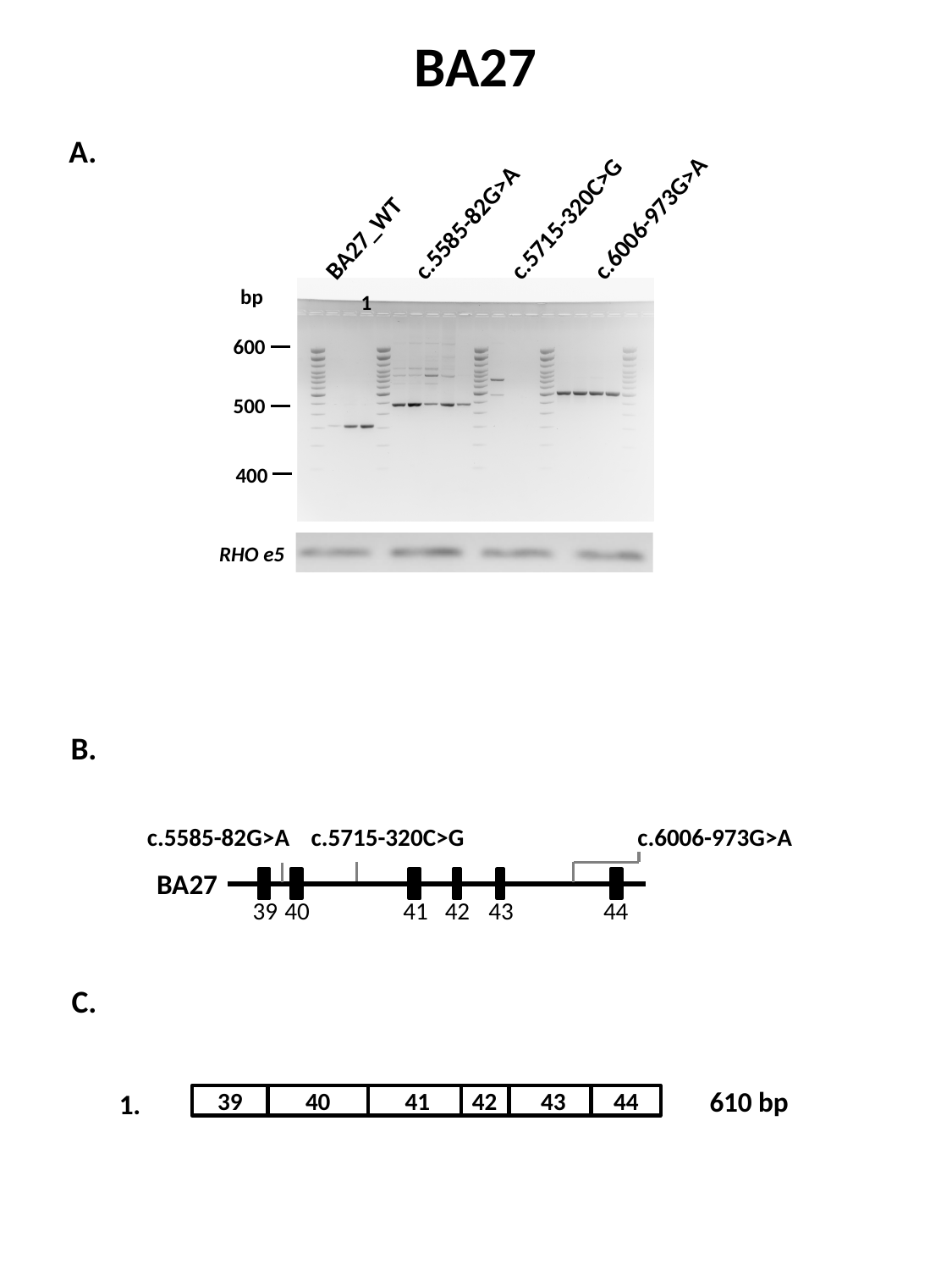

BA27
c.5715-320C>G
c.6006-973G>A
BA27_WT
c.5585-82G>A
bp
600
500
400
A.
1
RHO e5
B.
c.5585-82G>A
c.5715-320C>G
c.6006-973G>A
BA27
40
41
39
42
43
44
C.
610 bp
42
39
40
41
43
44
1.

### Slide 19
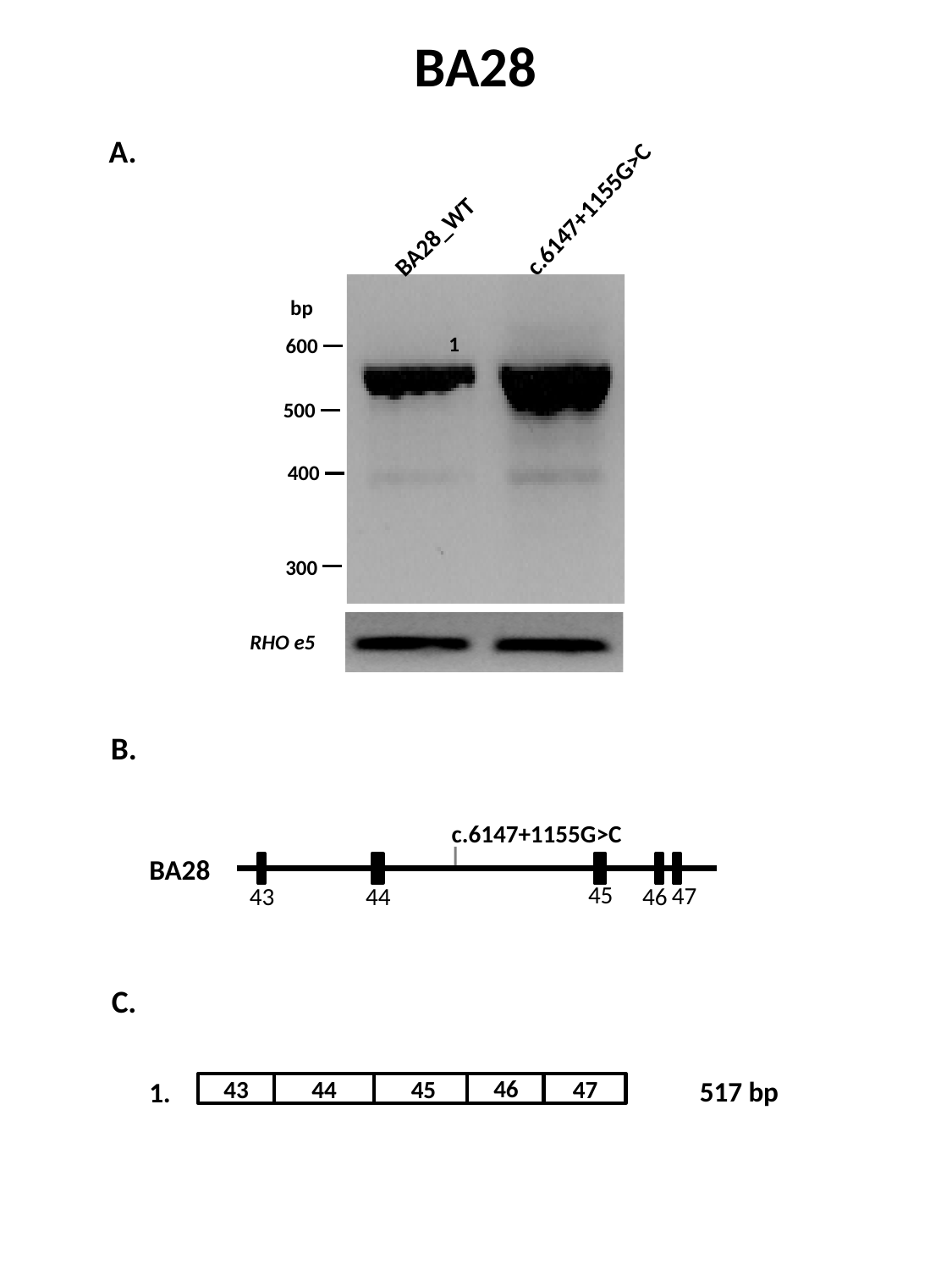

BA28
c.6147+1155G>C
BA28_WT
bp
600
500
400
300
A.
1
RHO e5
B.
c.6147+1155G>C
BA28
45
47
43
44
46
C.
46
43
44
45
47
517 bp
1.

### Slide 20
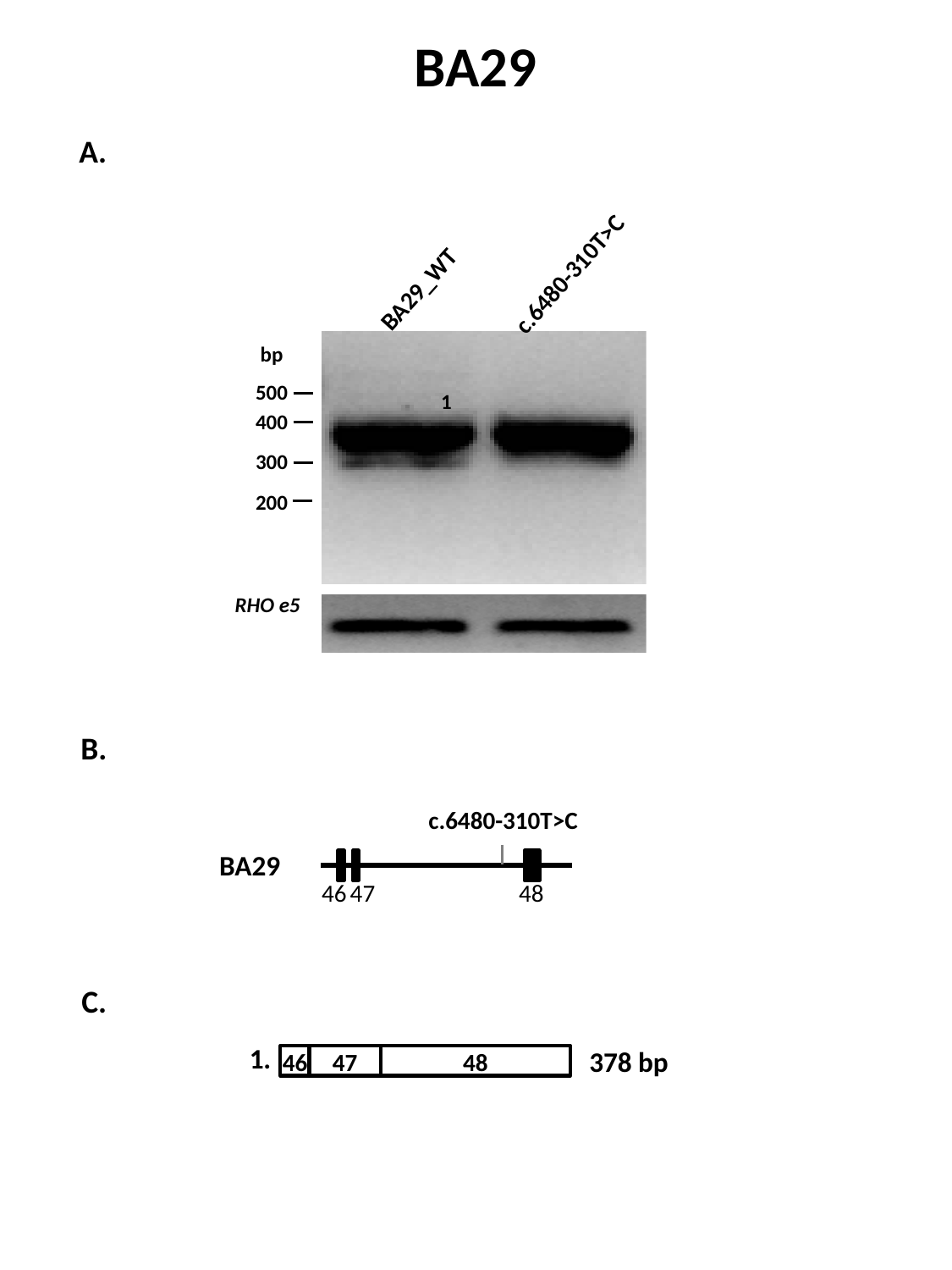

BA29
A.
c.6480-310T>C
BA29_WT
bp
500
400
300
200
1
RHO e5
B.
c.6480-310T>C
BA29
48
46
47
C.
1.
46
47
48
378 bp

### Slide 21
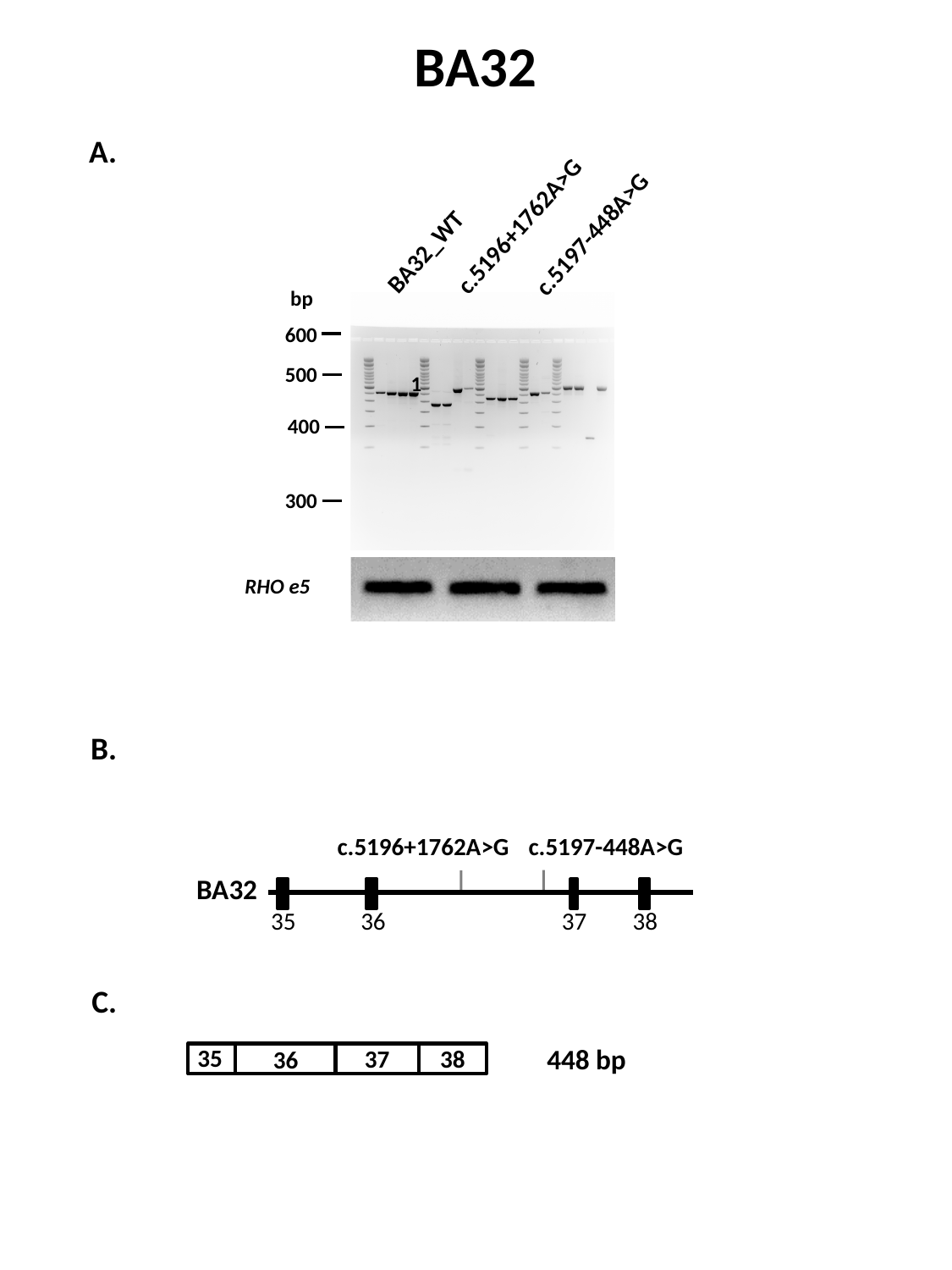

BA32
c.5197-448A>G
c.5196+1762A>G
BA32_WT
bp
600
500
400
300
A.
1
RHO e5
B.
c.5196+1762A>G
c.5197-448A>G
BA32
35
36
37
38
C.
448 bp
35
38
37
36

### Slide 22
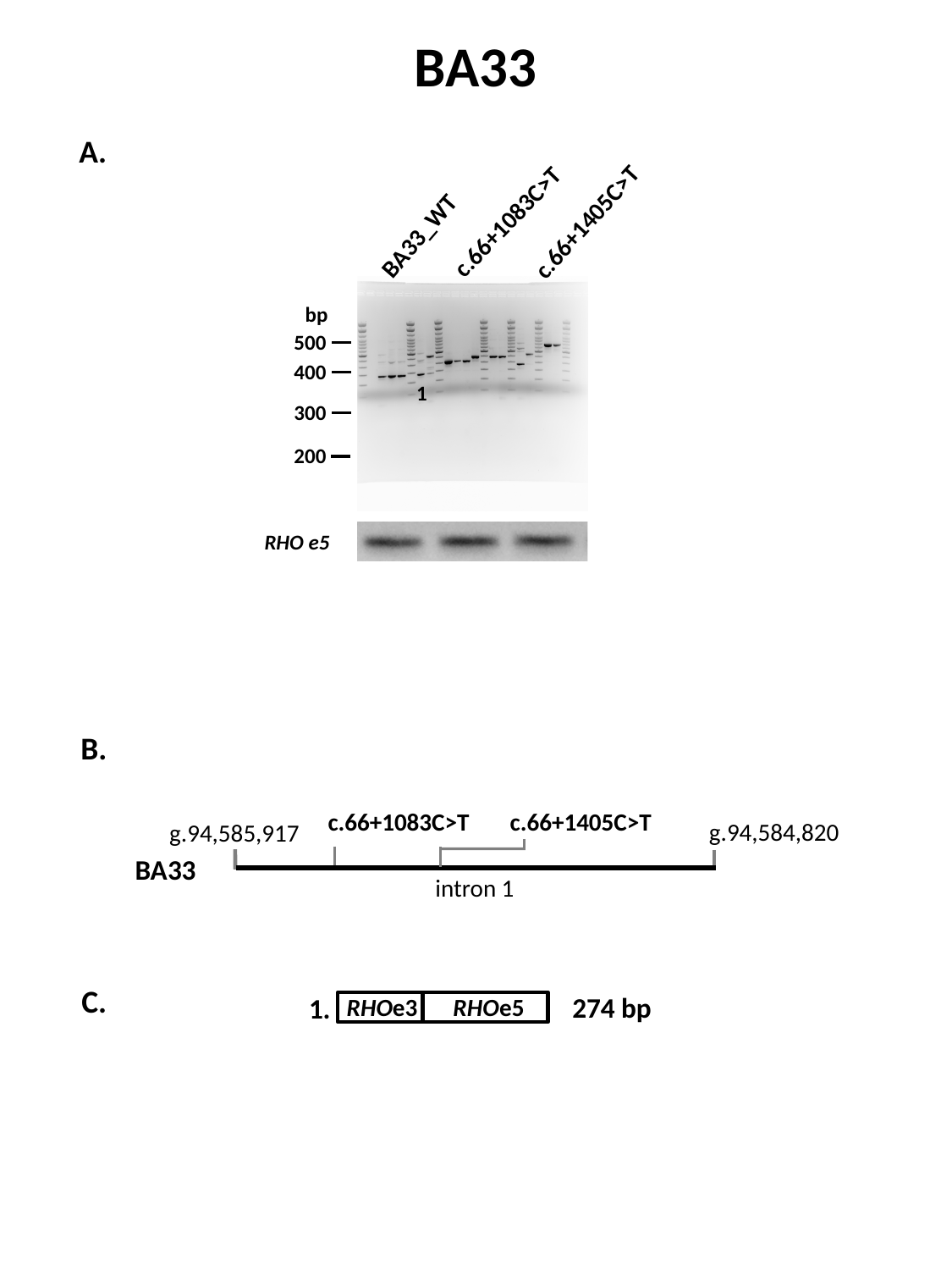

BA33
A.
c.66+1405C>T
c.66+1083C>T
BA33_WT
bp
500
400
300
200
1
RHO e5
B.
c.66+1083C>T
c.66+1405C>T
g.94,585,917
g.94,584,820
BA33
intron 1
C.
274 bp
1.
RHOe5
RHOe3

### Slide 23
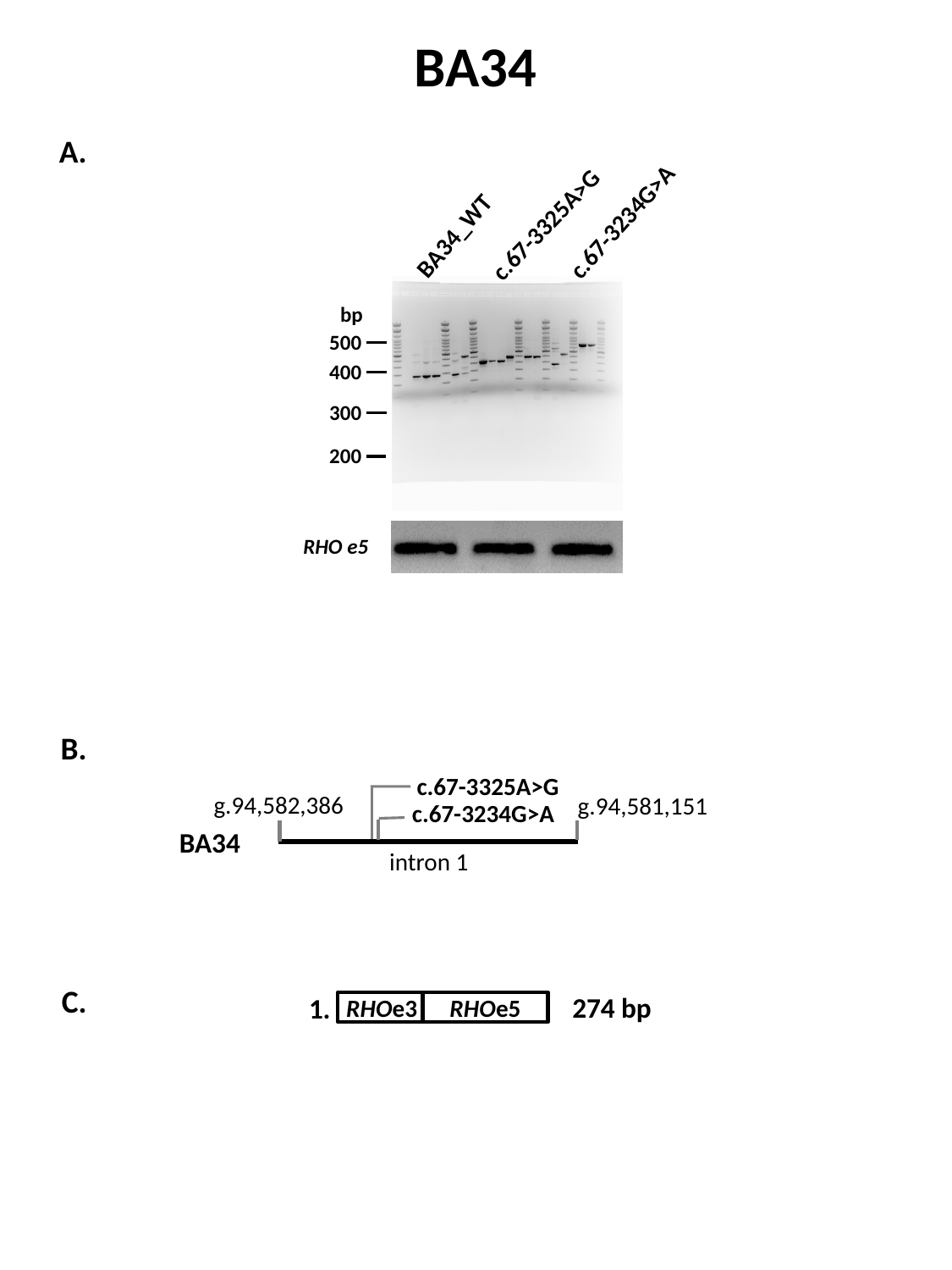

BA34
c.67-3325A>G
c.67-3234G>A
BA34_WT
bp
500
400
300
200
RHO e5
A.
1
B.
c.67-3325A>G
g.94,582,386
g.94,581,151
c.67-3234G>A
BA34
intron 1
C.
274 bp
1.
RHOe5
RHOe3
