## Supplemental Fig. S6. Deep-intronic variants Sanger sequencing results for "Resolving the dark matter of *ABCA4* for 1,054 Stargardt disease probands through integrated genomics and transcriptomics"

### Slide 1
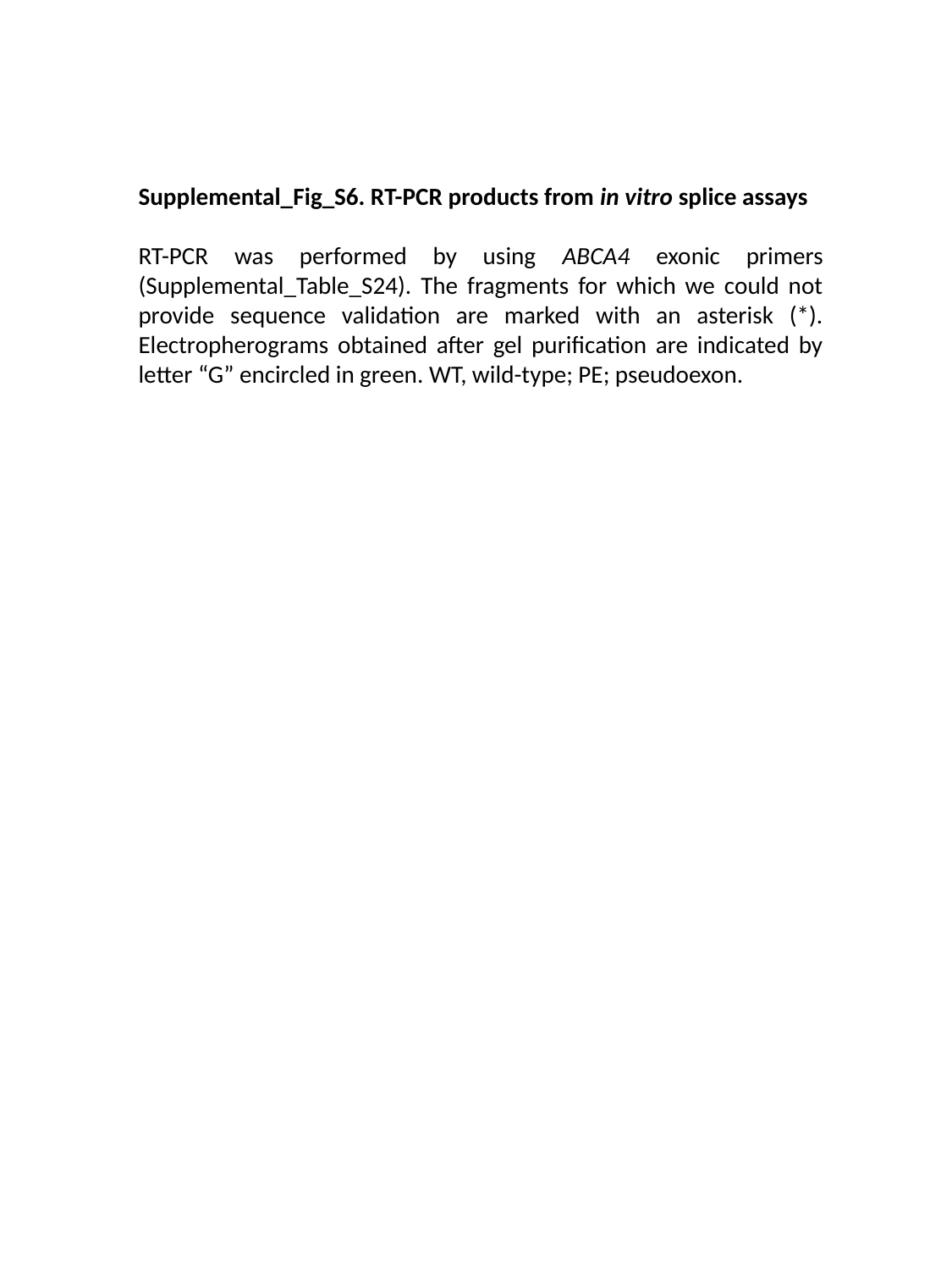

Supplemental_Fig_S6. RT-PCR products from in vitro splice assays
RT-PCR was performed by using ABCA4 exonic primers (Supplemental_Table_S24). The fragments for which we could not provide sequence validation are marked with an asterisk (*). Electropherograms obtained after gel purification are indicated by letter “G” encircled in green. WT, wild-type; PE; pseudoexon.

### Slide 2
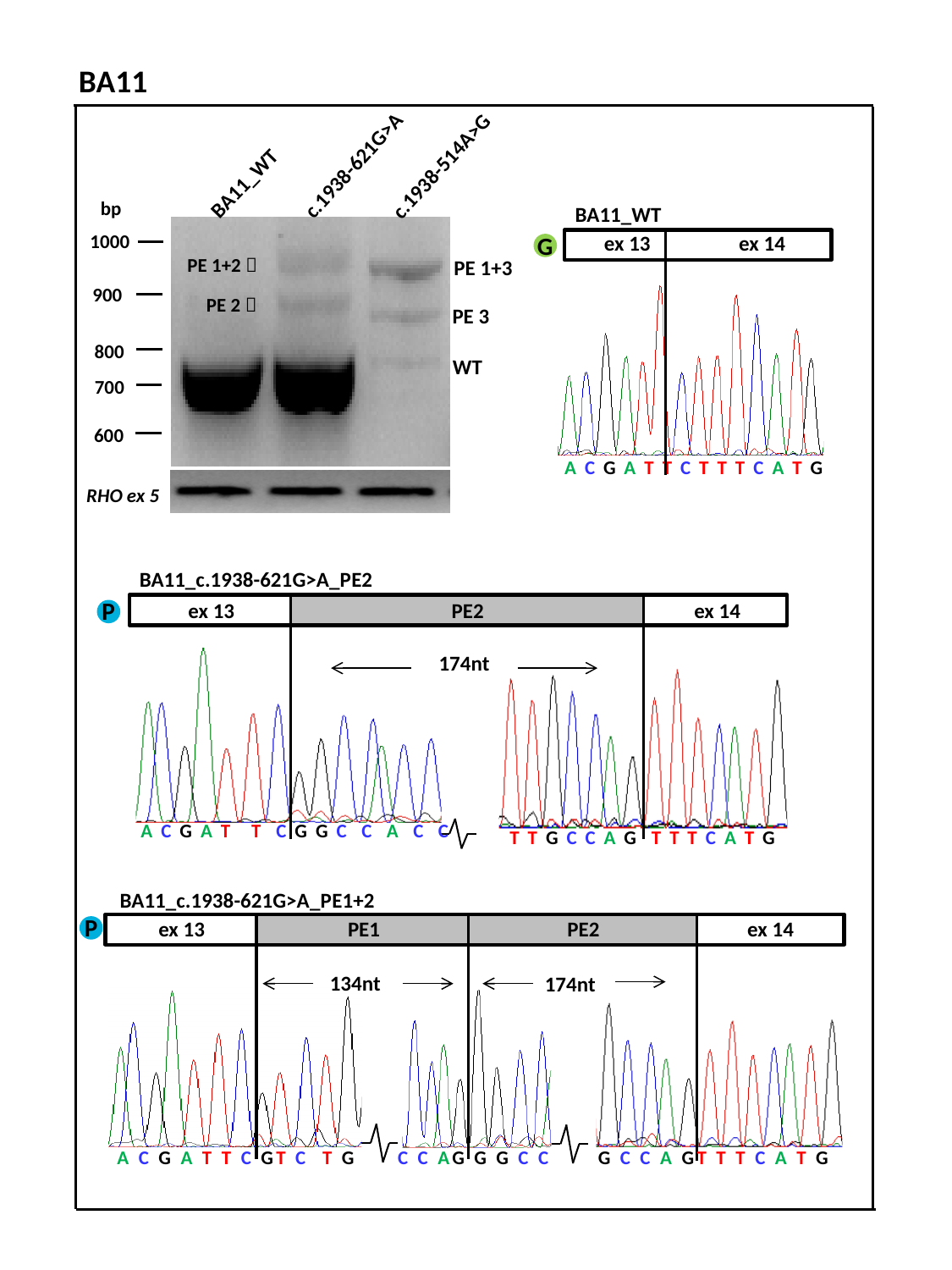

BA11
c.1938-621G>A
c.1938-514A>G
BA11_WT
bp
1000
900
800
700
600
RHO ex 5
PE 1+2 🡪
PE 1+3
PE 2 🡪
PE 3
WT
ex 3
BA11_WT
ex 13
ex 14
ex 4
A C G A T T C T T T C A T G
G
BA11_c.1938-621G>A_PE2
P
 ex 13
PE2
 ex 14
174nt
A C G A T T C G G C C A C C
T T G C C A G T T T C A T G
BA11_c.1938-621G>A_PE1+2
 ex 13
PE1
PE2
 ex 14
134nt
174nt
A C G A T T C GT C T G
C C AG G G C C
G C C A GT T T C A T G
P

### Slide 3
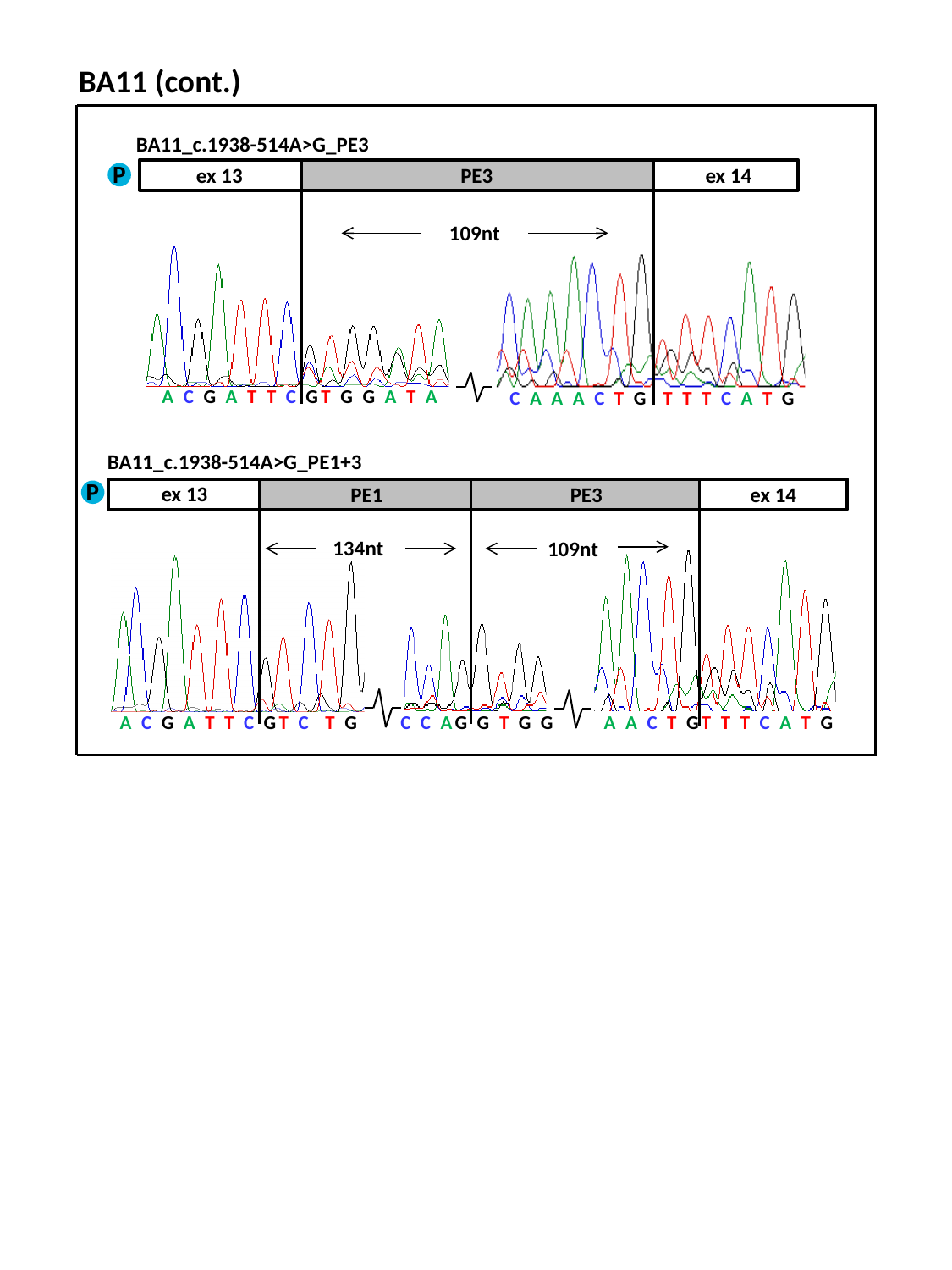

BA11 (cont.)
BA11_c.1938-514A>G_PE3
ex 3
P
ex 13
PE3
 ex 14
109nt
A C G A T T C GT G G A T A
C A A A C T G T T T C A T G
BA11_c.1938-514A>G_PE1+3
 ex 13
PE1
PE3
 ex 14
134nt
109nt
A C G A T T C GT C T G
C C AG G T G G
A A C T GT T T C A T G
P

### Slide 4
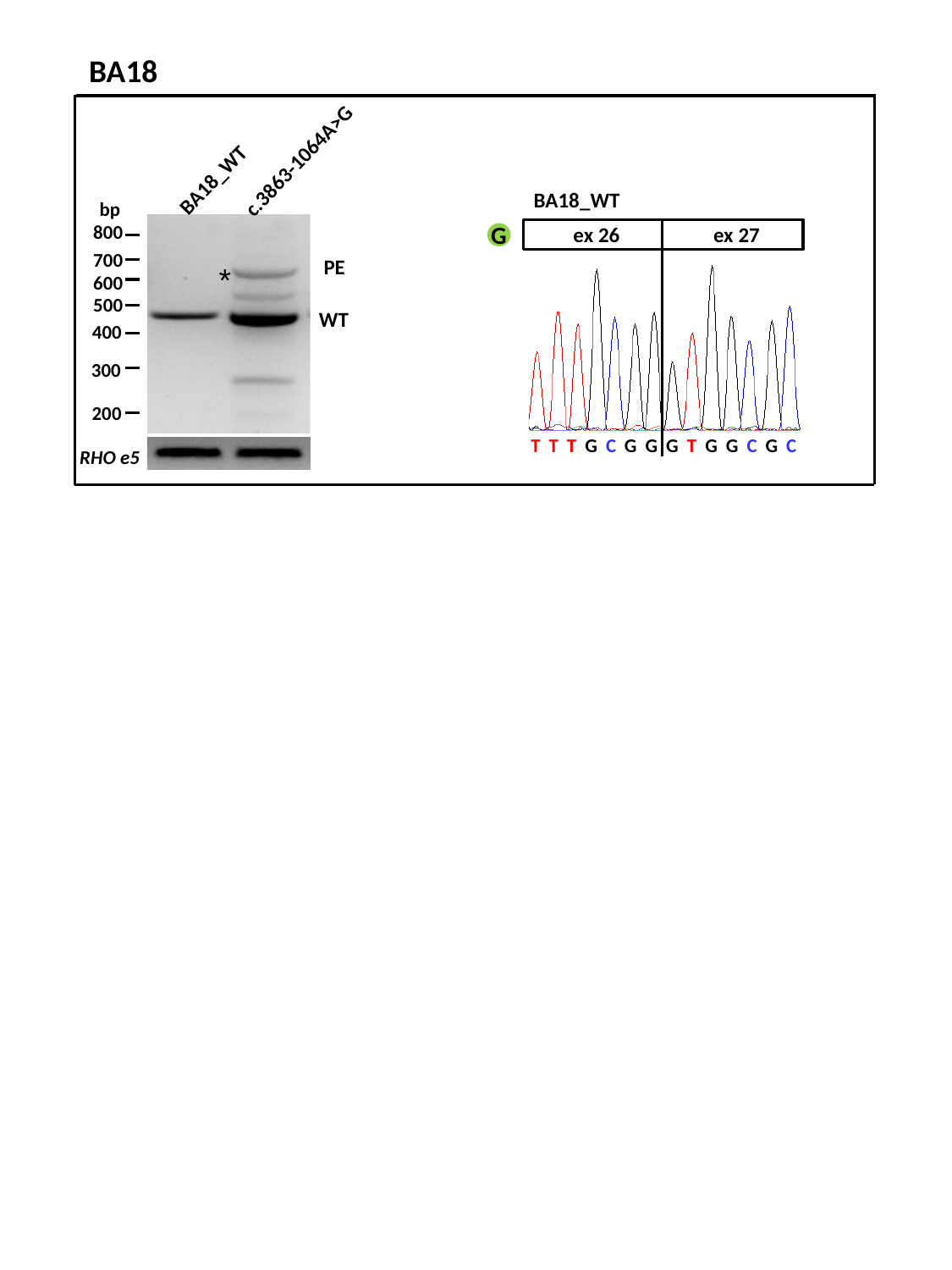

BA18
c.3863-1064A>G
BA18_WT
bp
800
700
600
500
400
300
200
RHO e5
BA18_WT
 ex 26
 ex 27
G
 PE
*
WT
T T T G C G G G T G G C G C

### Slide 5
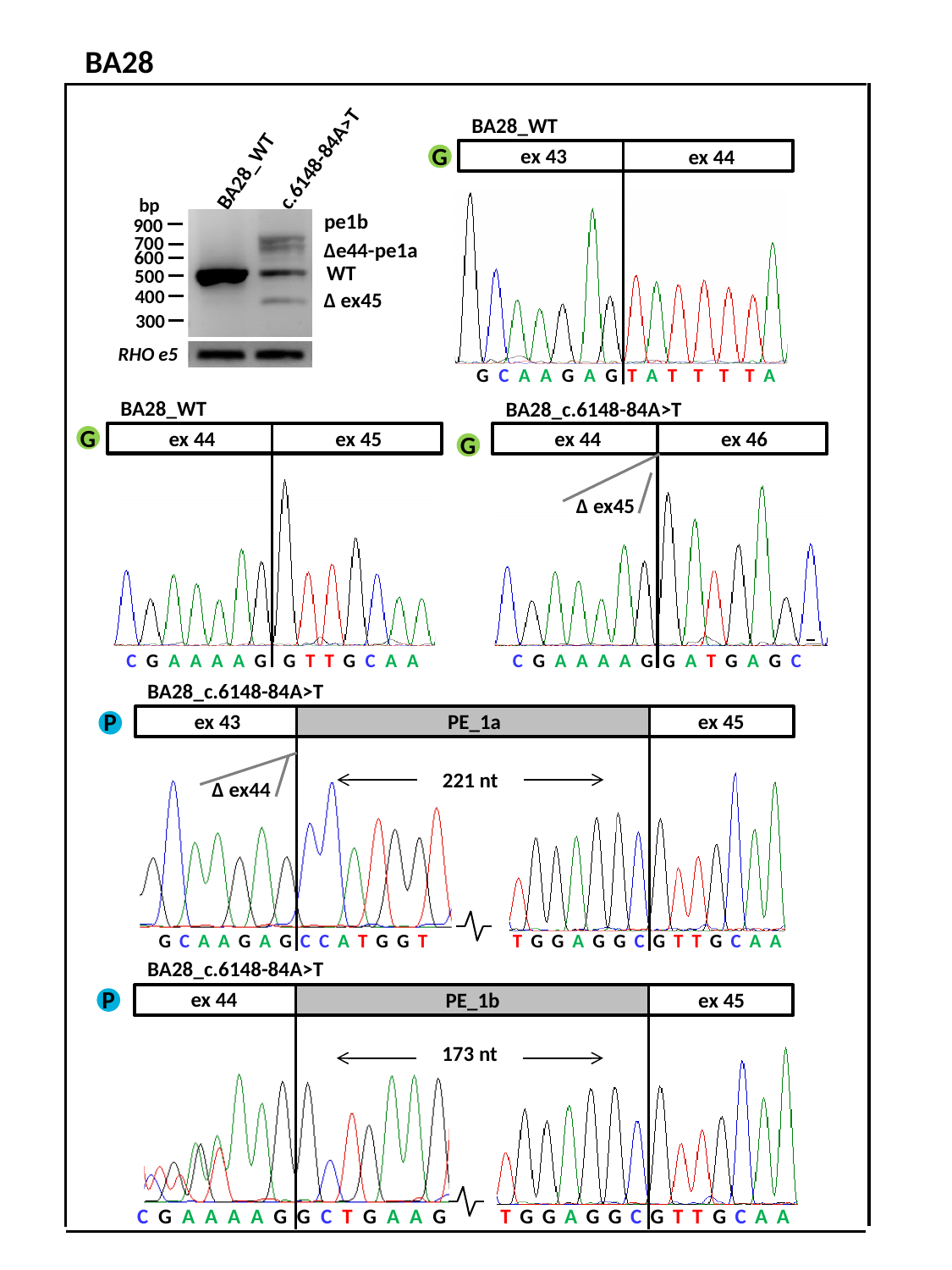

BA28
c.6148-84A>T
BA28_WT
bp
900
700
600
500
400
300
BA28_WT
 ex 43
 ex 44
G C A A G A G T A T T T T A
G
pe1b
∆e44-pe1a
WT
 ∆ ex45
RHO e5
BA28_WT
 ex 44
 ex 45
BA28_c.6148-84A>T
 ex 44
 ex 46
C G A A A A G G A T G A G C
∆ ex45
G
G
C G A A A A G G T T G C A A
BA28_c.6148-84A>T
P
PE_1a
 ex 43
ex 45
221 nt
∆ ex44
G C A A G A G C C A T G G T
T G G A G G C G T T G C A A
BA28_c.6148-84A>T
 ex 44
 ex 45
173 nt
C G A A A A G G C T G A A G
T G G A G G C G T T G C A A
P
PE_1b

### Slide 6
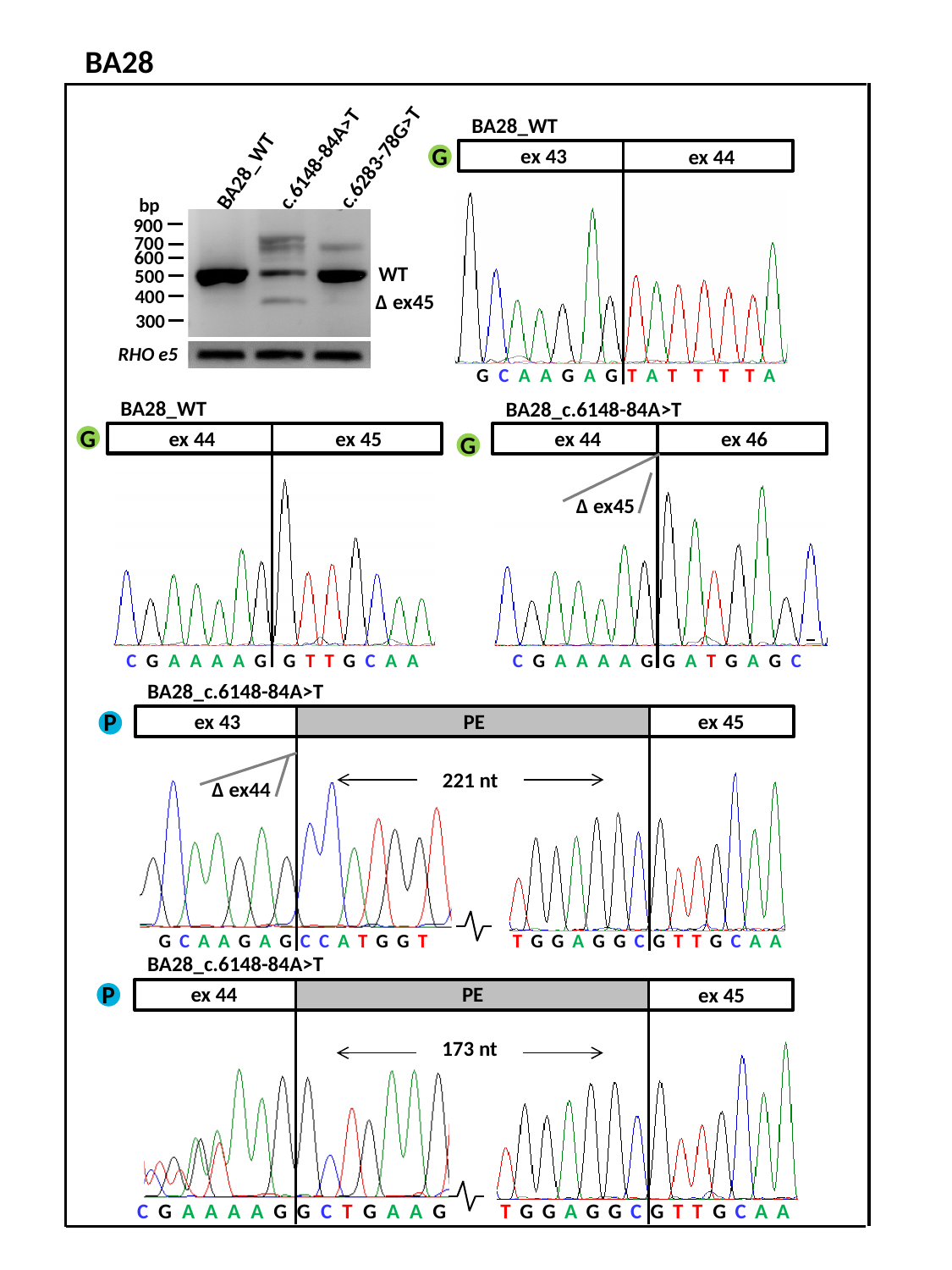

BA28
c.6283-78G>T
c.6148-84A>T
BA28_WT
bp
900
700
600
500
400
300
RHO e5
BA28_WT
 ex 43
 ex 44
G C A A G A G T A T T T T A
G
WT
 ∆ ex45
BA28_WT
 ex 44
 ex 45
BA28_c.6148-84A>T
 ex 44
 ex 46
C G A A A A G G A T G A G C
∆ ex45
G
G
C G A A A A G G T T G C A A
BA28_c.6148-84A>T
P
PE
 ex 43
ex 45
221 nt
∆ ex44
G C A A G A G C C A T G G T
T G G A G G C G T T G C A A
BA28_c.6148-84A>T
 ex 44
 ex 45
173 nt
C G A A A A G G C T G A A G
T G G A G G C G T T G C A A
P
PE

### Slide 7
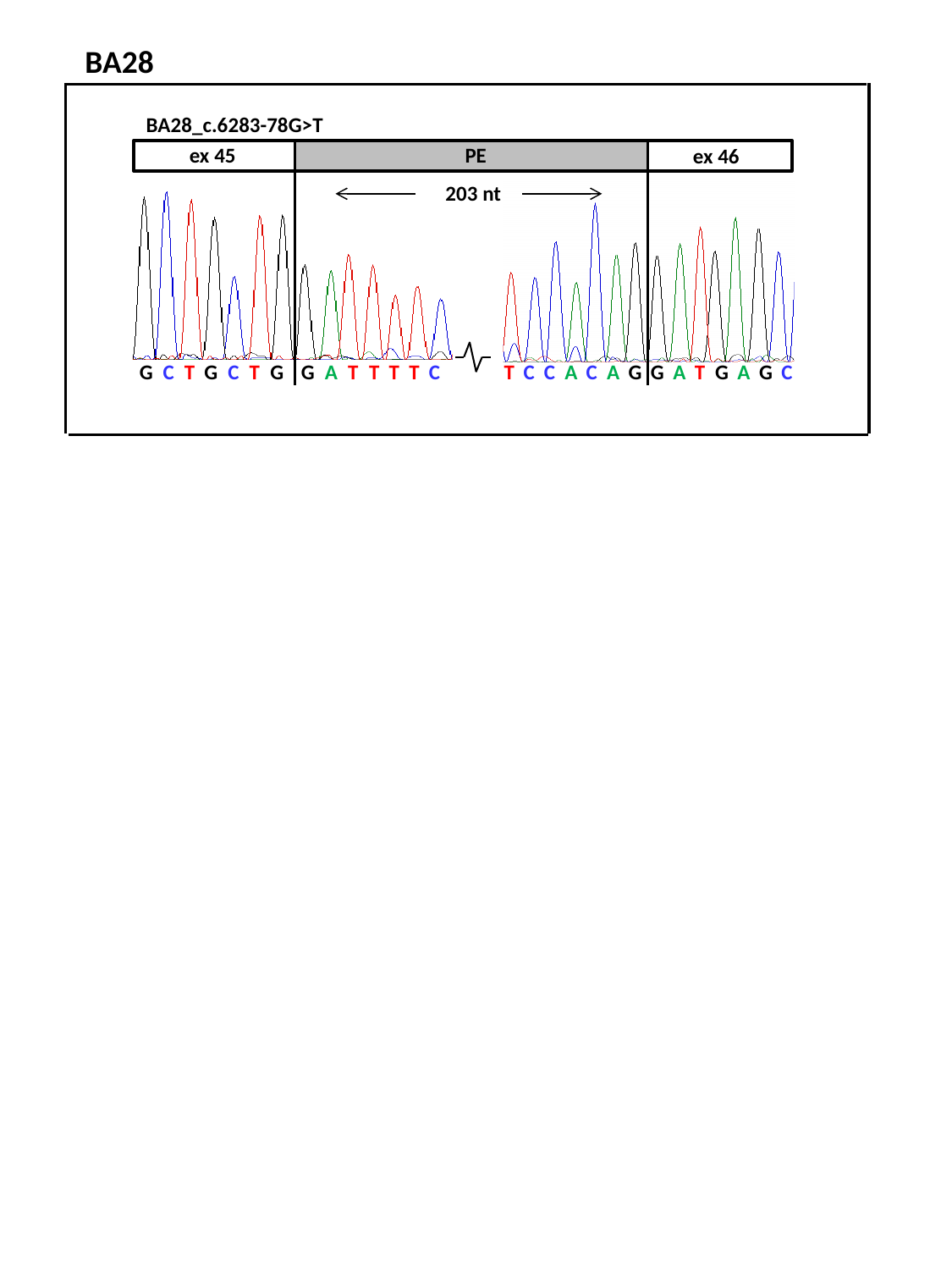

BA28
 ex 45
PE
 ex 46
203 nt
G C T G C T G G A T T T T C
T C C A C A G G A T G A G C
BA28_c.6283-78G>T
