## Supplemental Fig. S7. Control photoreceptor progenitor cells RT-PCR results for "Resolving the dark matter of *ABCA4* for 1,054 Stargardt disease probands through integrated genomics and transcriptomics"

### Slide 1
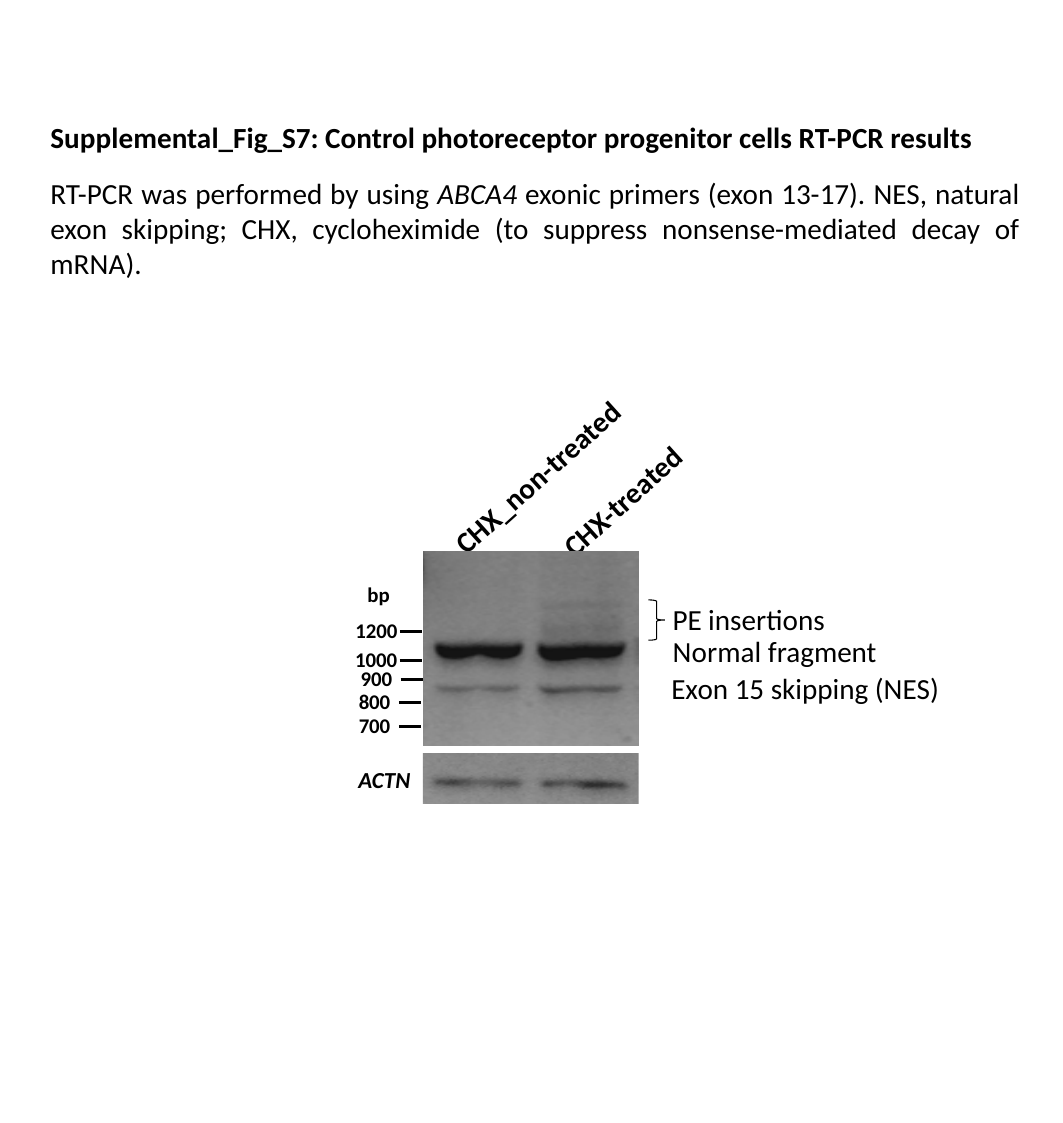

Supplemental_Fig_S7: Control photoreceptor progenitor cells RT-PCR results
RT-PCR was performed by using ABCA4 exonic primers (exon 13-17). NES, natural exon skipping; CHX, cycloheximide (to suppress nonsense-mediated decay of mRNA).
CHX_non-treated
CHX-treated
bp
1200
1000
900
800
700
PE insertions
Normal fragment
Exon 15 skipping (NES)
ACTN
