## Supplemental Fig. S8. Multiple sequence alignment and schematic overview of breakpoint analysis for "Resolving the dark matter of *ABCA4* for 1,054 Stargardt disease probands through integrated genomics and transcriptomics"

### Slide 1
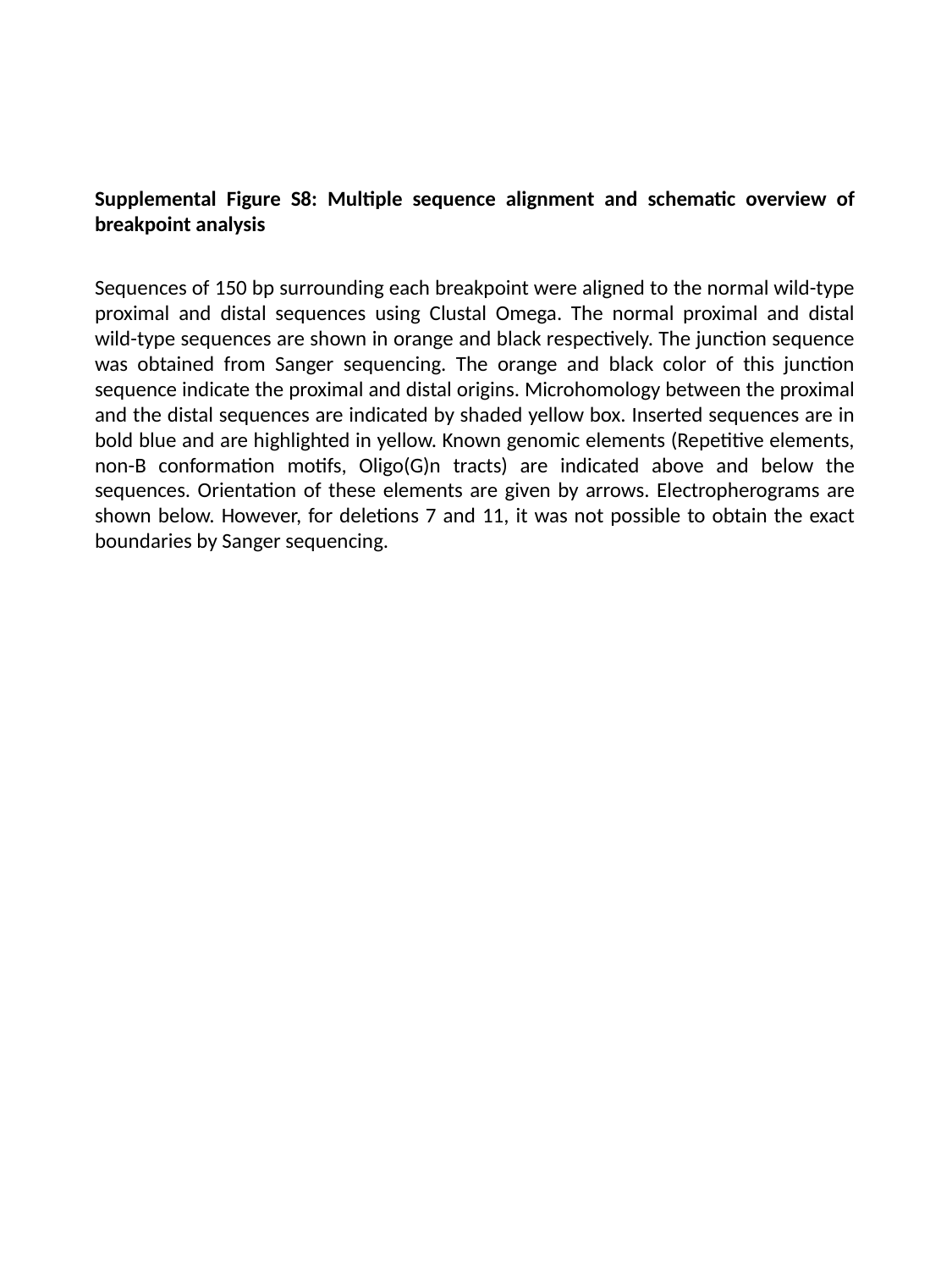

Supplemental Figure S8: Multiple sequence alignment and schematic overview of breakpoint analysis
Sequences of 150 bp surrounding each breakpoint were aligned to the normal wild-type proximal and distal sequences using Clustal Omega. The normal proximal and distal wild-type sequences are shown in orange and black respectively. The junction sequence was obtained from Sanger sequencing. The orange and black color of this junction sequence indicate the proximal and distal origins. Microhomology between the proximal and the distal sequences are indicated by shaded yellow box. Inserted sequences are in bold blue and are highlighted in yellow. Known genomic elements (Repetitive elements, non-B conformation motifs, Oligo(G)n tracts) are indicated above and below the sequences. Orientation of these elements are given by arrows. Electropherograms are shown below. However, for deletions 7 and 11, it was not possible to obtain the exact boundaries by Sanger sequencing.

### Slide 2
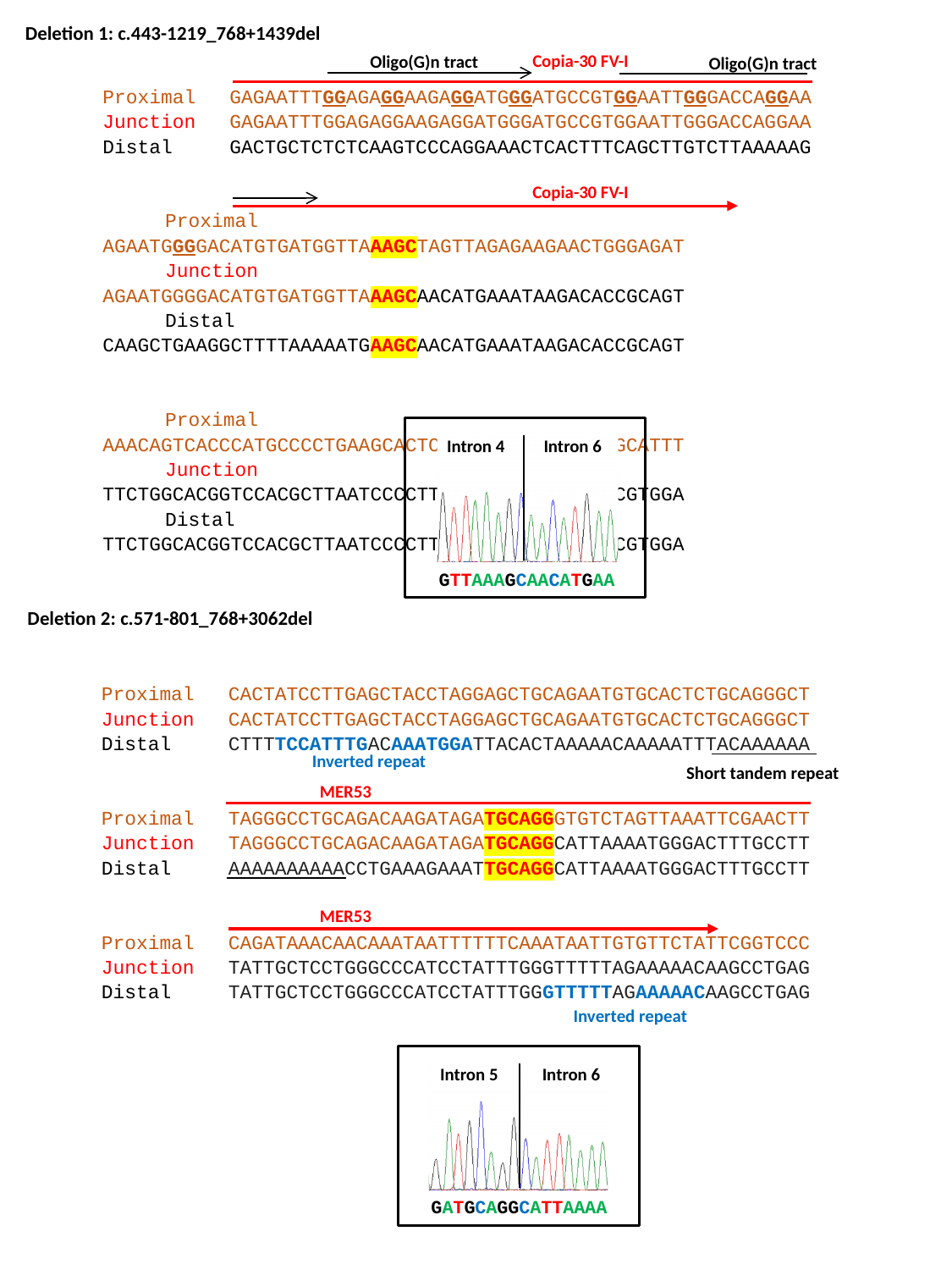

Deletion 1: c.443-1219_768+1439del
Copia-30 FV-I
Oligo(G)n tract
Oligo(G)n tract
Proximal 	GAGAATTTGGAGAGGAAGAGGATGGGATGCCGTGGAATTGGGACCAGGAA
Junction 	GAGAATTTGGAGAGGAAGAGGATGGGATGCCGTGGAATTGGGACCAGGAA
Distal 	GACTGCTCTCTCAAGTCCCAGGAAACTCACTTTCAGCTTGTCTTAAAAAG
Proximal 	AGAATGGGGACATGTGATGGTTAAAGCTAGTTAGAGAAGAACTGGGAGAT
Junction 	AGAATGGGGACATGTGATGGTTAAAGCAACATGAAATAAGACACCGCAGT
Distal 	CAAGCTGAAGGCTTTTAAAAATGAAGCAACATGAAATAAGACACCGCAGT
Proximal 	AAACAGTCACCCATGCCCCTGAAGCACTCGGGGTGAAGAGATTGGCATTT
Junction 	TTCTGGCACGGTCCACGCTTAATCCCCTTCAATGTGTGACTTTCCGTGGA
Distal 	TTCTGGCACGGTCCACGCTTAATCCCCTTCAATGTGTGACTTTCCGTGGA
Copia-30 FV-I
Intron 4
 Intron 6
GTTAAAGCAACATGAA
 Deletion 2: c.571-801_768+3062del
Proximal 	CACTATCCTTGAGCTACCTAGGAGCTGCAGAATGTGCACTCTGCAGGGCT
Junction 	CACTATCCTTGAGCTACCTAGGAGCTGCAGAATGTGCACTCTGCAGGGCT
Distal 	CTTTTCCATTTGACAAATGGATTACACTAAAAACAAAAATTTACAAAAAA
Proximal 	TAGGGCCTGCAGACAAGATAGATGCAGGGTGTCTAGTTAAATTCGAACTT
Junction 	TAGGGCCTGCAGACAAGATAGATGCAGGCATTAAAATGGGACTTTGCCTT
Distal 	AAAAAAAAAACCTGAAAGAAATTGCAGGCATTAAAATGGGACTTTGCCTT
Proximal 	CAGATAAACAACAAATAATTTTTTCAAATAATTGTGTTCTATTCGGTCCC
Junction 	TATTGCTCCTGGGCCCATCCTATTTGGGTTTTTAGAAAAACAAGCCTGAG
Distal 	TATTGCTCCTGGGCCCATCCTATTTGGGTTTTTAGAAAAACAAGCCTGAG
Inverted repeat
Short tandem repeat
MER53
MER53
Inverted repeat
Intron 5
Intron 6
GATGCAGGCATTAAAA

### Slide 3
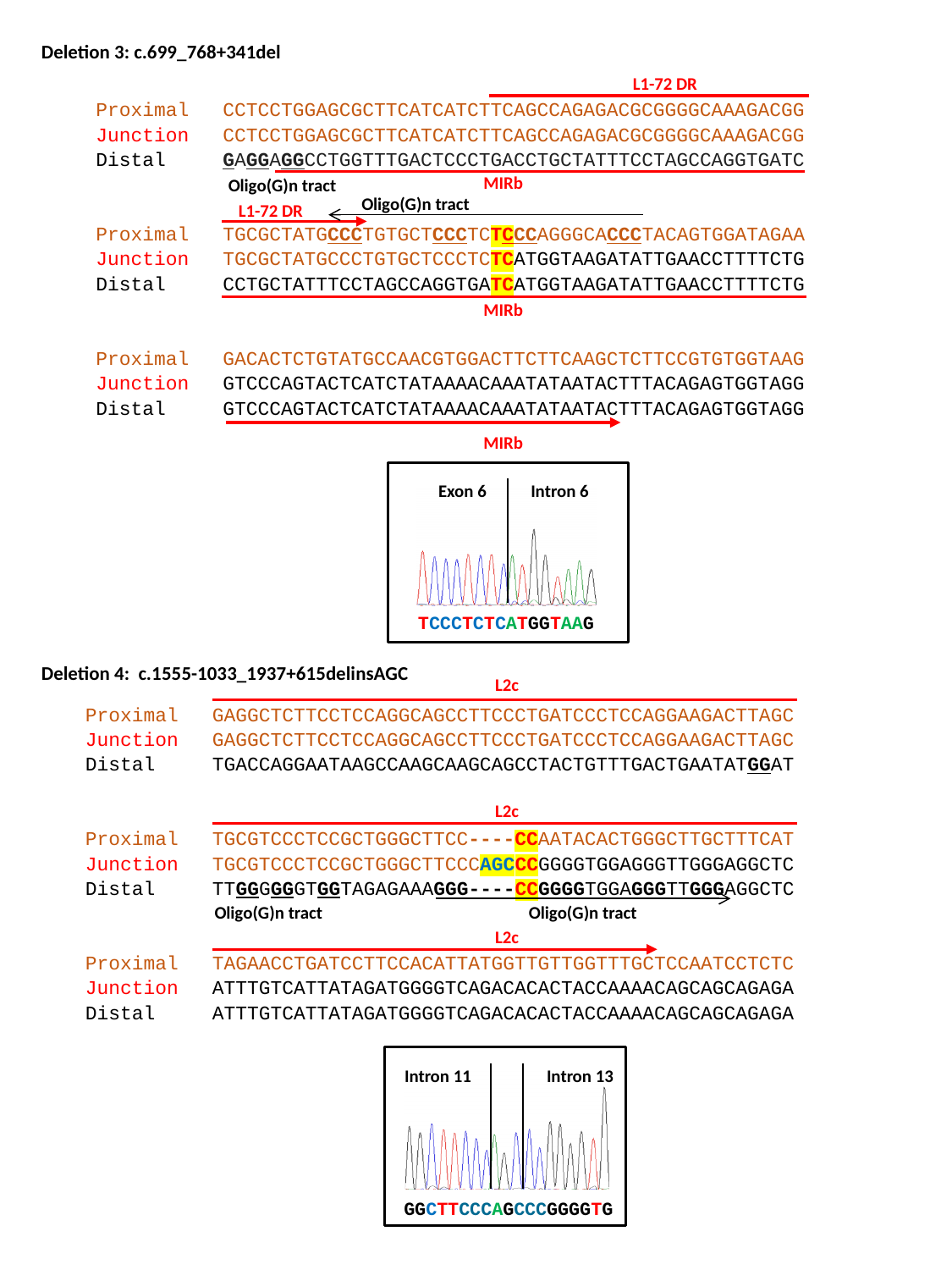

Deletion 3: c.699_768+341del
L1-72 DR
Proximal 	CCTCCTGGAGCGCTTCATCATCTTCAGCCAGAGACGCGGGGCAAAGACGG
Junction 	CCTCCTGGAGCGCTTCATCATCTTCAGCCAGAGACGCGGGGCAAAGACGG
Distal	GAGGAGGCCTGGTTTGACTCCCTGACCTGCTATTTCCTAGCCAGGTGATC
Proximal 	TGCGCTATGCCCTGTGCTCCCTCTCCCAGGGCACCCTACAGTGGATAGAA
Junction 	TGCGCTATGCCCTGTGCTCCCTCTCATGGTAAGATATTGAACCTTTTCTG
Distal 	CCTGCTATTTCCTAGCCAGGTGATCATGGTAAGATATTGAACCTTTTCTG
Proximal 	GACACTCTGTATGCCAACGTGGACTTCTTCAAGCTCTTCCGTGTGGTAAG
Junction 	GTCCCAGTACTCATCTATAAAACAAATATAATACTTTACAGAGTGGTAGG
Distal	GTCCCAGTACTCATCTATAAAACAAATATAATACTTTACAGAGTGGTAGG
MIRb
Oligo(G)n tract
Oligo(G)n tract
L1-72 DR
MIRb
MIRb
Exon 6
Intron 6
TCCCTCTCATGGTAAG
Deletion 4: c.1555-1033_1937+615delinsAGC
L2c
Proximal 	GAGGCTCTTCCTCCAGGCAGCCTTCCCTGATCCCTCCAGGAAGACTTAGC
Junction 	GAGGCTCTTCCTCCAGGCAGCCTTCCCTGATCCCTCCAGGAAGACTTAGC
Distal	TGACCAGGAATAAGCCAAGCAAGCAGCCTACTGTTTGACTGAATATGGAT
Proximal 	TGCGTCCCTCCGCTGGGCTTCC----CCAATACACTGGGCTTGCTTTCAT
Junction 	TGCGTCCCTCCGCTGGGCTTCCCAGCCCGGGGTGGAGGGTTGGGAGGCTC
Distal 	TTGGGGGGTGGTAGAGAAAGGG----CCGGGGTGGAGGGTTGGGAGGCTC
Proximal 	TAGAACCTGATCCTTCCACATTATGGTTGTTGGTTTGCTCCAATCCTCTC
Junction 	ATTTGTCATTATAGATGGGGTCAGACACACTACCAAAACAGCAGCAGAGA
Distal	ATTTGTCATTATAGATGGGGTCAGACACACTACCAAAACAGCAGCAGAGA
L2c
Oligo(G)n tract
Oligo(G)n tract
L2c
Intron 11
 Intron 13
GGCTTCCCAGCCCGGGGTG

### Slide 4
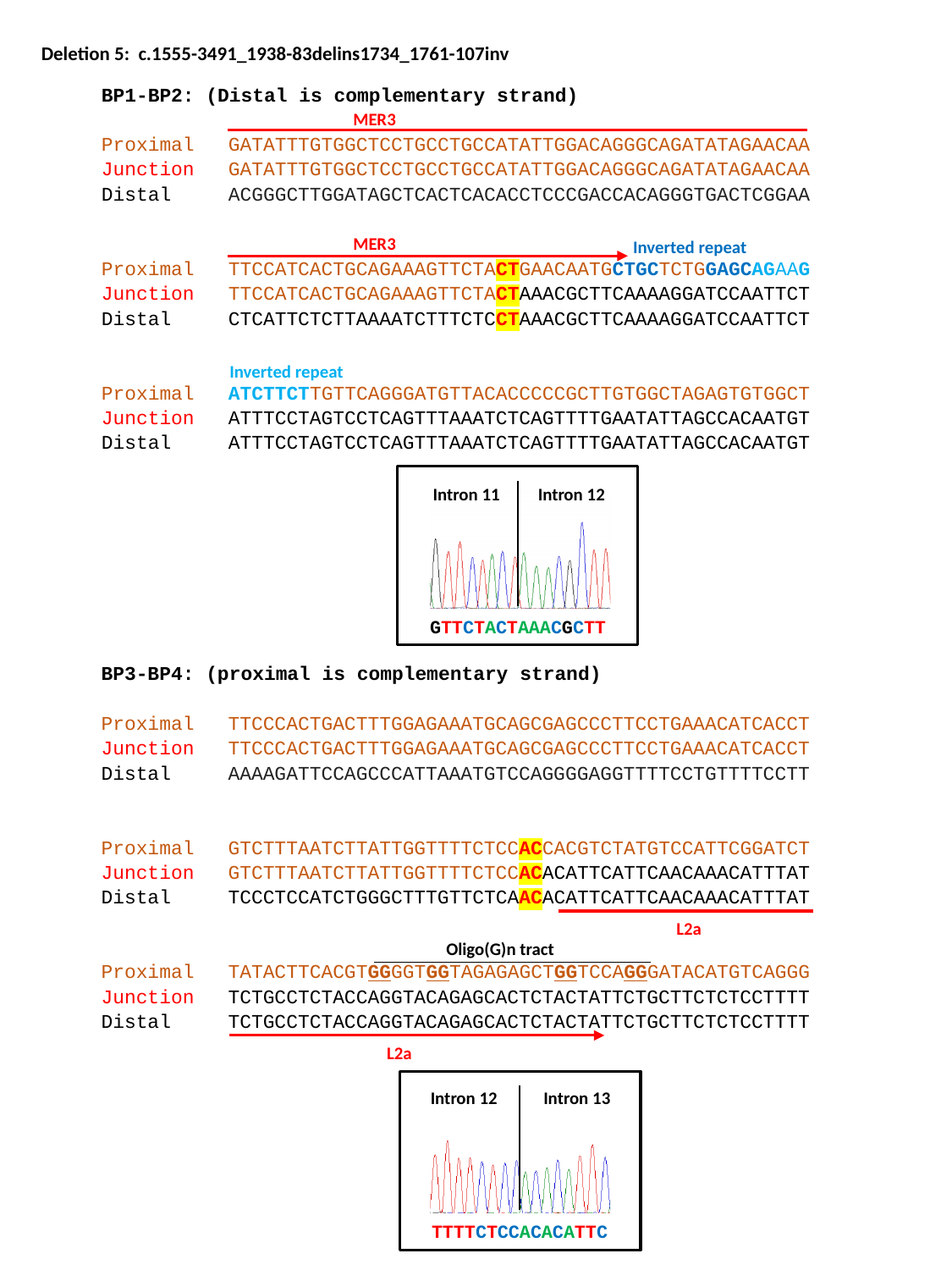

BP1-BP2: (Distal is complementary strand)
Proximal 	GATATTTGTGGCTCCTGCCTGCCATATTGGACAGGGCAGATATAGAACAA
Junction 	GATATTTGTGGCTCCTGCCTGCCATATTGGACAGGGCAGATATAGAACAA
Distal 	ACGGGCTTGGATAGCTCACTCACACCTCCCGACCACAGGGTGACTCGGAA
Proximal 	TTCCATCACTGCAGAAAGTTCTACTGAACAATGCTGCTCTGGAGCAGAAG
Junction 	TTCCATCACTGCAGAAAGTTCTACTAAACGCTTCAAAAGGATCCAATTCT
Distal	CTCATTCTCTTAAAATCTTTCTCCTAAACGCTTCAAAAGGATCCAATTCT
Proximal 	ATCTTCTTGTTCAGGGATGTTACACCCCCGCTTGTGGCTAGAGTGTGGCT
Junction 	ATTTCCTAGTCCTCAGTTTAAATCTCAGTTTTGAATATTAGCCACAATGT
Distal	ATTTCCTAGTCCTCAGTTTAAATCTCAGTTTTGAATATTAGCCACAATGT
MER3
MER3
Inverted repeat
Inverted repeat
Deletion 5: c.1555-3491_1938-83delins1734_1761-107inv
Intron 11
 Intron 12
GTTCTACTAAACGCTT
BP3-BP4: (proximal is complementary strand)
Proximal 	TTCCCACTGACTTTGGAGAAATGCAGCGAGCCCTTCCTGAAACATCACCT
Junction 	TTCCCACTGACTTTGGAGAAATGCAGCGAGCCCTTCCTGAAACATCACCT
Distal	AAAAGATTCCAGCCCATTAAATGTCCAGGGGAGGTTTTCCTGTTTTCCTT
Proximal 	GTCTTTAATCTTATTGGTTTTCTCCACCACGTCTATGTCCATTCGGATCT
Junction 	GTCTTTAATCTTATTGGTTTTCTCCACACATTCATTCAACAAACATTTAT
Distal	TCCCTCCATCTGGGCTTTGTTCTCAACACATTCATTCAACAAACATTTAT
Proximal 	TATACTTCACGTGGGGTGGTAGAGAGCTGGTCCAGGGATACATGTCAGGG
Junction 	TCTGCCTCTACCAGGTACAGAGCACTCTACTATTCTGCTTCTCTCCTTTT
Distal	TCTGCCTCTACCAGGTACAGAGCACTCTACTATTCTGCTTCTCTCCTTTT
L2a
Oligo(G)n tract
L2a
Intron 12
 Intron 13
TTTTCTCCACACATTC

### Slide 5
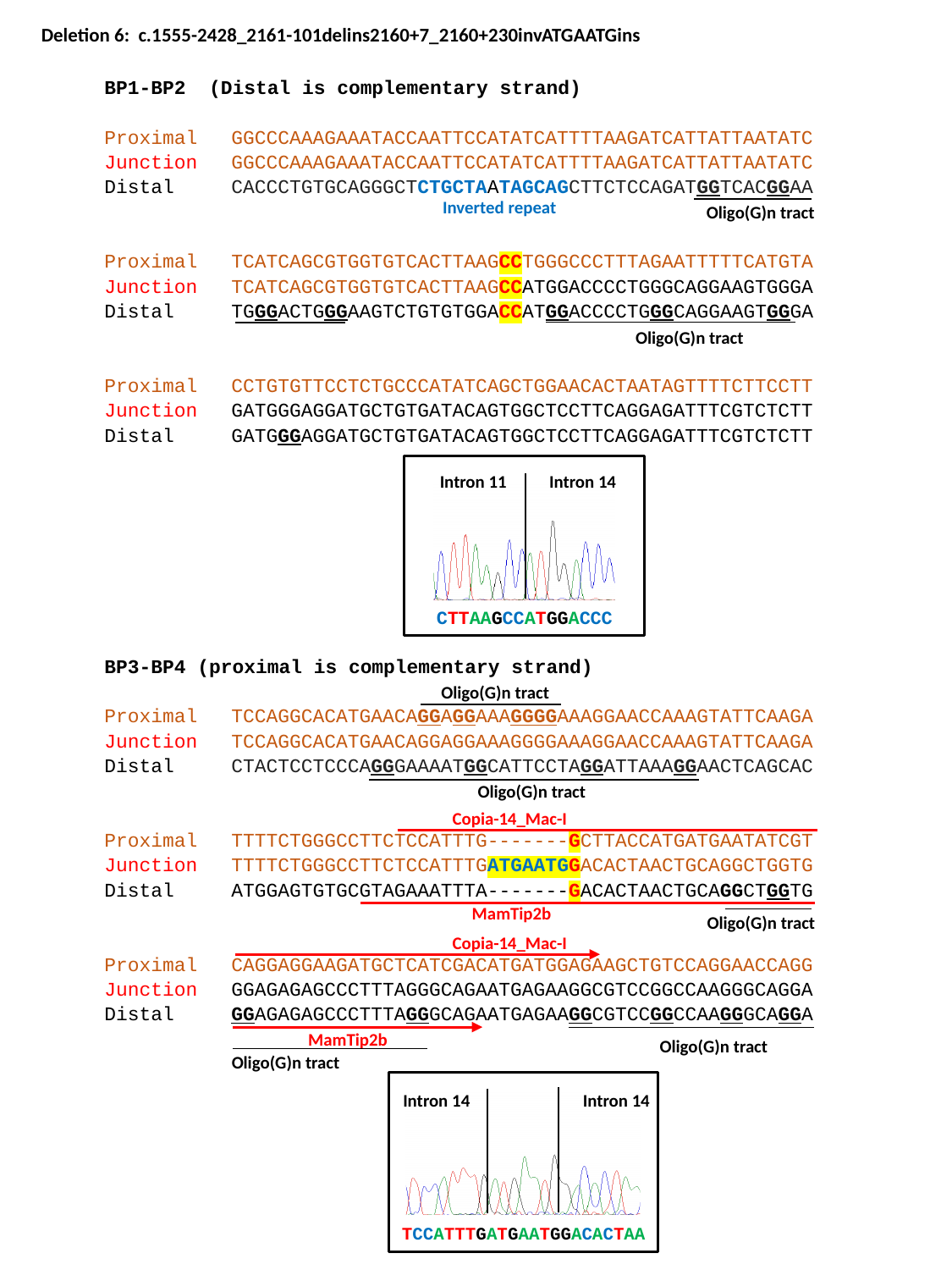

Deletion 6: c.1555-2428_2161-101delins2160+7_2160+230invATGAATGins
BP1-BP2 (Distal is complementary strand)
Proximal 	GGCCCAAAGAAATACCAATTCCATATCATTTTAAGATCATTATTAATATC
Junction 	GGCCCAAAGAAATACCAATTCCATATCATTTTAAGATCATTATTAATATC
Distal 	CACCCTGTGCAGGGCTCTGCTAATAGCAGCTTCTCCAGATGGTCACGGAA
Proximal 	TCATCAGCGTGGTGTCACTTAAGCCTGGGCCCTTTAGAATTTTTCATGTA
Junction 	TCATCAGCGTGGTGTCACTTAAGCCATGGACCCCTGGGCAGGAAGTGGGA
Distal 	TGGGACTGGGAAGTCTGTGTGGACCATGGACCCCTGGGCAGGAAGTGGGA
Proximal 	CCTGTGTTCCTCTGCCCATATCAGCTGGAACACTAATAGTTTTCTTCCTT
Junction 	GATGGGAGGATGCTGTGATACAGTGGCTCCTTCAGGAGATTTCGTCTCTT
Distal	GATGGGAGGATGCTGTGATACAGTGGCTCCTTCAGGAGATTTCGTCTCTT
Inverted repeat
Oligo(G)n tract
Oligo(G)n tract
Intron 11
Intron 14
CTTAAGCCATGGACCC
BP3-BP4 (proximal is complementary strand)
Proximal	TCCAGGCACATGAACAGGAGGAAAGGGGAAAGGAACCAAAGTATTCAAGA
Junction 	TCCAGGCACATGAACAGGAGGAAAGGGGAAAGGAACCAAAGTATTCAAGA
Distal 	CTACTCCTCCCAGGGAAAATGGCATTCCTAGGATTAAAGGAACTCAGCAC
Proximal 	TTTTCTGGGCCTTCTCCATTTG-------GCTTACCATGATGAATATCGT
Junction 	TTTTCTGGGCCTTCTCCATTTGATGAATGGACACTAACTGCAGGCTGGTG
Distal	ATGGAGTGTGCGTAGAAATTTA-------GACACTAACTGCAGGCTGGTG
Proximal 	CAGGAGGAAGATGCTCATCGACATGATGGAGAAGCTGTCCAGGAACCAGG
Junction 	GGAGAGAGCCCTTTAGGGCAGAATGAGAAGGCGTCCGGCCAAGGGCAGGA
Distal	GGAGAGAGCCCTTTAGGGCAGAATGAGAAGGCGTCCGGCCAAGGGCAGGA
Oligo(G)n tract
Oligo(G)n tract
Copia-14_Mac-I
MamTip2b
Oligo(G)n tract
Copia-14_Mac-I
MamTip2b
Oligo(G)n tract
Oligo(G)n tract
Intron 14
Intron 14
TCCATTTGATGAATGGACACTAA

### Slide 6
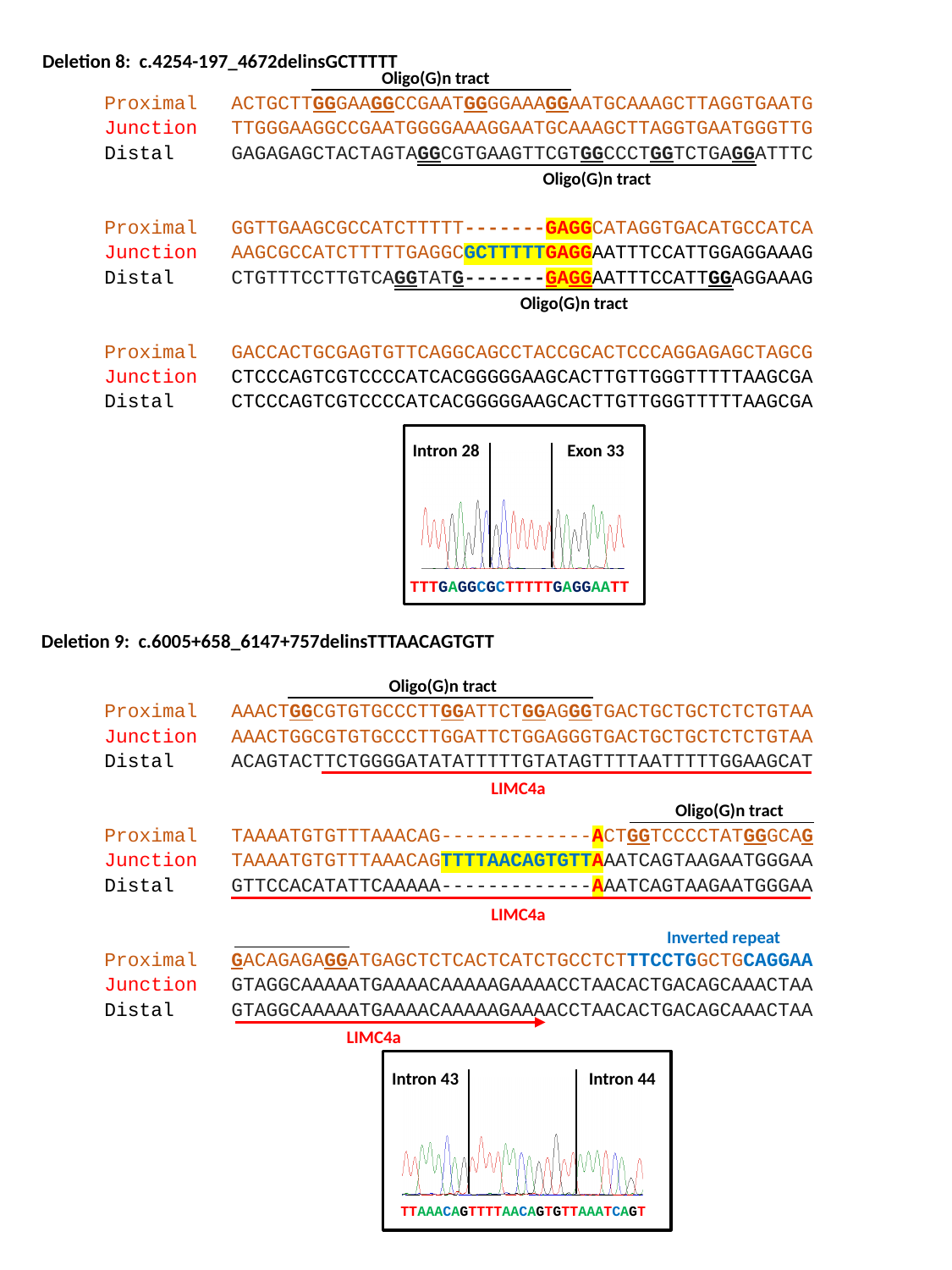

Deletion 8: c.4254-197_4672delinsGCTTTTT
Oligo(G)n tract
Proximal 	ACTGCTTGGGAAGGCCGAATGGGGAAAGGAATGCAAAGCTTAGGTGAATG
Junction 	TTGGGAAGGCCGAATGGGGAAAGGAATGCAAAGCTTAGGTGAATGGGTTG
Distal 	GAGAGAGCTACTAGTAGGCGTGAAGTTCGTGGCCCTGGTCTGAGGATTTC
Proximal 	GGTTGAAGCGCCATCTTTTT-------GAGGCATAGGTGACATGCCATCA
Junction 	AAGCGCCATCTTTTTGAGGCGCTTTTTGAGGAATTTCCATTGGAGGAAAG
Distal 	CTGTTTCCTTGTCAGGTATG-------GAGGAATTTCCATTGGAGGAAAG
Proximal 	GACCACTGCGAGTGTTCAGGCAGCCTACCGCACTCCCAGGAGAGCTAGCG
Junction 	CTCCCAGTCGTCCCCATCACGGGGGAAGCACTTGTTGGGTTTTTAAGCGA
Distal 	CTCCCAGTCGTCCCCATCACGGGGGAAGCACTTGTTGGGTTTTTAAGCGA
Oligo(G)n tract
Oligo(G)n tract
Intron 28
Exon 33
TTTGAGGCGCTTTTTGAGGAATT
Deletion 9: c.6005+658_6147+757delinsTTTAACAGTGTT
Oligo(G)n tract
Proximal 	AAACTGGCGTGTGCCCTTGGATTCTGGAGGGTGACTGCTGCTCTCTGTAA
Junction 	AAACTGGCGTGTGCCCTTGGATTCTGGAGGGTGACTGCTGCTCTCTGTAA
Distal 	ACAGTACTTCTGGGGATATATTTTTGTATAGTTTTAATTTTTGGAAGCAT
Proximal 	TAAAATGTGTTTAAACAG-------------ACTGGTCCCCTATGGGCAG
Junction 	TAAAATGTGTTTAAACAGTTTTAACAGTGTTAAATCAGTAAGAATGGGAA
Distal 	GTTCCACATATTCAAAAA-------------AAATCAGTAAGAATGGGAA
Proximal 	GACAGAGAGGATGAGCTCTCACTCATCTGCCTCTTTCCTGGCTGCAGGAA
Junction 	GTAGGCAAAAATGAAAACAAAAAGAAAACCTAACACTGACAGCAAACTAA
Distal 	GTAGGCAAAAATGAAAACAAAAAGAAAACCTAACACTGACAGCAAACTAA
LIMC4a
Oligo(G)n tract
LIMC4a
Inverted repeat
LIMC4a
Intron 43
Intron 44
TTAAACAGTTTTAACAGTGTTAAATCAGT

### Slide 7
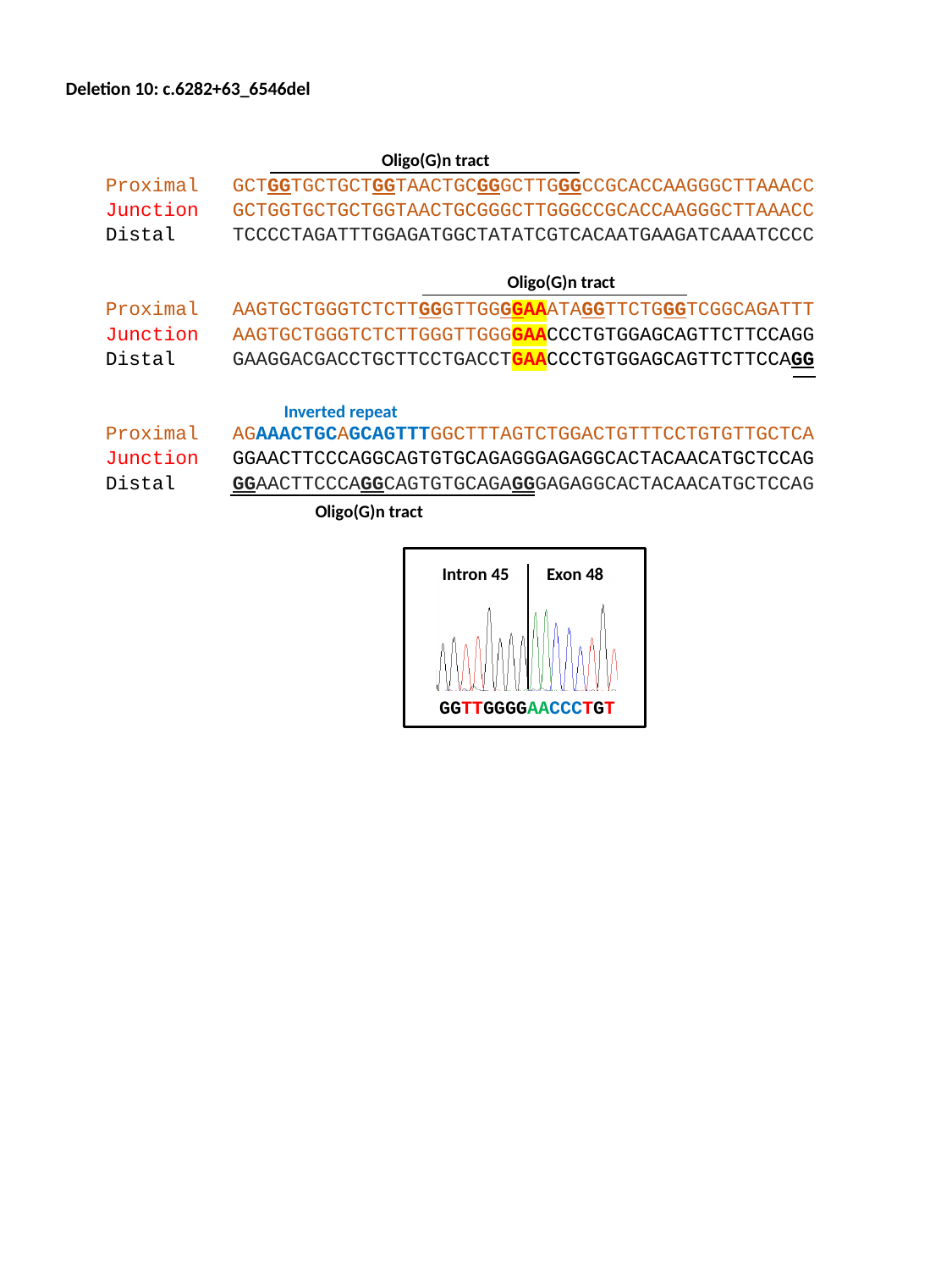

Deletion 10: c.6282+63_6546del
Oligo(G)n tract
Proximal 	GCTGGTGCTGCTGGTAACTGCGGGCTTGGGCCGCACCAAGGGCTTAAACC
Junction 	GCTGGTGCTGCTGGTAACTGCGGGCTTGGGCCGCACCAAGGGCTTAAACC
Distal 	TCCCCTAGATTTGGAGATGGCTATATCGTCACAATGAAGATCAAATCCCC
Proximal 	AAGTGCTGGGTCTCTTGGGTTGGGGAAATAGGTTCTGGGTCGGCAGATTT
Junction 	AAGTGCTGGGTCTCTTGGGTTGGGGAACCCTGTGGAGCAGTTCTTCCAGG
Distal 	GAAGGACGACCTGCTTCCTGACCTGAACCCTGTGGAGCAGTTCTTCCAGG
Proximal 	AGAAACTGCAGCAGTTTGGCTTTAGTCTGGACTGTTTCCTGTGTTGCTCA
Junction 	GGAACTTCCCAGGCAGTGTGCAGAGGGAGAGGCACTACAACATGCTCCAG
Distal 	GGAACTTCCCAGGCAGTGTGCAGAGGGAGAGGCACTACAACATGCTCCAG
Oligo(G)n tract
Inverted repeat
Oligo(G)n tract
Intron 45
Exon 48
GGTTGGGGAACCCTGT
