## Supplemental Fig. S9. Uniparental isodisomy analysis for "Resolving the dark matter of *ABCA4* for 1,054 Stargardt disease probands through integrated genomics and transcriptomics"

### Slide 1
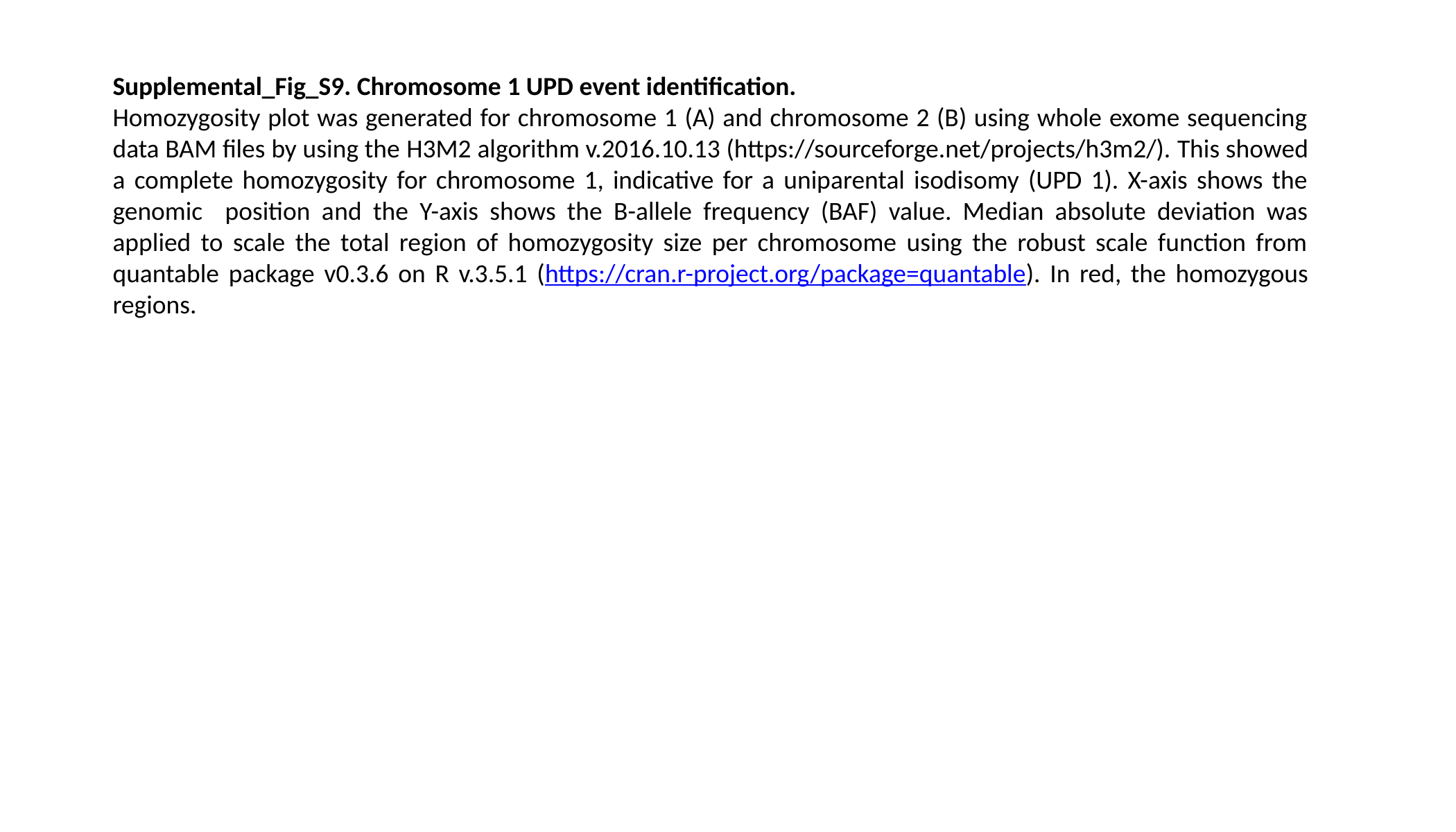

Supplemental_Fig_S9. Chromosome 1 UPD event identification.
Homozygosity plot was generated for chromosome 1 (A) and chromosome 2 (B) using whole exome sequencing data BAM files by using the H3M2 algorithm v.2016.10.13 (https://sourceforge.net/projects/h3m2/). This showed a complete homozygosity for chromosome 1, indicative for a uniparental isodisomy (UPD 1). X-axis shows the genomic position and the Y-axis shows the B-allele frequency (BAF) value. Median absolute deviation was applied to scale the total region of homozygosity size per chromosome using the robust scale function from quantable package v0.3.6 on R v.3.5.1 (https://cran.r-project.org/package=quantable). In red, the homozygous regions.

### Slide 2
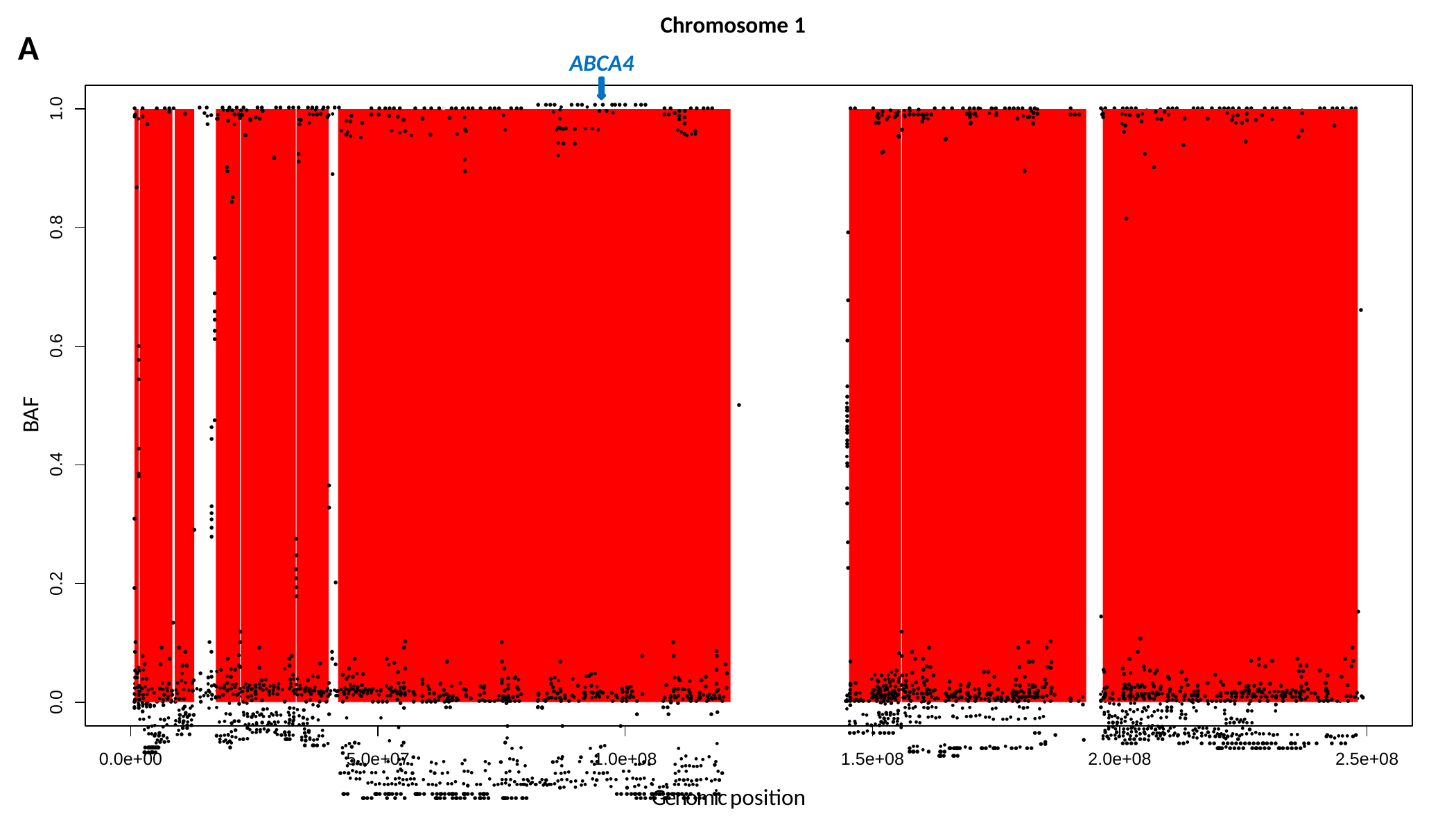

Chromosome 1
A
ABCA4
1.0
● ● ● ●●● ●_●_●
● ● ● ● ● ● ●● ●● ●● ● ●●● ●● ●● ●
●
● ● ●●● ● ● ● ● ●● ●● ● ●● ● ●●● ● ●● ● ●
● ●●● _●_ ● ●● _●_ ● ● ●●● ● ● ●●
_●_	● ● _●_
^●^ ● ●
● ●	● ● ●●● ●●
●●
●
●	● ● ● ● ●●● ●● ●●●●● ●● ●●●●●●
● ●●●
●● ● ●●●● ●
●●● ●●● ● ●●●● ● ●● ● ●● ●● ●● ●●
●● ●● ●●●● ●
●
●
● ●
● ^●^ ^●^	●
_●_	^●^ _●_ ●
● ●
●
●
●
●
●
●
●
●●^●^
● ^●^
_●_● ^●^ ●
● ^●^●●
_●_^●^●
^●^_●_●_●_^●●●●^●●
_●_●● _●_● ●
^●^	^●^	● ●
^●^ _●_	^●^ _●_ ●	_●_	^●^	^●^	^●^ ^●^ ^●^	^●^_●_
●_●_ _●_	_●_	●	^●^	_●_ ^●●^
^●^ ● ●	_●_
^●^	●
●	●	●
●
●
_●_	_●_	●	●	●
_●_	●	●	●	_●_
_●_ ●_●_	^●^ ●	●	_●_
_●_ ^●^ ●	^●^	_●_ _●_	_●_ ●^●^	_●_	●
_●_	●	_●_	_●_^●^	● ^●^	^●^ _●_ ^●^
^●^ ^●^	^●^ ^●●^ ^●^	^●^	^●^	^●^ ● ^●●^	●^●^ ^●^	^●^ ^●^ ^●^	●	^●^	^●^ ^●^
_●_ ^●●●^●	^●^ ^●●^	^●^ ^●^	^●^	^●^	^●^ _●_	_●_	^●^	^●●^ ^●^	^●^ _●●●●●●_	●	_●_
●^●^●^●●●●^ ^●●^ ^●^ ^●●●^ ●	^●^● _●_ ^●^	^●●●^	^●^	^●●^ ^●^	_●_ _●_^●●^	^●^ ^●^ _●_●^●^	^●^ ^●^	_●_^●^ ^●^	^●^● _●_
^●^ ^●●^	^●^ ^●●●●●●^ ^●^	^●^ ^●●^ ^●^_●_ ^●●^ ^●^● _●_^●^ ^●^_●_ _●_ _●_ ^●^● ^●●^ ^●^ ^●^ ^●●●●^ ^●^	_●_^●^●^●^_●_ ●^●^ ^●^ _●_^●^● _●●_^●●^_●_^●^ ●_●_
^●●●●^ ^●●●●●●^ ^●●●^ ^●●●●^ ^●●●^ ^●^● ^●●^ ^●●●●●^ ^●^ ^●●●^ ^●^ ^●●●●^	^●●^ ^●^	^●^ ^●^ ^●●^ ^●●●●^	●^●^ ^●^	^●^ ● ^●●^ ^●^ ^●^ ^●●●^
●●^●●^ ^●^ _●_	●^●^	● ^●●^ _●_●^●^●●	● ●_●_^●●^ _●_	_●_ _●_●^●^ _●_●	^●^	^●●^ ^●^	●	● ●^●^ ●●
●●●	●●●● ●●●●●●●	●●●●●● ●●●●●●●●●● ●●●●● ●●●●●	●● ●●●●●●● ●●●●●●● ●●●●●●●●● ● ●●● ●●●● ● ● ●	●●●●● ●●●●● ● ●●●● ●●●●●●● ●●●	●●● ●● ●●● ●●● ●●●
●
●
●
●
● ● ^●^
●
^●^● ^●●●●^
●	_●_
^●●^ ^●^ ^●^●^●^ _●_
_●_^●●^ ^●●^ _●●_●
^●^ ●●●
_●●_ ^●^ _●_●●^●^
● ● ●●●●●●●●●●●●●● ●●●●●●●
^●^ ●●
^●^	●
●
●
●
●
●
_●_● ^●^● ^●^
^●^●
●●
^●^ ^●^ ●
_●●_ ^●^_●_● _●_ ●
_●_●
^●^ ●^●●●^_●●●_
^●^ ^●^● ^●●●●^
●
● ● ^●^
●
_●_ ●
●
●
●
●
●
●
●
●
_●_	●
●^●^	^●^
^●●^ ●^●^● ^●^ ^●^ ^●^ ^●^
^●●^●^●●●^ ^●●●●^	^●^● ●^●^
^●^ ● ● ^●●^ ^●●●^
^●●●●^ ^●●●^ ^●●^ ●
^●^ ^●●●^● ^●^
^●●●^ ^●^ ● ●● ^●^●^●●●^
●^●^	●●●●
^●●^ ^●^ ●^●^
_●_	●
●
●
^●^	●
^●^ ●
^●^ ●
●
^●^●_●_ ● _●_ _●_●^●^
^●●^ ^●^ ^●●●^● ●^●^
^●^● ^●^ ●^●^
_●_●_●_ ^●●^ ^●^●^●●●^
● ●●● ●^●^
● _●_
●_●_ _●_ ^●^
●
●
●
●
●
●
●
^●^ ●● ^●^ ^●^
^●^ _●_ ●^●^_●●_
^●^ _●_ ● ^●^
●^●^ ^●●^ ● ^●^ ●^●^ ^●^ ●●
●
●
● ●●
^●●^ ^●●●●●●^ ^●^●^●^ ^●●^
^●^ ^●^	●^●^ ●^●^
^●^ ^●^ ^●^●^●^●^●●^ ^●●^ ^●^
●●^●^ ●^●^ ● ●●●●
_●_ _●_ ●	^●^ ^●^ ^●^	_●_ ^●^
_●_	^●^_●_	^●^	●
_●_	^●^	^●^ ●
●
●
●
●
●	●
●	●	●
●
● ●	_●●_
●
●● _●_	^●^
_●_●	_●_	^●●^
●_●_	^●^	_●_ ^●^	_●_
^●●^ ^●^ ^●^●^●●^	^●^ _●_ ^●^	_●●_	^●^●^●^ _●_ ^●^
^●●^ ● _●_^●^_●●_ ●_●_ ^●^● ^●●^ _●●●_ _●●●_ _●_ _●_ _●_ _●_^●^ ^●^ ^●●^
^●●●^ ^●●●^ ^●●●●●●●^● ^●●●●●●●●^ ^●●●^	^●^	^●^●^●^ ^●●●^●^●●●^ ● ^●●^
●●
●
^●●^ ^●^ ^●●^ ^●●^	^●●^ ^●●●●^ ^●^● ●^●^ ^●^●● ● ^●●^ ^●^ ●●^●^ ^●^ ^●●●●^ ^●^
^●●^	^●●^ ^●●^ ● ^●^ ●
●
●
●●
●
●^●●^	^●●^ ●●●●●●●● ^●^ ● ●●●^●^ ●●●● ^●^ ●●^●^ ●● ●●●● ^●^ ●	●●●	●●●● ●●●
●	^●^_●●_ ^●^_●_● ^●^ ^●●^	^●^ _●_ _●_ ^●^ _●_	●
^●^	^●^	^●●^	_●_	_●_	_●_ _●_ _●_ ●
●	^●^ ●
●	●
●	^●^
●
●
●
●
●
●
●
●
●
●
●	_●_ ●● _●_	●	● _●_
● ●	●
^●^● ^●^	^●●^ ^●^ ^●^	●	^●^	●● ●
● ^●^	^●^ _●_	●	_●_	_●_ _●_
^●^ ●
^●^ ^●^ ●	^●^	^●^ ^●^ ^●^	^●^ ^●^	^●^	^●^
_●●_ ^●^_●_^●^_●_^●^ _●_^●^ _●_^●●^ ●●	^●^ ^●^	^●^	●^●^ ^●^ _●_ _●_ ^●^	●^●^	^●^
^●^ ^●^ ^●^_●_^●^ _●_ _●_ ^●●^ ^●^ ^●^ ^●^ ●	_●_ ● _●●●_ _●_●●_●_^●^ _●_^●^_●_	_●●_● ^●●^_●_ _●_ _●_●^●^ ●^●^ ^●●^ ^●^●
_●_^●^ ^●^ ^●●●●●^ ^●●●^●^●●^ ^●●^ ^●^ ^●●●●^ ^●●^	^●^ ● ● ● ●_●_^●^ ^●^_●_● _●●●●_^●^ _●_ _●_ ^●●^ ● ^●^_●●_ ^●^
^●●^ ● ^●●^ ^●^ ^●^	^●^● ^●●^ ^●^ ^●^ ^●●●●^ ^●^●^●^	^●^ ^●^	●	●
^●^●^●●^●^●^ ^●^ ^●●^ ^●●●●●●^ ^●●●^ ^●^_●●_ ^●●^● ●●_●_●^●^ ^●^	●●_●_^●●^ ^●^ ● ^●^ ^●^ ^●^ _●_ ●● ●●_●_ ^●^●	^●^ ●
●●●●	●●^●^	● ●● ●●●●●●●●● ●●●●● ●●● ●● ● ● ●●	●●●●●^●^●●●● ●●●●●● ●●●●●●●^●^
_●_● _●_	● ●
^●●^● ^●^ ^●^
^●●^ ^●^ ●^●^ ^●^
● ● ^●^
●
● _●_	_●_	^●^
●
●
●
●_●_
●
●
●●_●_
● ^●^
●
●
●
●	●
●
●
●
●
●
●
^●^●
●
●
●
●
●●
●
● ● ^●^
●
●
●
●
●
●
●
●
^●^●
●
●
●
●
●	●
^●^ ●
●
●
●
●
●
●
●
●
●
●
●
●
●
●
●
●
●
●
●
●
●
●
●
●
●
●
●
●
●
0.8
●
●
●
●
●
●
●
0.6
●
●
●
●
●
BAF
●
●_●_
^●^●
●
●
●
●
●
●^●^
●
●
●
●
●
●●
●
●
^●^●
0.4
●
●
●
●
●
●
●
●
●
●
●
●
●
●
●
●
●
●
0.2
●
●
●
●
●
●
●
●
●
●
●
●
●
●
●
●
●
●
●
●
●
●
●
●
●
●
●
●
●
●
●
●
●
●
●
●
●
●
●
●
●
●
●
●
_●_ ●
●
● ●
●
●●
●
● ●
●●
●
●
_●_ ● ●
●
●
●
● ^●^
●
●
●●
●	●
●
●
●
_●_ ●
●
^●^ ●
●
●
●●^●^
●
●
●
_●_ _●_● ^●^● ●
●
●●^●^ ●
● ^●^●
●
_●_ ●
●
● ^●^
^●^ ● ●
●
●
●
^●^ ●
_●_ ●^●^
●●
●
●
●
●
●
●
●
^●^ ● ^●^ ^●^ ●^●^
●
^●^●
●
●
●
^●^● _●_
●
^●^ ^●^● ^●●●^
●^●^_●_ ^●^
●
●
●
●
●
●
●^●^	^●●^ ●
_●_●
^●^ ● ^●●^●
●
●
●
●
●
●	●
●
●
●
●
_●_ _●_ ●
●_●_
● _●_
●
^●^ ● ^●●●^ ^●^
●
●_●_●_●_ ^●^_●_
● _●_ ● _●_ ^●^
●	●
●
●
^●^_●●_●
●
^●^ ●	●
^●^ ● ^●^ ^●●^ ^●●^ ●^●^● ^●^ ^●^ ^●^ ^●●^
●
● ^●^ ^●^
●	●
●
^●^●●
● _●_ ● _●_
●
● ●
●
● ^●^ ^●^ ●	^●^●^●^
●^●^
●
●
●
●
●^●^ ●^●^
●●
_●●_ ^●^●
●
●^●^
^●^ ^●^●	^●^
^●●^ ●_●_ ^●^ _●_^●^●_●_^●^● _●_^●●●^	^●^_●_ _●_ ●_●_ ^●^ _●_●^●^
●
●
●
●
●
●
●●
_●_● ^●^ ^●^
●
●
● ●
● _●_ ^●^_●_●_●_^●^●●
●
●
^●^●●
●^●^
^●●^●^●^
●●
● ●
●
●
●
^●^ ●
^●●●^ ^●●^ ^●●^ ●^●^ ^●^ ^●^●^●●^ ● ^●●^ ●^●^ ●●^●^
●
●	● _●_● _●_●● ●
● ●
●
● ● ●
●
^●^●
●
●^●^ ^●^●	●
●
●
^●^●
● ●	●
●
^●^● ^●^ ●^●^ ^●●^ ^●●●●^●^●^ ^●^ ● ^●^ ●^●●^ ^●●●^●^●●●^ ^●^ ^●^ ^●^ ●^●^ ●
●
● ●
●
●
●	●
●^●^ ^●^● ●^●^
●
●●	●
●
●	●
●^●^
●
● ●
●
●	●
●
●	●
^●^ ^●^ ●^●^
_●_●● ●
●
●●●
●	●
●^●^ ^●^	^●^
●
● ●
^●●^ ● ● ^●^ ● ●●
●●	●
●^●●●^ ●
●
●
●
● ●^●^ ^●^	^●^
● ^●^ ●● ● ●	●
● ●^●^
^●^ ●
●
●
●
●
^●●^ _●_● _●_^●^● ^●^ _●_ ● ^●^ ^●^ ● _●_ ●^●^ _●_ _●_^●●●^ _●_	●^●●^
●
●	^●●^ ●
●●
●
● ●	●
●
^●^ ●^●●●^ ^●^	^●●^
● ●	●	●
^●^ ●
0.0
^●^●
●●●
●^●^ ●
^●^●●^●^ ^●^ ●_●_● ^●^ ^●●●●^_●_
● ●
●
●●
_●_●_●_ ^●^ ^●●^ _●●_●_●●_ ●^●^ _●●_ _●_ _●_
● ●
● ●
^●^ ●
●
● ●
●
●	●
● ●
●
^●^●^●^ ●^●^ ●^●^● ^●^ ●
● ● ● ●	^●^
●	●
● ●
● ●
● ●^●^ ^●^●
●^●^ ●^●^ ^●^ ●
● ^●^ ^●^
●
^●^● ^●^ ●
●
●●●^●^ ^●^
^●●●●●^ _●_ _●_●^●●●^_●_ ●
●
● ^●^ _●_●
●^●^ ●● ●	●
●
●
●
^●^●	● ^●^
●
●
●
●
● ●
^●^● ● ^●^ ^●●^ ^●^ ^●^	● ^●^
●● ●	●	●
●
^●^ ^●^ ● ●
●
●
●
^●^	● ● ^●●^ ^●^ ●
^●^ ^●●^ ^●^●^●●●●^ ● ^●^ ●^●●●●^ ^●●^ ●^●●^	●
● ^●^	●	^●^
● ● ● ●
●●
●	●	● ^●^
^●^ ● ^●^	^●●^ ●	●
●
● ^●^ ^●^ ^●^ ^●^
^●^●
^●^ ●
^●^ ●^●^	●
● ● ^●^●●^●^
●
^●^●
●
^●^ ^●●^ ^●^ ^●^● ●^●●^ ^●●●^ ^●^ ^●^ ^●^ ^●●●●^ ^●●^
●
_●_ _●_●^●^ _●_● ^●^ ^●^ ^●●●^ ^●^ ^●^ ^●●●^ ^●^●
●●
●	^●●●^ ●^●^ ^●^ ^●^ ●
^●^ ^●^● ●	^●^
● ●● ●
●
●
^●●●^ ● ^●^ ^●^ ^●●^ ^●^● ^●^●^●^ ^●^●
^●^	●	^●^
●	●
^●^ ^●^ ^●^ ^●●●●●^●●
●
● ^●^●	●^●^
● ● ●●	●
● ●
^●^ ^●^ ^●^ ^●^ ●
●
●
●
● ●
●
●
^●^	^●●^ ● ^●^ ^●●^ ^●^ ^●●●^ ^●●●^ ^●^ ^●^ ^●^ ^●●^
^●^ ^●^ ^●^ ●^●^
^●^ ●
●
● ^●^	^●^ ^●^●^●^ ●
● ^●●^ ●●
●
●^●^● ^●^ ^●●●●^	^●^
^●●^● ● ^●^ ^●^
● ●
●
●^●^ ● ● ● ●●	● ●●
●	●^●^ ● ● ●	● ●●●	● ●●●● ●● ●
●	● ●●●● ● ●●●● ●
● ● ● ● ●	● ●● ● ●●
● ●	● ● ● ● ●●
●●●●● ● ● ●●	●	● ● ● ● ● ●● ●● ●	●
● ●
●● ●
●	● ● ●●●●● ●● ● ● ● ● ●	●●● ● ● ●●
0.0e+00
5.0e+07
1.0e+08
1.5e+08
2.0e+08
2.5e+08
Genomic position

### Slide 3
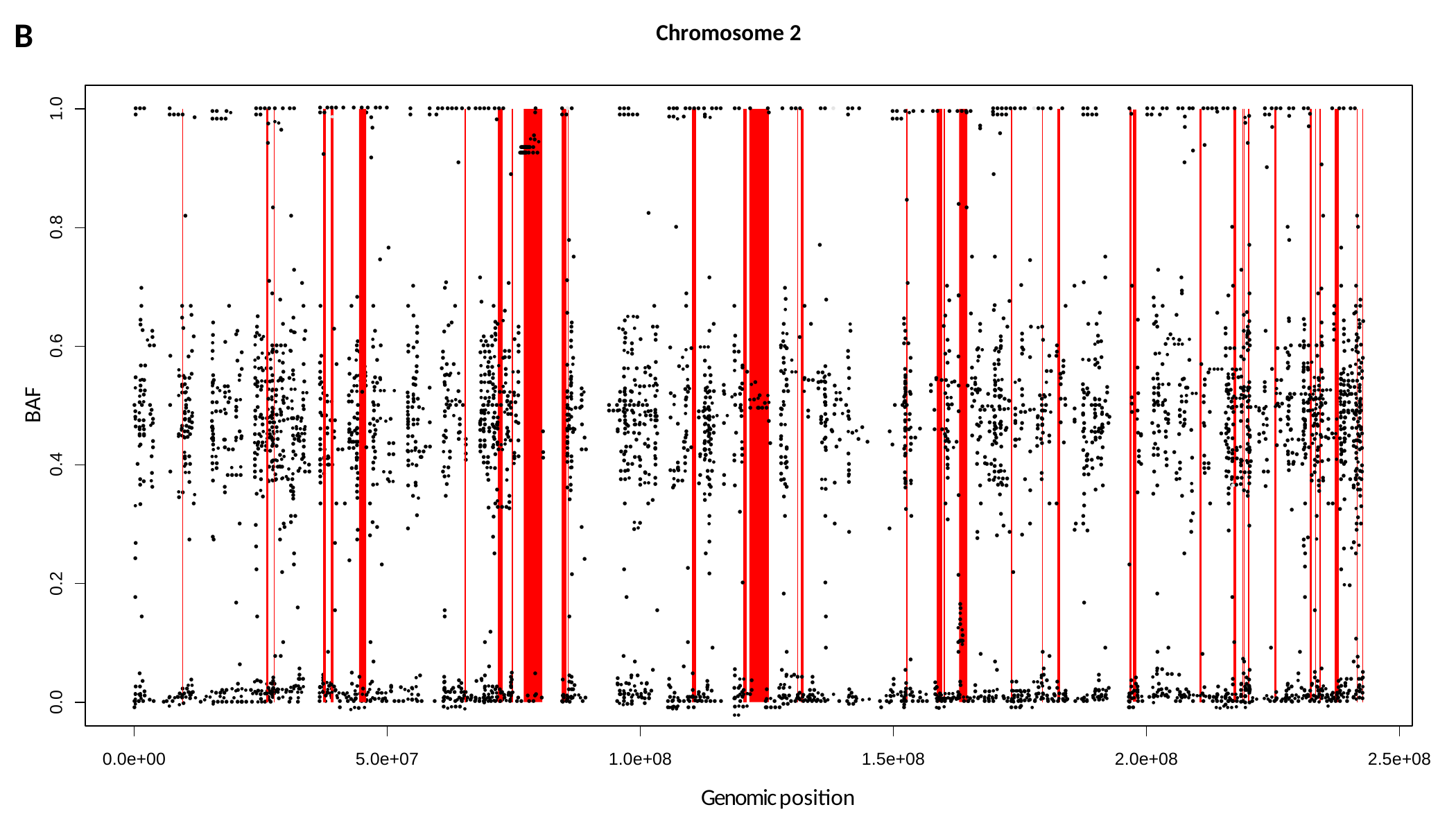

B
Chromosome 2
1.0
●●● ●
● ●●●●
●● ●_●_ ●●●●
●●●● ● ● ●● ●●
● ^●●^
● ●●● ● ● ● ● ●● ● ●●
^●^	●
●●
● ●● ●●● ● ● ●●●● ●●● ● ●
●
● ●● ●●
●●● ●●●●●
●●● ●●●● ● ●●● ●● ● ●●_●_ ●
●
● ●●●
●● ●
●● ● ●
●● ●_●_● ● ●● ● ●●●● ●●●
●●●●●● ●● ●●● ● ● ●●●●
●● ● ●●●●
●
●
● ● ●● ●● ●● ●●●● ●● ● ●● ●
● ●●● ●● ●● ●●
● ●● ●●
●
●
● _●_
●
●
●
^●^●
^●^ ●
●●●●●
●
● ^●^●_●_
●●●●●●●●●●●●● ●●●●●●●●●●●●
●
_●_●
● ^●^
^●^●^●^
●^●^
● ●●●●
●
●
●
●
●
●
●
●
^●^●
●^●^_●_
●●●●●^●^
_●_●●
●
●
●
●
●
●
●
●
●
●
●
●
●
●
●
●
●
●
●
●
●
●
●
●
●
●
●
●
●
●
●
●
●
●
●
0.8
●
●
●
●
●
●
●
●
●
●
●
●
●
●
●
●
●
●
●
●
●
●
●
●
●
●
●
●
●
●
●
●
●
●
●
●
●
●
●
●
●
●
●
●
●
●
●
●
●
●
●
● ●
●
●_●_
●
●
●
●
●
●
●
● ●
●
●
●
●
●
●
●
●
●
●
●
●
●
●
●
●
●_●_●_●_
●●●
●
●
●
●
● ●
●
●
●
●
^●^ ^●^●
^●^● ^●^
●
●
●
●
●
●
●
●
●
●
●
●
●
^●^●●
● ^●^
●●
●
●
●
●_●_
●●
●
●●
●
●
●
^●^●
●
●
●●
●
●
●
●
●
●●
●●
●
●
●
●
^●●^●
●
●
●
●
● _●_●
●
●
●
●
●●
●●
●
●
●●
●●
●●
●
●
●●
●
●●●
●●
●_●_ ●
●
●
●
●
●●
●
●
●
_●_●
●
●●
● ●^●^
●●
●
●
●^●●^ ^●^
●
●●●
●^●^
●
●
●●
0.6
●
●
●
^●^●
_●_ ●
●
●
^●^ ●
●
_●_ ●
●
●
●
_●_ ●
●●
●●
●
●
●●
●
●
●
●●
●
^●^ ^●^ ● ^●^
● ^●●●^
●^●^ _●_
●
●
_●_ ●
●
●●
_●_●
●●
●●●●	●●
●
●●
●●● ●●
●●
_●_ ● ^●●●^
● ●●
●
●_●_ ●● _●_
●●
●●● _●_ ●
●
● ●●● ● ●
●
●●
●
●
● ●
● ^●●^
^●^●
●
●
●
●
●
●
●
●
●^●^
^●^ ●^●^ ^●^ ^●^
●
●● ●●
^●^ ^●^●
●
●●
●
●●
●
●
●
●
●
^●^ ●_●_●
●^●^
●
● ●_●_ _●_
●
● _●_
● ●
●_●_
●
_●_●
^●^ ●^●^ ●
● _●_
●
●
●
●●
●
^●^ ●
^●^ ^●●^●
●●
●
●
●
_●_ ●
●
●
●_●_
●
●^●●^ ●●
●
●
●
●
●
● ^●^
●
●
●●
●
●
●
●
●
^●^ ●● ^●^ ^●^
● ●
●●
●
^●^ ●
^●^●
_●_ ●●
●
^●^ ●●
●
●^●^ ● ^●^
●
●●
● _●_
^●^ ^●^ ^●●^ ^●^●^●^
●
●
●
● ●
●
●
●_●_
●
●
●
●^●^
●
●^●^
●_●_
●
●
●
●
^●^●●
●
● ^●●^
^●^ _●_●
●● _●_
●
●
●^●^ ^●^
●
●
●
_●_● ●
^●^ ●
●
●
●
^●^●
●●
●
●
●●
_●_●
●
● _●_
^●^ _●_ ●
●^●^
●
●●^●●^
● _●_
●
●
●
●
●● ^●^
^●^●●^●^
●
●
●●
●
●●
^●^ ^●^● ^●^
●●●
●●
● ^●●^ ^●^ _●_●
● ●
●●
●^●^
●
●●
●●
^●^●● _●_
●
●_●_
^●^ ●● _●_
●
●●
^●●●^●^●^●●
_●_●
●●
●●
●●
●
●●
●●
_●_●
●
●●	●
●
●●
●●
●●
●●
●
^●^●
●
●● ^●^
_●_ ●^●^
^●^ ●●
●●●● ●●
^●^●●
●
●●
●
●
●
●●
●
●
●
●
● ●
^●●^ ● _●_
●●
●●
BAF
●
●●
_●_●
● ●_●_
●
● _●_
^●^ _●_ ●
●
^●^●● ^●●●^ ^●^
●
●
●●
●●
● ^●^
^●^ ^●^●● ^●●^ ●●
● ^●●^ ^●^ ^●●^
●
●
●^●●^
●
●●
●●
●
●●
●
●●
●● _●●_●●
●●
●
●
● ●●
_●_●
●
●●
●●
●
^●^● ^●^ ^●^
●
●
●●	●
●
^●^ ●^●^●^●●^ ●● ^●●^
●
●●
●
●●
^●^ ●	^●^
●
^●^●● ●●
●
●●
●●
●●
^●^ ^●^ ●
●	^●^ ^●●^ ^●^
^●^ ●
^●^ ^●^ ^●^ ^●^ ^●^●
^●^ ●●
●
^●^ ^●●●^●●
●●
● ^●^ ●^●^
● _●_
_●_ ●
^●●●^ ^●^●●
●
^●●^●●
^●●^ ●●
●●^●^ _●_
●
^●^● ^●^
●
●
_●_ ●●
●● _●_
●
_●_ _●_ ^●●^ ●●
● ^●^ ●
●
●
●● ^●^
●
● _●_
^●●^ ●● ^●^●_●_^●^ _●_ ●●
●● _●_
●● ●
_●_ ●
●
●^●^ ^●^
●
●●^●^●
●
● ^●^
^●^ ^●^ ^●^●
●
●●● _●●_
●
●_●_●
●
_●_●	●
●●
^●●^ ●●
^●^●^●^● ^●^
●
●
^●^ ●
^●^●_●_●^●^
●
●●
^●^ ●●
●
^●^●
●
●^●^
^●^ ●_●_ ●●^●^
●
●●
^●●●●^ ●
●●
●●
●●
●
●●
●
●
^●●^_●_ ●^●^ _●_ ^●^
●●
●^●●^
●●
●
^●●^●
●
●
●
●● ● _●_ ^●^ ^●^_●_^●^ ●
●
●●●
●●
●●
●
●^●^ _●_
●●^●^
●
●● _●_
^●^●● ^●^
●●
^●^● ●
● ^●^
●
●●
● ^●^	●●
●	●
^●^●^●^
●
●
●●
●
●
●●
●●
●
●
^●^ ●
●
●● ● _●_●●^●^
●● ●● ●^●^ ^●^
● ● ^●^● ●
● ● ●
●
●
● _●_ ●_●_ ●●●
●
●● ●
●● ●_●_ ●
● ● ●
●●●●● ● ●
●_●_●● ●
● ● ●●●●●● ●●●
^●^●^●●^
● ●
●
● ● ●
● ●●
●
●
●●
● ●●
●●● ●
^●^ ^●●^ ^●^ ● ^●^ ^●^ ^●●^
●
● ●
●
● ●●●●●
●●●
^●^ ●
●●
●
●
●^●^ ^●●^
●●
●●
●●
●●
●
●
●●
●
●
●●
●
^●●^_●_ ●●
●
^●^●●
●● ^●^ ^●^ ^●^
●
●
● ● ^●^ ^●^
●
●● ● ^●^
●●
● ^●^
●●
● ^●^ ● ^●^
●●^●^
● ^●^
●
●
^●^ ●
●^●^
●
●●
●●
●●
●
●
●● ●
● ● ^●^
●
●
●
●
^●●^●^●^
●●
●
● ●
●
●
●
●
●
^●^ ●● ^●●^
●
●●
●
●●
_●_●_●_^●^ ^●^ ●
●
● ^●^
●
●●
●●^●^
●●
●
●
●
●●
●
●	●
●
●
^●^●
_●_^●^ ^●^ _●_ _●_● _●●_
●●
●
●
●
^●^●●●
●
●
●●
●	●
●
●
●
●●
●●
● ^●^
●
●
●
● ^●^
●
●
^●^ ^●^●^●^
●●●
●●
●
●
●	^●^
●● ^●^
^●^●● ●
●● ●^●^
●^●●^● ^●^
^●^●^●^
● ●
●● ●
●
●
●●
●
●
●●
● ^●^
●
●
●
●
●
●
●
●
●
●
●●
●
●
●
●
●
●
●
●
●●
●^●^
●	●
●
●
●
●	^●^
●
●
● ●
●
●
●_●_●
●●_●_
●
●●
●
●
●
●●
●
●●^●^
●
●
●●
^●^ ● ^●^
●^●^
●
●
●
● ●
●●
●
^●●^●
●	●
●
● ^●^
●
●
●
●
^●●^●
●●
●
●
● ●
●
●
●●
● ●
●
●
●
●●	^●^ ^●^
● ●●
●
●
● ●
●
●
● ^●^
●	●
● ●
●
●
●
●
●● _●_
●
●
● ●●
●
●
●
^●^●
^●●●^● ●●
●
● ●^●^
●
●
●
●●
●●
● ●
●
●
^●^●^●^ ●
●
●
●● ●
●●
●
● ●●
●●
●
●
●
●
●
_●_ ●
●
^●^●
●
●
●
●
^●^ ●
●
●
●
●
●
●● ●●
^●●●^ ●●
●
● ^●^
●
●
●
●
●
●
●
●
●
^●^ ●
●
●
●
●
●
●
^●●^ ●
●
●
●●^●●^ ^●●^
●●
●
●^●^	_●_ ^●^
●●
● ●
●
●
●●
●
●
●^●^
●
●
● ● ●
●
●
●
● ●^●^● ^●^
●
●
●●
●●
●
^●^ ●●
●
●	●
●	^●^
●
●
●
●
●
●
●
●
●
●
●
●
●●
●
●
●
●●
●
^●^ ^●^●
●
●● _●_
●
●
●
●
●
^●^ _●_●
●
●
●●_●_
^●^●
● ^●^ ●●
●
●
●
_●_ ●
●
●	● _●_
●
●
●
●●
^●^●●
●● ^●^
●^●^
●
●^●^●
^●^ ●●
^●^●
●
●●
●
^●^ ●
●
●
●
●● ^●^
●
●
●
^●^ ●
●
●●
●
●● ●
●
●●
●●
●
●●
●
●
●
●
●●
●●
●●
●
●
●●
●●^●^
●
●
●
●
^●^●●^●^
●●^●^ ●
●
●
●●
●● ●●
●
●
●
●
● ●
●●
●^●^
● ^●^
●
●●
●
●
●
●●●●●
●●
●●
●●
●
● ●
●
●● ^●●●^ ●
●
●
●
●●
●●
^●^ ● ^●●^
●●
●
●
●
●
●●
^●^●● ^●^
●
●
●
●
●
●
^●^●
●●
●
●
●
●
●
●
● ^●^
● ^●●^
● ●
●
●
●
●●
^●^ ● ●
^●^ ^●^ ●●
●
●
●●
● ^●^●●
^●^ ●^●^
●
●● ●
●●	●●
●
●^●●^
●
●
●
●
●● ●
●●
●^●^
●
●
●
_●_●
●
●
●● ●● _●_●●
●
●
●
●
●^●^●
●●●
●
●
●● ●● _●_
●
●
●
●●
●
●
●
●●
^●^●●^●^
●●
●
●
●
●
● ●
●_●_ ● _●_
●^●^
●
●●
_●_ ●
●
●
●
●●●
●●
●●
●
^●^●
●
●
● ●●
^●●^ ●●
●
_●_ ●●
●
●●
●
● ● ●
●●
●●
0.4
●
●
●
●
●● ●●
●
●●
●●_●_
●●^●^
●
_●●_ _●_ ^●^ ●
●● ●●
●
●
●●
●●●●
●●
● ^●^
●
●^●^
●
●
●
^●^ ●_●_
●
●^●^ ^●^
●
●
●
●●
●●
●
^●^●
●
●● ●●●^●^●
●●
●●
●
●●
●
^●^ ●
●●
●
●● ^●^
●
●●
●
●●
●
^●^ ^●^●●
●
●●
●
●●
^●^● ●●
●●_●_
●^●^●
●●
●
●●
● ^●●^●●
●
●
●●^●●^
●●●● _●_ _●_ _●●_
●● ●
●●
● ●●
●
●● ● ●
●
●●
●
●●
●
^●●^●●
●
_●_●
●_●_● _●_
●
● ● ●●
_●_ ●
^●^ ^●^●●
●●
●_●_
●●
●●
●● ●●
_●_ ●● ^●^ ●●
_●_● ^●^ ^●^
●●●●
●
●_●_ ^●●^
●
●●
●● ●●^●^
●●●
●●^●●^
●
_●_●
●
●
●
●
●●
●
●
●●
●●
●●
●●
●
●●
●
●
●
●
_●_ ●● ●●
^●●^●● ●●
●
● _●●_● ^●●^ _●_
^●^ ●●
^●●●^●
●
●●
●
●●
●
●●
●
●●
●●
●
● _●_
●
● ● ●
● ●
●
^●^●
●
●
●●●
●
●
●^●^
●
● ^●^
●
●
●
●
●
●
●
●
●●
●●
●●
●
● ●
●
●
●
●
●
●
●
●
●
●
●
●
●
●
●
●
●
●
●
●
●
^●^● _●_
●
●
●●
●
●
●
●
● _●_
●
●	●
●
● ●
●
●
●
●
●
^●^ ●
●
●
●
_●_ ●●●
●
●
●
●
●
●
●
_●_ ●
●
●
● ●
●
●
●
●
●
●
● ●
●
●
●●
_●_ ●
●
●
●
●
●
●
●
●
●● ●
●^●^
●
●
●
●
●
●
●
●
●
●
●
●
_●_ ●
●
^●^ ●
●^●^
●
●
●●●
●
●
●
●
●
●
●
●
●
●
●
●●
●
●
●
● _●_
●
●
●
●
●
●●
● _●_
●
●
●
●
● ^●^
●
●
●
●
●
●
●
●
●
●
●
●
●
●
●
●
●
●
●
●
0.2
●
●
●
●
^●^ ●
●
●
●
●
●
●
●
●
●
●
●
●
●
●
●
●
●
●
●
●
●
●
●
●
●
●
●
●
●
●
●
●
●
●
●
●
●
●
●
● ●
●
●
●
●
●
●
●
●
●
●
●
●
●●
●
●
●
_●_ ●
●●
●●
●
●
●●
●
●
●
●
_●_● ●● _●_
●●
●●
●
●
●
●
_●_●
●
_●_ ●
●
_●_●
●
●^●^
●●
^●^ ^●^ ●●
● ●
●●
●
●
●●
●
●
●
●●
●●
●●
^●●^ ●
●●
^●^ ●● ●●
●
●●
●●
●●
●●^●^ ^●^
●
● ^●^
●
●●
●●
●●^●^
●
●
_●_ ●●
●
●
●● ^●●^ ●^●^●^●^
●●
●
●
●●
●
●
●●
●
●●
●●
●●
^●^ ^●^ ^●●^●
●
● _●_ ^●^
●
^●^_●_ ● _●_ ^●^ ●
●
●●
●●
^●●^ _●_ _●_●●
●●
^●●^ ●●
●●
●●
●
●	●
_●_ ●
^●^ ^●●^●
●
●
●
_●_^●^●● ^●^_●●_ _●_
●
●●
^●^●^●^
●
^●^●	^●^ ● ^●^
●	●
●
●^●^●●
●
●
●
●
●●
●●●
●
●●
●
^●^ ●
●
^●●●●^●	^●^ ^●^ ^●^
●●
●
●● ^●●^
●
●
^●●^ ●
_●●_^●^ ●
_●_ _●_●● ●^●^
●
_●_●
●●
●
●
^●^ ^●●●●^●● ^●^
●
●^●●^
●
●
●_●●_●●
●
●
●●●^●^
●
●
●
●● _●_●●
●
●
●^●●^
●● _●_
●
^●●^_●_ ● _●_
●
●
●^●^ ^●^ ^●^ ^●^ ●
^●^●●
^●^ ●
●● ●●
●●
●
●●^●^
●
●
●●^●●●^
●
_●_ _●_● ^●●^
● ● ^●●●●^
●
●
●
●●
●● ●_●_
●●
●^●^
●●
● ●
●●
●●●●●
●
^●^●● _●_
●●
●
●
●^●^
_●_ _●_●^●^● ●^●^ ●^●^
●●
● ●
^●●^●^●^●^●^ ^●●^ ^●●^ ^●●●●^
●
● ●
^●●^●
●
●
^●●●^●^●●^●^●^ ^●^ ●^●●^ ^●●●^ ^●^ ^●^
●●●
●●
●● _●_ _●_●
●
●
●
●●^●^ _●_●_●_ ^●^_●_●●
_●_●
●
●●● ●● ^●^
●
^●●●^ ●
●
● ●_●●●_^●^
^●^ ●^●^
^●^ ●
●
● ^●^ ●
^●●●^ ●●
0.0
●
^●^ ●●
● ●
●
●
●●
_●_● _●_ ●● ●●
^●^ ^●^●
● ^●^● ^●^
● ●● ● ●
●●
●
●
●● ●
●
●●
●●
●●_●_^●●^ ● ●
^●^ ●
●
●^●^ ^●^
●●
●^●^ ● ●
●
^●^ ●● ^●^
●
● ^●^● ●
●
●● ^●^_●_ ^●●^ ●_●_ _●●_^●●●^● _●_ ^●^ ^●●^●^●^ ●
●
● ^●^ ^●^ ●
● ●
●	^●^ ●●
^●^ ^●^ ●^●^
^●●●●●●^ ● ^●^ ^●^ ^●●●●^ ^●^
●
^●●^ ●
●●
●
●
●^●^
●
● ^●^ ^●^ ^●^ ^●●^ ^●^
●
●
●
●●
●
_●●_ _●_^●^ ● ^●^ ●●
^●^●_●_ ●●
● ●●^●^ ^●^
● ●
●^●^_●_ ●●^●^●^●^ _●_●^●^
●^●^
●
●	●● ^●^
●
●
_●_● ●_●_●_●_● ●_●_ ^●^_●_● _●_ ●_●_^●^●●
●	●
_●●_^●^ ●●^●^ ● _●_^●●●●^ ^●^_●_●^●^
●●
● ●
●
^●^ ●● ^●●^ ● ^●^
● ●^●●●^
● ●
●
● ● ●
●^●^●
● ●
● ^●^ ● ^●^
●●●
● ●^●^
●
● ●
●
^●^ ^●●●^ ● ^●^ ^●●^ ^●●^ ^●^	^●^ ^●^ ●^●^ ● ^●^
● ^●^● ●^●^● ^●●●●^ ●^●^ ^●●^ ^●^ ^●^ ^●^
^●^ ^●^ ^●^ ● ^●^ ●
● ^●^
●^●^ ^●^
●^●^● ^●●^● ^●^ ●●
●
●	^●^ ●●
●
●● _●●_●^●^● _●_^●^● ^●^
_●_ ● _●_^●^
^●●●●●^ ^●●●^ ^●^ ^●^●^●^ ● ^●^
^●^●● ^●^●	●
●^●^ ^●●^ ^●^
●
●
●
●
^●^ ^●●^ ● ^●^ ^●●^ ^●●●^● ^●^ ^●^ ^●●●●^ ^●●^ ^●●^●^●●^ ^●●●●●^ ^●^ ^●^
●
●
_●_ _●_● ^●●^ ● ^●^_●_●●_●_● ^●^
^●^●
^●^●
^●^ ^●^ ●
●
●
● ^●^ ^●^	● ^●●^ ^●^●^●●^ ^●^
●
^●^ ^●^ ^●^●^●^● ^●●^	^●^ ^●^
●
● ●^●^●^●^ ^●●^
●
●●●	●
● ● ^●^ ●● ^●●^
●
^●^ ●^●^
^●^	●
●●
●	●^●^ ●●●● ● ●●● ● ● ●	● ●
●●● ● ● ● ●
●	● ● ● ●●● ●● ●●● ●● ● ● ● ● ●●● ●●●	●
● ● ●● ●●
●●	●
● ● ●
●● ●● ● ●●●●● ● ●●● ● ●
● ● ●● ●●●●●● ● ● ● ●●●●●
● ●● ●● ● ●●● ●● ● ●
●	●	●●● ●●● ● ● ● ●
● ● ● ● ●● ● ● ●● ●●●
● ●●	●● ● ● ●●● ●●● ● ● ●● ●● ● ● ●●	●●●● ● ●●
0.0e+00
5.0e+07
1.0e+08
1.5e+08
2.0e+08
2.5e+08
Genomic position
