## Supplemental Fig. S10. Overview of all known structural variants in ABCA4 for "Resolving the dark matter of *ABCA4* for 1,054 Stargardt disease probands through integrated genomics and transcriptomics"

### Slide 1
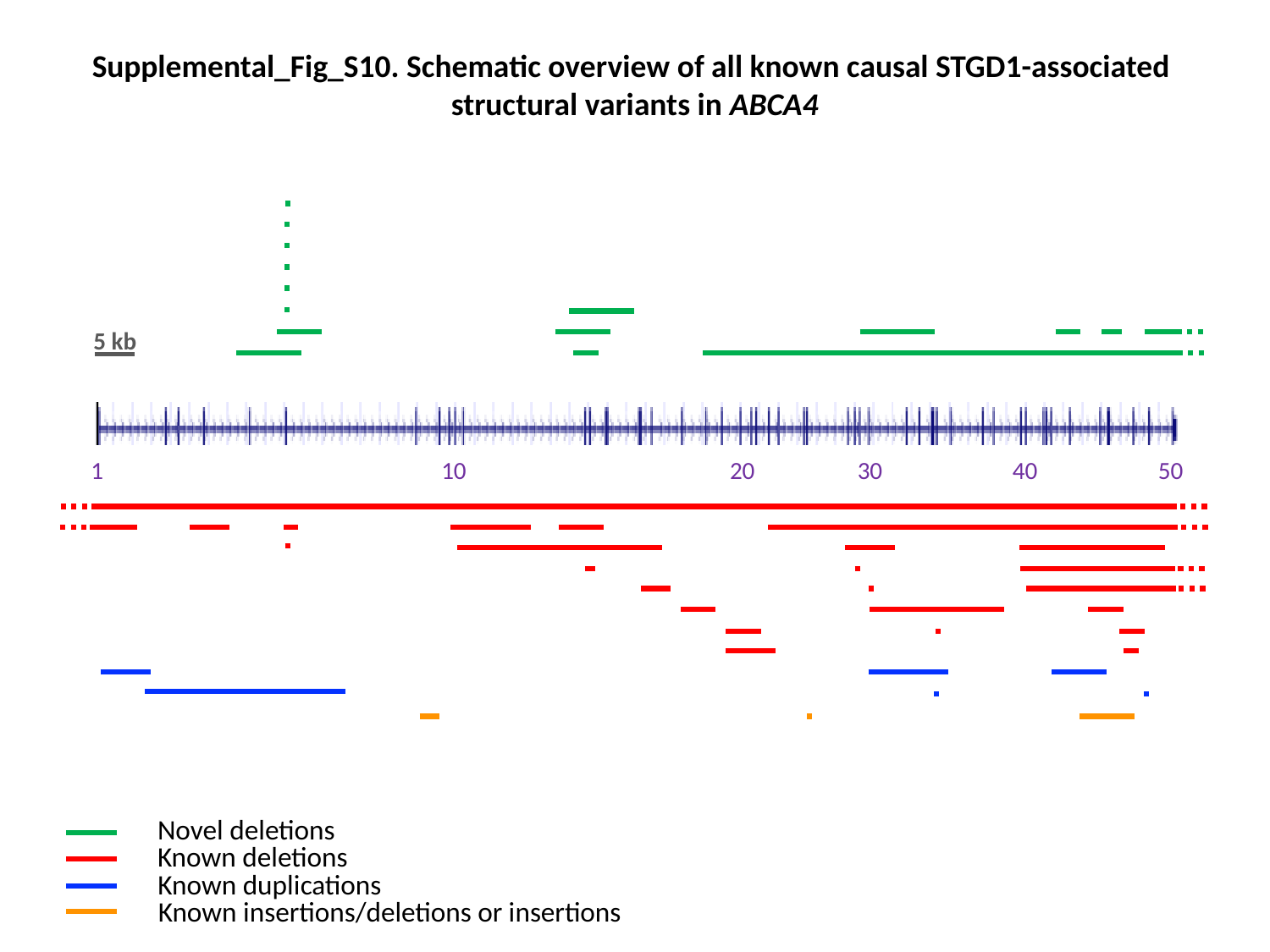

Supplemental_Fig_S10. Schematic overview of all known causal STGD1-associated
structural variants in ABCA4
5 kb
10
50
 30
1
20
40
Novel deletions
Known deletions
Known duplications
Known insertions/deletions or insertions
