## Supplemental Table S5. Copy number variants in control STGD1 cases for "Resolving the dark matter of *ABCA4* for 1,054 Stargardt disease probands through integrated genomics and transcriptomics"

**Supplemental_Table_S5. Details of structural variants in controls**

| DNA variant | Protein variant | Consequences | Reference |
| --- | --- | --- | --- |
| c.(?_-1)_(66+1_67-1)del | p.(?) | Exon 1 deletion | This study |
| c.66+520_67-389dup | p.(?) | 7 kb duplication in intron 1 | Bauwens et al, 2019 |
| c.67-975_769-4582dup{insA} | p.(Ile23_Val256dup) | Exons 2 to 6 duplication | Bauwens et al, 2019 |
| c.442+1195_ 571-1400del | p.(Gly148Valfs*89) | Exon 5 deletion | This study |
| c.699_ 768+341del | p.(Gln234Phefs*5) | Partial exon 6 deletion | This study |
| c.1239+291_1555-5574del | p.(Ala414_Glu518del) | Exons 10 to 11 deletion | Bauwens et al, 2019 |
| c.2918+775_3328+640del^#^ | p.(Ser974Glnfs*64) | Exons 20 to 22 deletion | Maugeri et al, 1999* Bauwens et al, 2019 |
| c.3863-1241_4539+290del | p.(Gly1288Glufs*41) | Exons 27 to 30 deletion | This study |
| c.4353-13_ 5091dup | p.(Val1698Cysfs*4) | Exons 30 to 36 duplication | This study |
| c.4352+282_ 5461-451del^#^ | p.(Glu1452Argfs*9) | Exons 30 to 38 deletion | This study |
| c.5585-166_*1254del | p.(Gly1862*) | Exons 40 to 50 deletion | Bauwens et al, 2019 |
| c.(5584+1_5585-1)_(*1_?)del | p.(Gly1862*) | Exons 40 to 50 deletion | This study |
| c. 6006-30_6371dup | p.(Leu2125_Asp2273delins76) | Exons 44 to 46 duplication | This study |

Thirteen different structural variants in 15 DNA samples were used as controls. All variations are described using HGVS (Human Genome Variation Society) nomenclature. Data are provided for DNA variant (human genome version 19; hg19), as well as protein effect when possible. DNA variant positions indicated are the first and the last deleted nucleotides. Functional consequences of deletions are reported in indicating the exons included in the deletion or duplication. ^#^ Each deletion was identified in two patients, * was first described by Maugeri et al, 1999, but exact breakpoints were shown in Bauwens et al, 2019.
