## Supplemental Table S14. In silico breakpoints analysis and underlying mechanisms for copy number variant formation for "Resolving the dark matter of *ABCA4* for 1,054 Stargardt disease probands through integrated genomics and transcriptomics"

**Supplemental Table S14. *In silico* breakpoints analysis and underlying mechanisms for structural variants formation**

|  |  |  |  |  | **Proximal breakpoint region** | | **Distal breakpoint region** | |  |
| --- | --- | --- | --- | --- | --- | --- | --- | --- | --- |
| **Deletions and breakpoints** | **Genomic position (hg19)** | **Deletion size (bp)** | **Micro-homology (bp)** | **Inserted sequence at breakpoint** | **Repetitive element** | **Number of non-B DNA conformation prediction motifs** | **Repetitive element** | **Number of non-B DNA conformation prediction motifs** | **Proposed mechanisms** |
| **1_BP1-BP2** | 94569917-94562911 | 7,007 | 4 | - | [Copia-30_FV-I](https://www.girinst.org/protected/repbase_extract.php?access=Copia-30_FV-I&format=EMBL) | 2 | - | - | Replicative/NHEJ |
| **2_BP1-BP2** | 94565348-94561288 | 4,061 | 6 | - | MER53 | - | - | 3 | Replicative/NHEJ |
| **3_BP1-BP2** | 94564419-94564009 | 411 | 2 | - | [L1-72_DR](https://www.girinst.org/protected/repbase_extract.php?access=L1-72_DR&format=EMBL) | 1 | MIRb | 1 | Replicative/NHEJ |
| **4_BP1-BP2** | 94529906-94527518 | 2,389 | 2 | AGC | L2c | - | - | 2 | Replicative/NHEJ |
| **5_BP1-BP2** | 94532364-94528694 | 3,671 | 2 | - | MER3 | 2 | - | 0 | Replicative/NHEJ |
| **5_BP3-BP4** | 94528416-94526398 | 2,019 | 2 | - | - | 1 | L2a | - | Replicative/NHEJ |
| **6_BP1-BP2** | 94531301-94526086 | 5,216 | 2 | - | - | - | - | 3 | Replicative/NHEJ |
| **6_BP3-BP4** | 94525863-94522479 | 3,385 | 1 | ATGAATG | Copia-14_Mac-I | 1 | MamTip2b | 3 | Replicative/NHEJ |
| **7_BP1-?** | 94514513-94458793 | >55,721 | n.a. | n.a. | n.a. | n.a. | n.a. | n.a. | n.a. |
| **8_BP1-BP2** | 94496279-94487503 | 1,225 | 4 | GCTTTTT | - | 1 | - | 2 | Replicative/NHEJ |
| **9_BP1-BP2** | 94470240-94472532 | 2,293 | 1 | TTTTAACAGTGTT | - | 3 | L1MC4a | 0 | Replicative/NHEJ |
| **10_BP1-BP2** | 94467351-94463600 | 3,752 | 3 | - | - | 3 | - | 1 | Replicative/NHEJ |
| **11_BP1-?** | 94461751-94458793 | >2,959 | n.a. | n.a. | n.a. | n.a. | n.a. | n.a. | n.a. |

Deletions 7 and 11 have been excluded from this *in silico* analysis as their exact breakpoints were not known. Deletions 5 and 6, each comprised of two separate rearrangements and were considered as independent events. They therefore were named as following: deletion 5 (BP1-BP2) and deletion 5 (BP3-BP4), as well as deletion 6 (BP1-BP2) and deletion 6 (BP3-BP4). When two different events appeared in one deletion, they were noted as BP1-BP2 and BP3-BP4.

BP, breakpoint; NHEJ, non-homologous end joining; n.a., not applicable. Replicative stands for replicative-based mechanisms and includes FoSTes (fork stalling and template switching), MMBIR (microhomology-mediated break-induced replication), SRS (serial replication slippage) and BISRS (break-induced serial replication slippage).

**Classification of the identified repetitive elements:**

7 non-LTR retrotransposons: 1 SINE (MIRb) and 4 LINEs (L1-72_DR, L2c, L2a and LIMCAa)

3 DNA transposons from the *hAT* superfamily (MER53, MER3 and MamTip2b)

2 retrotransposons from the LTR superfamily (Copia-30_FV-I and Copia-14_Mac-I) (Kojima, 2018).
