## Supplemental Table S15. Sequences of non-B DNA conformations for "Resolving the dark matter of *ABCA4* for 1,054 Stargardt disease probands through integrated genomics and transcriptomics"

| **Deletion breakpoint** | **Inverted Repeats and cruciform motifs** | **Direct repeat and splipped motifs** | **Mirror repeats and triplex motifs** | **Short tandem repeats** | **Z-DNA motifs** | **Oligo(G)n tract** |
| --- | --- | --- | --- | --- | --- | --- |
| SV1 proximal BP | - | - | - | - | - | **GG**AGA**GG**AAGA**GG**ATG**GG, GG**AATT**GG**GACCA**GG**AAAGAAT**GG** |
| SV2 distal BP | TCCATTTGACAAATGGA, GTTTTTAGAAAAAC | - | - | AAAAAAAAAAAAAAAA | - | - |
| SV3 proximal BP | - | - | - | - | - | **GGG**TGCCCT**GGG**AGA**GGG**AGCACA**GGG** |
| SV3 distal BP | - | - | - | - | - | **GG**AA**GG**A**GG**A**GG** |
| SV4 distal BP | - | - | - | - | - | **GG**ATTT**GG**G**GG**GT**GG**, **GGG**CCG**GGG**TGGA**GGG**TT**GGG** |
| SV5 BP1-BP2 proximal BP | CTGCTCTGGAGCAG, AGAAGATCTTCT | - | - | - | - | - |
| SV5 BP3-BP4  distal BP | - | - | - | - | - | **GG**GGT**GG**TAGAGAGCT**GG**TCCA**GG** |
| SV6 P1-BP2  distal BP | CTGCTAATAGCAG | - | - | - | - | **GG**TCAC**GG**AATG**GG**ACTG**GG, GG**ACCCCTG**GG**CAGGAAGT**GG**GAGATG**GG** |
| SV6 BP3-BP4  proximal BP | - | - | - | - | - | **GG**A**GG**AAA**GGGG** |
| SV6 BP3-BP4  distal BP | - | - | - | - | - | **GG**GAAAAT**GG**CATTCCTA**GG**ATTAAA**GG**, **GG**CT**GG**TG**GG**AGAGAGCCCTTTA**GG**, **GG**CGTCC**GG**CCAA**GG**GCA**GG** |
| SV8 proximal BP | - | - | - | - | - | **GG**GAA**GG**CCGAAT**GG**GGAAA**GG** |
| SV8 distal BP | - | - | - | - | - | **GGC**GTGAAGTTCGT**GG**CCCT**GG**TCTGA**GG, GG**TAT**GG**A**GG**AATTTCCATT**GG** |
| SV9 proximal BP | TTTCCTGGCTGCAGGAAA | - | - | - | - | **GG**CGTGTGCCCTT**GG**ATTCT**GG**AG**GG, GG**TCCCCTAT**GG**GCA**GG**ACAGAGA**GG** |
| SV10 proximal BP | AAACTGCAGCAGTTT | - | - | - | - | **GGT**GCTGCT**GG**TAACTGC**GG**GCTTG**GG, GGG**TTGG**GG**AAATA**GG**TTCTG**GG** |
| SV10 distal BP | - | - | - | - | - | **GGGG**AACTTCCCA**GG**CAGTGTGCAGA**GG** |

Sequences of 150 bp surrounding each breakpoint of the normal wild-type proximal and distal sequences have been analyzed by QGRS and non-B DB softwares. Inverted motifs are indicated in red and oligo(G)n tracts are underlined and bold.
