## Supplemental Table S19. STGD1 collaborators and cases for "Resolving the dark matter of *ABCA4* for 1,054 Stargardt disease probands through integrated genomics and transcriptomics"

**Supplemental_Table_S19. Collaborators and Stargardt disease and Stargardt disease like cases details**

| **Collaborator; city** | **Total # of cases** | **# of unscreened probands** | **# of prescreened probands** | **Prescreening method** |
| --- | --- | --- | --- | --- |
| AlTalbishi, A.; Jerusalem | 37 | 11 | 26 | TSS |
| Ayuso, C.; Madrid | 80 | 0 | 80 | MIPs, Microarray, WES |
| Banfi, S.; Naples | 40 | 0 | 40 | Sanger, Microarray, NGS |
| Ben-Yosef, T; Haifa | 27 | 0 | 27 | Sanger |
| Dhaenens, C-M., Lille | 184 | 0 | 184 | dHPLC, HRM, Sanger, NGS |
| Fakin, A.; Ljubljana | 14 | 0 | 14 | Microarray |
| Farrar, J.; Dublin | 36 | 0 | 36 | Microarray, NGS |
| Ferraz Sallum, J.M.; Sao Paulo | 9 | 0 | 9 | NGS |
| Fujinami, K.; Tokyo | 28 | 0 | 28 | WES |
| Gorin, M.B.; Los Angeles | 26 | 0 | 26 | Sanger, Microarray, NGS |
| Kamakari, S.; Athene | 6 | 0 | 6 | Sanger, WES, MLPA |
| Liskova, P.; Prague | 52 | 50 | 2 | Microarray |
| MacDonald, I.; Alberta | 3 | 0 | 3 | NGS, WES |
| Oldak, M.; Warsaw | 22 | 13 | 9 | NGS |
| Podhajcer, O.; Buenos Aires | 27 | 0 | 27 | NGS |
| de Roach, J. & Lamey, T.; Perth | 69 | 27 | 42 | Microarray, NGS |
| Roberts, L.; Cape Town | 149 | 21 | 128 | TSS, SSCP, Microarray, NGS |
| Sharon, D.; Jerusalem | 22 | 3 | 19 | TSS, Sanger |
| Tracewska, A.; Wroclaw | 4 | 0 | 4 | smMIPs, Sanger |
| Vincent, A.; Auckland | 28 | 0 | 28 | Microarray, NGS |
| Weber, B., Regensburg | 148 | 94 | 54 | Microarray, NGS |
| Hoyng, C.B.; Nijmegen | 33 | 0 | 33 | Sanger, smMIPs, NGS, WES, MLPA |
| Boon, C.B.; Amsterdam | 1 | 0 | 1 | NGS |
| Van den Born, L.I; Rotterdam | 8 | 2 | 6 | Sanger, NGS, WES, MLPA |
| Klaver, C.CW.; Nijmegen | 1 | 0 | 1 | NGS |
| **Total numbers** | **1054** | **221** | **833** |  |

#, Number; n.a., not applicable; dHPLC, denaturing high performance liquid chromatography; HRM, high resolution melt; Microarray, *ABCA4* targeted mutation detection; MIPs, molecular inversion probe based-sequencing; MLPA, multiplex ligation-dependent probe amplification; NGS, next-generation sequencing; Sanger, *ABCA4* exonic sequencing; smMIPs, single molecule molecular inversion probe based-sequencing; SSCP, single strand conformation polymorphism; TSS, *ABCA4* targeted mutation Sanger sequencing; WES, whole exome sequencing
